# A bifidobacterial enzyme orchestrates ecology and function of infant gut bacterial community

**DOI:** 10.64898/2026.04.28.718440

**Authors:** Yangwei Shan, Nicholas Pucci, Coen Berns, Rick Hoogendijk, Myrthe Beijnvoort, Shijia Li, Alejandro Sánchez-Cano, Gertjan Kramer, Wei Du, Daniel R. Mende, Aalt Dirk Jan van Dijk, Meike Wortel, Jianbo Zhang

## Abstract

Human milk oligosaccharides (HMOs) are abundant and structurally diverse glycans that shape the development of infant gut microbiota. Yet, how individual HMOs and bacterial genes drive the community assembly remain elusive. Here, we reconstructed an eight-member infant Bacterial Community (iBaCo) from representing dominant taxa in human infant feces. When individual HMOs were the sole carbohydrate source, they showed deterministic effects on the iBaCo composition and metabolic output. Notably, the tetramer HMO lacto-N-tetraose (LNT), in spite of its identical monomer composition as lacto-N-neotetraose (LNnT), showed a strong effect on maintaining *Bifidobacterium breve* abundance in iBaCo, whereas LNnT did not. Monoculture growth profiling, proteomics, enzymatic kinetic assay, and molecular docking revealed that *β*-galactosidase D4BMY8 and the relevant downstream pathways are induced by LNT and that D4BMY8 has substrate preference on LNT over LNnT, enabling a faster growth of *Bi. breve* and accumulation of acetate and lactate in LNT compared to LNnT. Metabolic flux analysis indicated that the substrate-preference of *β*-galactosidase D4BMY8 drives the skewed energy cost toward lactate/acetate metabolic output. Finally, the D4BMY8-encoding gene *lacZ5* is widely spread in all isolated *Bi. breve* genomes, but divergently distributed in infant metagenome-assembled *Bi. breve* genomes. Together, we demonstrated that a single enzyme-substrate interaction could orchestrate the composition and metabolic function of an infant bacterial community, which may contribute to the assembly of dynamic infant gut microbiota. Our integrative approach provides a mechanistic framework for understanding the interaction between diet, microbial community, and infant gut health.

## Introduction

Human milk provides nutritional, immunological, and developmental support for newborns^1–3^. Carbohydrates constitute approximately 7.1% of human milk, of which lactose accounting for ∼80% and human milk oligosaccharides (HMOs) ∼20%.^1^ Compared to lactose, HMOs exhibit greater structural complexity due to their diverse monosaccharide compositions and glycosidic linkages. Despite the complexity, most HMOs share five monosaccharides: glucose (Glc), galactose (Gal), α-L-fucose (Fuc), α-N-acetylglucosamine (GlcNAc), and α-D-N-acetylneuraminic acid (Neu5Ac).^4^ Based on their glycosylation patterns, HMOs are categorized into three major groups: fucosylated (35–50%), sialylated (12–14%), and non-fucosylated neutral HMOs (42–55%).^5^ These HMOs can not be digested by human pancreatic enzymes,^6,7^ allowing them to pass through the upper gastrointestinal tract and reaching the colon, where they can be utilized by infant colonic microbiota.^8,9^

The establishment and assembly of infant colonic microbiota starts from microbial colonization immediately after birth, and is associated with various factors including delivery modes and feeding practices^10,11^. Among early colonizers, *Bifidobacterium* species are abundant in the fecal microbiota of breastfed infants, often as dominant bacteria^12^, while Enterobacterales, Bacteroidales, or Clostridiales can be dominant bacteria in other infants^13^. Importantly, reduced *Bifidobacterium* levels during infancy have been linked to immune dysregulation and increased risk of allergic and inflammatory disorders.^14,15^ Their metabolic activities support gut barrier integrity and immune regulation, while their competitive niche occupation helps suppress pathogen overgrowth^16,17^, underscoring the importance of sustaining their abundance during infancy, a critical developmental window of human life.

Although HMOs are known to shape infant gut microbiota, we still lack mechanistic insight into how individual HMO structures, and the specific microbial genes they engage, determine community assembly and function in a defined system. Synthetic microbial communities (SynComs) have emerged as essential models to capture the microbiota complexity and interrogate microbiota assembly principles in a defined setting.^18^ For instance, Ioannou et al.^19^ established a 13-member Bifidobacterium-dominant SynCom (BIG-Syc) derived from breastfed neonates and revealed extensive resource sharing among the members.^19^ Among Bifidobacterium species, priority effects were observed during the assembly using SynComs consisting of only bifidobacteria^20^. Theoretical and experimental work suggest that competitive interaction often predominant over cooperative interaction in shaping microbial communities^21,22^, highlighting competition is another important driver of the community assembly.

Here, we asked whether a single HMO-enzyme interaction can steer the assembly and metabolic output of an infant gut SynCom, thereby providing a mechanistic bridge between molecular traits and ecological behavior. To this end, we constructed a SynCom termed infant Bacterial Community (iBaCo) comprising eight infant gut species: four from Bifidobacteriaceae, two Lachnospiraceae, one from Bacteroidaceae and Coriobacteriaceae, respectively.^13,23,24^ We mapped the iBaCo response to individual HMOs and identified a specific increase of *Bi. breve* and *Bi. pseudocatenulatum* abundance when grown on the HMO lacto-N-tetraose (LNT). Paired monoculture of each species-HMO combination pointed to diverse strategies of HMO utilization. Proteomic profiling, biochemical analysis, and recombinant protein experiments confirmed that substrate preference of LNT over its isomer lacto-N-neotetraose (LNnT) by *Bi. breve* is primarily driven by a specific *β*-galactosidase, orchestrating iBaCo composition and function. These results demonstrated that a single gene/enzyme can be a major determinant of microbial community composition and function, highlighting the value of mechanistic approach for microbial ecology and gut microbiome research.

## Material and methods

### Material, strains, reagents and software

All strains, plasmids, primers, reagents, and software used in this study are listed in Supplementary Table 1-5.

### Bacterial cultivation

*Bifidobacterium breve* JCM1192/DSM20213, *Bifidobacterium breve* UCC2003 wild-type*, Bifidobacterium longum subsp. infantis, Bifidobacterium bifidum, Bifidobacterium pseudocatenulatum, Bacteroides fragilis, Blautia luti, Anaerostipes caccae and Collinsella aerofaciens* were cultivated in Yeast Casitone Fatty Acids (YCFA) broth medium under anaerobic conditions (10 % H_2_, 10 % CO_2_, 80% N_2_, Whitley A35 anaerobic workstation). *Escherichia coli* DH5α and BL21 strains were cultured in LB broth or agar, supplemented with 50 mg/L kanamycin. *Bifidobacterium breve* UCC 2003 mutants were cultured in YCFA broth or agar, supplemented with 10 mg/L tetracycline. For growth-curve measurement, human milk oligosaccharides (HMOs) were dissolved in YCFA without carbohydrate (YCFA-no sugar) medium at a final concentration of 6 g/L to yield YCFA-HMO, then filtered via a 0.22-μm membrane. For monoculture, bacteria were seeded in 96-well flat bottom plates at inoculation rate of 1% (v/v), with three technical replicates for each condition, negative control (no bacteria), and a positive control (YCFA-GMC with glucose maltose and cellobiose). The plates were sealed with sealing film (Greiner Bio-One, 676070), and the optical density was measured every 15 minutes for 72 hours using the microplate reader Absorbance-96 (BYONOY, Germany). After 72 h, cell pellets and supernatants were separately harvested by centrifuge at 5,000 *g* for 5 min. This was filtered through a 0.2 μm PVDF membrane (Agilent, 203982-100) on a new round-bottom 96-well plate. The 96-well filter was centrifuged for 3 minutes at 1,000 g (4 °C). The filtered samples were stored in HPLC vials (BGB Analytik, PPSV0903K) at - 20 °C until further analyses. Cell pellets were stored at −20 °C for strain identity confirmation.

### PCR and gel electrophoresis

PCR was used to amplify multiple target genes, including 16S rRNA gene for strain identification and sequencing, as well as *β*-galactosidase genes fragments for cloning. Reactions were assembled using gene-specific primers (Supplementary Table 4), DNA template (either bacterial colony or purified genomic DNA, depending on the application), 2× Rapid Taq Master Mix (or 2× Phusion Master Mix), and nuclease-free water in a final volume of 50 μL. Primer concentrations, template input, and cycling parameters were optimized for each target and are described in the corresponding sections below. PCR products were verified by agarose gel electrophoresis. A 1.5% agarose gel was prepared in 1× TAE buffer (50× stock: 242 g Tris base, 57.1 mL glacial acetic acid, 100 mL 0.5 M EDTA, pH 8.5) and supplemented with 1,000× Midori Green. 5 μL PCR product or 1 kb Plus DNA Ladder were loaded and electrophoresed at 120 V for 30 min. DNA bands were visualized and captured in Gel Imaging System.

### Strain identity confirmation

For bacterial identification, colony PCR was performed directly from single colonies. Briefly, a small amount of bacterial biomass was suspended in 20 μL MilliQ water. 0.5 μL of cell suspension was used as template in a 50 μL PCR reaction containing 25 μL 2× Rapid Taq Master Mix (Vazyme, P222), 0.5 μM of each universal 16S rRNA primers (8f and 1492r, Supplementary Table 4), and 20.5 μL nuclease-free water. Cycling conditions were: initial denaturation at 95 °C for 3 min; 30 cycles of 95 °C for 15 s, 55 °C for 15 s, and 72 °C for 30 s; followed by a final extension at 72 °C for 5 min. PCR products were screened by gel electrophoresis. After PCR, amplicons were purified by using magnetic beads (abm, G951). PCR products were mixed with beads at a 1:0.9 ratio in a 96-well plate and placed on a magnetic rack to settle the beads. After condensation, the supernatant was removed and the beads were washed twice with 70% ethanol. The plate was then removed from the magnetic rack, and DNA was eluted from the beads with Milli-Q water. After a final magnetic separation, the eluate was collected as the purified PCR product. Afterwards, the purified amplicons were quantified by Nanodrop, sent for Sanger sequencing (MAD core facility at University of Amsterdam). The returned DNA sequences were Blasted to confirm the identity of the bacterial species.

#### iBaCo construction and cultivation

iBaCo consisted of the 8 species (*Bi breve* JCM1192/DSM20213*, Bi. infantis, Bi. bifidum, Bi. pseudocatenulatum, Ba. fragilis, Bl. luti, A. caccae and C. aerofaciens*). To construct iBaCo, all 8 species were subcultured overnight twice. The OD_600_ values were then measured and normalized for each species. The inoculating OD_600_ value was set 0.0025 per mL for each species. After mixing the 8 species as an inoculum, the inoculum was inoculated into a Hungate tube containing 10 mL 10% YCFA media in DPBS and then cultured at 37 °C. The starting OD_600_ value was 0.02. All procedures were conducted in the anaerobic chamber. After 24 h of culturing, 1 mL liquid cultures were transferred to ep tubes and centrifuged at 10,000 *g* for 1.5 min. The resulting cell pellets and supernatants were collected separately for further analysis.

#### Short chain fatty acid quantification

High-performance liquid chromatography (HPLC) was used to analyze bacterial metabolites (acetate, propionate, butyrate, lactate, succinate, and formate). Samples were prepared by 1 : 1 (v/v) mix with 1 mM H_2_SO_4_, and filtered through a 0.22 μ m filter. Standards were prepared with a set of concentrations within 0-50 mM in 10 mL MilliQ water as stock solutions. The HPLC program and analysis method were described previously^25^. In brief, the HPLC was equipped with a refractive index detector (RID), and the temperature of the detector cell was maintained as 40 □. Isocratic flow of mobile phase (5 mM H_2_SO_4_) was used at 0.5 mL/min and the column was maintained at 55 □. The quantification limit of butyrate is 1.2 mM (**Supplementary Figure 4**).

#### Thin layer chromatography (TLC) analysis

TLC was used to detect the HMO degradation and monosaccharides. Briefly, the spent media were centrifuge-filtered at 5,000 *g* for 5 min, and 2 μL of filtered spent media were sampled via a capillary tube, then spotted on the TLC plate (Sigma-Aldrich, 1.05549). The plate was air dried and transferred into the mobile phase (1-butanol: acetic acid: water = 2: 1: 1 v/v). Once the mobile phase reached the top of the TLC plate, it was taken out from the mobile phase, air-dried in chemical hood. To visualize the HMO and monosaccharides, the whole plate was dipped into sulfuric acid: ethanol (5: 95 v/v) solution and then heated with a blow-dryer until clear bands were visible.

#### Nanopore sequencing

The cell pellets of iBaCo were used to extract the genomic DNA using PureLink™ Genomic DNA Mini Kit. To obtain the clean 16s rRNA gene amplicons, genomic DNA was amplified with universal 16s primers (8f and 1492r, Supplementary Table 4), followed by purification with DNA Purification magnetic beads (abm, G951). Briefly, 2 μL purified genomic DNA was used as template in a 50 μL PCR reaction containing 25 μL 2× Rapid Taq Master Mix (Vazyme, P223), 0.5 μM of each universal 16S rRNA primers. Cycling conditions were: initial denaturation at 95 °C for 2 min; 35 cycles of 98 °C for 10 s, 52 °C for 30 s, and 68 °C for 30 s; followed by a final extension at 68 °C for 5 min. The purified PCR products were then quantified using a NanoDrop spectrophotometer, 400 ng of DNA from each sample was used for library preparation. Sequencing libraries were prepared according to the gDNA ligation sequencing protocol using Native Barcoding Kit 24 V1 (SQK-NBD114.24). During library preparation, samples were labeled with unique barcodes and pooled into a single library. The pooled library was quantified using a Qubit fluorometer according to the manufacturer’s instructions, then diluted to 20 fmol in 12 µL of fresh Milli-Q water. The final library was prepared and loaded onto a Flongle Flow Cell following the protocol provided by Oxford Nanopore Technologies. Data analysis was performed on the EPI2ME agent (Oxford Nanopore Technologies), and 16s workflow was used for the iBaCo composition analysis. The reads of every iBaCo species were recorded and normalized by their corresponding 16S rRNA gene copy numbers.

#### Analysis of microbial growth kinetics

The growth kinetics of eight gut microbial strains were determined based on time-optical density (time-*OD*) curves obtained from the monoculture experiments. For each carbohydrate source, the maximum optical density (*OD_max_*) was first extracted. Growth curve fitting and parameter estimation were then performed using the R package *growthrates*, with the *fit_easylinear* function employed to identify the exponential growth phase and calculate key parameters, including the maximum specific growth rate (*µ_max_*) and lag phase (*λ*) (**Supplementary Figure 7**). Under ideal conditions, microbial cell numbers increase exponentially during cultivation, and the changes in *OD* can be expressed as a function of its natural logarithm rather than *OD* itself. Thus, during the exponential phase, we assumed a linear relationship between the natural logarithm of cell density and time, expressed as:

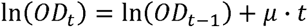

where *OD* represents the cell density and *µ* denotes the specific growth rate (*h*^-1^). In practice, *OD* values were log-transformed to linearize the exponential phase. A sliding time window of 12 h was applied throughout the 72 h cultivation period, within which local linear regressions were fitted between ln(*OD_t_*) and time *t*. The regression model with the highest coefficient of determination (R^2^) among all windows was selected as the optimal representation of the exponential phase, with its slope taken as the maximum specific growth rate (*µ_max_*). Consequently, the tangent line to the logarithmic growth curve at the point of maximal growth rate can be expressed as:

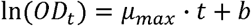

where *b* is the intercept of the regression line. The lag phase (λ) was determined by extrapolating this regression line back to its intersection with the initial cell density baseline ln(*OD*_0_), with the corresponding coordinate on the time axis represent the lag duration:

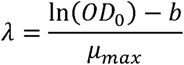

To ensure biological plausibility, calculated lag times were constrained between 0 and 7 h (i.e.,*λ*<0 was set to 0 h, and *λ*> 72 was set to 72 h).

#### High-performance anion exchange chromatography (HPAEC)

HPAEC was used to quantify HMOs and monosaccharides. The spent media or HMO hydrolysate were centrifuged at maximum speed at 4 °C for 10 min and diluted 100-fold with fresh Milli-Q water, then 1 mL sample was transferred to Verex vials (Phenomenex, AR0-39P0-13). Samples were then injected onto an HPAEC system (Dionex™ ICS-6000, Thermo Fisher Scientific) equipped with pulsed amperometric detection using a gold electrode and a carbohydrate quad waveform. For the analytical program, 100 mM NaOH was prepared as eluent A, and 100 mM NaOH with 1 M NaAc was prepared as eluent B. The flow rate was maintained at 0.8 mL/min with the following elution profile: 100% A for the first 10 min; from 10 to 28 min, eluent B was gradually increased to 20%; eluent B was then increased to 100% at 31 min and maintained until 35 min. Subsequently, eluent B was decreased to 0% at 36 min, and 100% A was maintained until 41 min. Chromatographic data were analyzed using Chromeleon (version 7.2.10 ES MUj, Thermo Fisher Scientific).

#### Bacterial cultivation for proteomics analysis

*Bi. breve* JCM1192/DSM20213 was cultured in YCFA-GMC, YCFA-LNT, or YCFA-LNnT, respectively until the OD_600_ was above 1 (late exponential to early stationary phase). Then, 1 mL bacterial culture was pelleted by centrifugation (10,000 rpm, 4 °C, 5 min, Eppendorf Centrifuge 5415R), then washed with DBPS. After that, 300 μL lysis buffer (25× cOmplete inhibitor cocktail, 1M Ambic, and 20% SDS) was added to resuspend each cell pellet and then incubated for 10 min at room temperature on a rotator. Afterwards, lysates were sonicated for 6 rounds (15 seconds sonication and 15 seconds pause for each round). Following centrifugation (10,000 rpm, 4 □ for 90 sec), the supernatants were collected to measure the protein concentration using BCA assay according to the manufacturer’s instructions. After that, the samples were diluted to a concentration of 0.1 mg/mL at a volume of 250 μL. Samples were then reduced and alkylated in one step by incubation with tris-(2-carboxyethyl) phosphine and chloroacetamide (Sigma Aldrich) at an end concentration of 10 mM and 30 mM respectively for 30 minutes at 70 °C^26^. Subsequently, samples were prepared for mass spectrometry analysis using the single-pot, solid-phase-enhanced sample preparation (SP3) protocol with optimization^27^. Briefly, no detergents were added to the samples to enable optimal precipitation of soluble proteins, and the precipitation time was extended to 30 minutes at room temperature. Subsequently, the washed beads were air-dried and resuspended in 100 mM ammonium bicarbonate (Sigma Aldrich) after which trypsin (Sequencing Grade Modified, Promega) was added at a protease-to-protein ratio of 1:50 (w/w) at 37 °C. After overnight digestion, formic acid was added to a final concentration of 1%, adjusting the solution to an approximate pH of 2. The samples were then placed on a magnetic separator device and the peptides were recovered for LC-MS-based proteomic analysis.

#### LC-MS-based proteomic analysis

Samples were analyzed by reversed phase chromatography using an Ultimate 3000 RSLCnano UHPLC system (Thermo Scientific, Germeringen, Germany) and peptides were separated using a 75 μm × 250 mm analytical column (C18, 1.6 μm particle size, Aurora, Ionopticks, Australia), maintained at 50 □ and operated at a flow rate of 400 nL/min with 3% solvent B for 3 minutes (solvent A: 0.1% formic acid in water, solvent B: 0.1% formic acid in acetonitrile, ULCMS-grade, Biosolve). Next, a multi-stage gradient was applied (17% solvent B at 21 minutes, 25% solvent B at 29 minutes, 34% solvent B at 32 minutes, 99% solvent B at 33 minutes, kept at 99% solvent B till 40 minutes). At 40.1 minutes, the system was returned to the initial conditions and maintained until 58 minutes to allow equilibration. The eluted peptides were subsequently introduced into a TIMS-TOF Pro mass spectrometer (Bruker, Bremen, Germany) via electrospray ionization using a captive spray source connected to the column emitter. The instrument was operated in PASEF mode for standard proteomics acquisition. MS/MS scans were initiated 10 × with a total cycle time of 1.16 seconds, a target intensity of 2×10^4^, an intensity threshold of 2.5×10^3^, and a charge state range of 0-5. Active exclusion was enabled for a period of 0.4 minutes and precursors re-evaluated when the ratio of current intensity: previous intensity exceeded 4.

#### Spectral data processing

LC-MS data were processed using MaxQuant software (version 1.16.14.0) with standard settings: trypsin/P as the protease, allowing up to two missed cleavages, carbamidomethylation of cysteine as a fixed modification, and oxidation of methionine as a variable modification. Searches were performed against the *Bifidobacterium breve* DSM20213 proteome database (Uniprot ID: UP000003191, downloaded 2024-11-29). MaxQuant output tables were imported into Perseus (version 2.1.3.0) for statistical analysis. Proteins identified only by site, reverse hits, or potential contaminants were removed. Sample columns were grouped as LNT, LNnT, and GMC. LFQ intensity values were log□□□-transformed, and proteins present in less than 30% of samples in at least one group were excluded. For differential expression analysis, pairwise two-sided Student’s t-tests were performed with a permutation-based false discovery rate (FDR) of 0.05, 250 randomizations, and an S□ parameter of 0.1. The Perseus “Significance B” method was applied, generating a dataset-specific significance curve in the volcano plots. This curve defines the threshold for statistical significance as a function of both –log□□(*p*) and log□ fold change; proteins above this curve were considered significantly differentially expressed. The curve coordinates were exported from Perseus for record-keeping. Protein identifiers were mapped to UniProt entries, and volcano plots were visualized using GraphPad Prism (version 9.0.0.121).

#### Pre-processing of metagenomic reads and MAG reconstruction

To investigate the distribution of the lacZ5 *β*-galactosidase (equivalent to D4BMY8; **Supplementary Figure 16**) and the HMO utilization potential of *Bi breve* (annotated as *Bifidobacterium. breve* DSM20213 = JCM1192 in genomic databases and hereafter also referred to as *B. breve* DSM20213), we analyzed publicly available *B. breve* genomes from the ProGenomes3 database^28^ and metagenome-assembled genomes (MAGs) of both *Bi. breve* and *Bi. infantis* reconstructed from publicly available metagenomic samples from infants of the prospective Amsterdam Infant Microbiome Study (AIMS) cohort (study accession number PRJEB66728). Raw metagenomics reads were pre-processed using BBDuk and BBMap^29^. Raw reads were trimmed to remove adapters, poor quality bases (Q15) and reads shorter than 45bp (minlength = 45). BBMap was used to discard human reads (minID = 0.95) and microbial reads were merged into longer single reads using BBMerge (min overlap=16). The SPAdes^30^ assembler (metagenomic mode) was used for metagenomic assembly. For MAG reconstruction, we used SemiBin for multi-sample binning to generate MAGs (SemiBin 1.0)^31^. For each mother-infant pair, mapped short reads were mapped to a set of assembled contigs (500 bp) from the same mother-infant pair via Bowtie2^32^. Samples were binned in multi-sample binning mode (*SemiBin multi_easy_bin*). MAG quality was estimated using CheckM^32,33^ and GUNC v.0.1^34^, and genomes were taxonomically classified using the GTDB database. Exclusively high-quality *Bi. breve* and *Bi. infantis* MAGs (completeness ≥ 90%, contamination ≤ 5%) were considered suitable for downstream analyses.

#### Pathway enzyme cost analysis

To estimate investment in cellular resources for achieving a certain flux through a pathway we use the enzyme cost. For calculations of the lower bound of the enzyme cost for all enzymes in the pathway, we assumed all substrates are saturated (*K_M_*« [*S*]) and that no inhibition occurred. The only exception was the *β*-galactosidase responsible for cleaving LNT or LNnT, for which substrate saturation was explicitly considered, as differences in substrate affinity are central to our hypothesis.

For the other enzymes, the lower bound of the enzyme cost *E_i_* for reaction *i* in the pathway (ignoring reversibility and regulatory constraints) simplifies to:

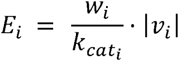

where *w_i_* is the weighing factor of enzyme *i*, (defined as the molecular weight per active site),|*v_i_*| is the flux, relative to the LNT/LNnT flux, of the reaction catalyzed by enzyme *i*, and *E_i_* represents the required enzyme mass per unit of relative flux. We used the BRENDA^35^ and SABIO-RK^36^ enzyme databases and the UniProt^37^ database to obtain values for all enzymatic parameters MW and *k_cat_* (see Data and script availability). When values were unavailable for *Bi. Breve* JCM1192/DSM20213 or other *Bifidobacterium* species, parameters from the phylogenetically closest organism with comparable enzyme function were used. For *β*-galactosidase, substrate saturation was explicitly incorporated because enzyme kinetics differ between LNT and LNnT. Kinetic parameters (*k_cat, LNT_*, *k_cat, LNnT_*, *K_M,LNT_*, *K_M,LNnT_*) reported for *Bi. infantis* DSM20088 = ATCC15697^38^ were used, as full kinetic characterization of *Bi. breve* JCM1192/DSM20213 was not available yet. Intracellular HMO concentration was assumed to be 1 mM. Under these assumptions, the enzyme cost of *β*-galactosidase catalyzing either HMO becomes:

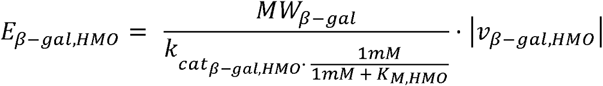

Next, we calculate the total enzyme cost of a pathway *C_P_* as:

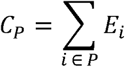

where *P* denotes the set of reactions in a pathway. In the main text, *P* = formate refers to the set of reactions from the HMO to formate production, and *P* = lactate refers to the set of reactions to lactate production, respectively.

#### Profiling of *β*-galactosidase genes in isolate genomes and MAGs of *Bi. breve*

Proteomes for all genomes and MAGs were predicted using Prodigal^39^. For *β*-galactosidase gene identification and annotation, we used *bifidoAnnotator* (https://github.com/nicholaspucci/bifidoAnnotator), a specialized tool for fine-grained annotation of bifidobacterial enzymes involved in human milk glycan utilization. Briefly, *bifidoAnnotator* leverages MMseqs2 to map protein sequences against a manually curated database of 22,699 bifidobacterial enzymes involved in HMG-utilization. We applied a modified version of the *bifidoAnnotator* pipeline with default settings (coverage ≥ 0.50, bitscore ≥ 200) restricting outputs to GH2 and GH42 family *β*-galactosidases in all *Bi. breve* and *Bi. infantis* genomes/MAGs.

#### Plasmid and strain construction

The *β*-galactosidase gene fragments were PCR-amplified from the *Bi. breve* JCM1192/DSM20213 genome using Phusion High-Fidelity PCR Master Mix (Thermo Scientific) to ensure accurate amplification. Each 20 μL reaction contained 10 μL 2× Phusion Mix, 0.5 μM of each primer (primers for each gene are listed in Supplementary Table 4), 0.5 μL *Bi. breve* JCM1192/DSM20213 genomic DNA (or 1 μL colony lysate), and 9 μL of nuclease-free water. Cycling conditions consisted of an initial denaturation at 98°C for 3 min; 30 cycles of 98 °C for 15 s, annealing at 60 °C for 20 s, and extension at 72 °C for 15 s; followed by a final extension at 72 °C for 5 min. PCR products were purified with the Monarch® PCR & DNA Cleanup Kit (NEB). For construction of pYS01, the BIFBRE_03435 fragment (D4BMY8 gene) and the pET28a plasmid were digested with BamHI and NdeI at 37 □°C for 1 h. For construction of pYS02 and pYS03, HindIII and NdeI were used to digest the corresponding gene fragments (BIFBRE_04539 as D4BR09 gene; BIFBRE_03324 as D4BMM7 gene) and pET28a plasmids. After digestion, inserts and vector backbones were gel-purified using the Monarch® DNA Gel Extraction Kit (NEB). The purified fragments and backbones were then ligated with T4 DNA ligase at 16 °C for 4 h. Ligation mixtures were transformed into *Escherichia coli* DH5α competent cells by heat shock. After recovery at 37 °C, DH5α cells were plated on LB agar containing kanamycin (50 mg/L) and incubated at 37 °C overnight. Colonies were randomly picked and screened by colony PCR targeting the BIFBRE_03435, BIFBRE_04539, and BIFBRE_03324, respectively. Positive colonies were retained as plasmid carriers. Plasmids were prepared by culturing the carriers in LB broth supplemented with kanamycin (50 mg/L), followed by extraction using the Monarch® Plasmid Miniprep Kit (NEB). Purified plasmids were Sanger sequenced to confirm the absence of mutations (sequencing primers are listed in Supplementary Table 4), and verified constructs were transformed into *E. coli* BL21 by heat shock for recombinant protein production. Plasmid maps are shown as **Supplementary Figure 13**.

#### Purification of *β*-galactosidase

The *E. coli* BL21 strains were revived using an LB agar plate with 50 mg/L kanamycin. After overnight growth, a single colony was picked to grow in liquid LB medium overnight. After that, 100 µL of *E. coli* was inoculated into 10 mL LB-media containing 50 mg/L kanamycin and incubated at 37 °C while shaking at 200 rpm (New Brunswick Innova44 M1282-0000). OD_600_ was measured every 2 h until it reached 0.6. Then, isopropyl *β*-d-1-thiogalactopyranoside (100 mM IPTG, ThermoFisher Scientific R1171) was added to a final concentration of 50 μM, whereafter the flask was incubated overnight at 25 □ and 200 rpm. To harvest the cells, the liquid culture was transferred to a 15 mL conical tube and centrifuged at 12,000 g and 4 □ for 5 min, and the pellet was resuspended with 500 µl His-binding buffer (Zymo Research, P2003-3) from the His-Spin Protein Miniprep kit (Zymo Research, P2001). The resuspended cells were then transferred to a screw-cap tube containing 0.17 - 0.18 mm glass beads (approx. 40% of volume), and lysed using a bead beater for 3 cycles at 6000 rpm for 30 sec per cycle. The tubes were kept on ice in between cycles to avoid thermal denaturation of the protein. The cell lysates were then centrifuged at 21,500 g and 4 □ for 5 min, and 450 µl supernatant was used to purify the *β*-galactosidase using the His-Spin Protein Miniprep kit according to the manufacturer’s protocol. After using the kit, the purity of the sample was confirmed using an SDS-PAGE gel, and the identity of the purified protein was confirmed using LC-MS, as described above.

The SDS-PAGE gel was run using the SurePAGE™ kit (GenScript, M00656) following the instructions of the manufacturer’s protocol. Protein samples were prepared by resuspending 10 µL of 2× Laemmli Sample Buffer (Bio-Rad, 1610737) with 10 µL of the protein sample, followed by denaturing at 95 □°C for 5 minutes (ThermoFisher Thermomixer, 5436). For gel electrophoresis, one pre-cast gel (Genscript SurePage Gels, M00677) was loaded with 5 µL of a PageRuler Prestained Protein Ladder (ThermoFisher, 26619), followed by 10□µL of the protein sample in separate wells. Electrophoresis was initiated at 60□ V until clear band formation was observed, after which the voltage was increased to 110□V for an additional 30 minutes. Finally, the gel was stained with InstantBlue^®^ for 20 minutes, washed, and imaged.

To confirm the enzymatic activity of the purified proteins, they were individually dissolved in PBS + 0.05% Tween20 after which X-Gal (Duchefa, X1402.5000) dissolved in DMSO was added alongside a blank without X-Gal. Significant differences in colouration were observed and the method was thus deemed successful to harvest and purify the *Bi. breve* derived *β*-galactosidase from the transfected BL21 strain.

#### Enzyme kinetic assay and quantification of LNT and LNnT

To measure the enzyme kinetics of the purified *β*-galactosidases in reaction with LNT and LNnT, HPAEC was used to measure the concentration of LNT, LNnT, and galactose. In brief, 3 purified *Bi. breve β*-galactosidases (0.15 mg/ml), D4BMY8, D4BR09 and D4BMM7, were individually incubated with 1 m M LNT or LNnT at 37 □ for 2 h. Reactions were sampled every 15 min during the incubation. The samples were kept on ice to stop the reaction. After 2 h, samples were heat-shocked for 10 min at 80 □ to denature the enzyme, and then diluted 100 times in ddH_2_O and transferred to Verex vials (Phenomenex, AR0-39P0-13). Standard curves were made individually using LNT, LNnT, and galactose at concentrations of 0, 0.1, 0.5, 1.0, and 2.0 mM, respectively. Data were processed as described in HPAEC.

#### *In silico* docking model

For the model, an AutoDock Vina pipeline was used^40,41^. The three-dimensional structure of *Bi. infantis β*-galactosidase and LNT were extracted from the protein data bank (PDB) file 8IBT (*Bifidobacterium infantis β*-galactosidase complexed with LNT crystal structure). Vina requires the definition of a 3-dimensional docking grid, which specifies the region of the protein where the ligand will be sampled for low-energy conformations based on predicted binding affinities. The docking grid was defined to encompass the entire active site, using a box size of 25 × 30 × 24 Å centered at coordinates (x = 73.000, y = −27.800, z = −61.900) by measuring the distance between the residues of the active site in a 3-dimensional manner^42^. Docking analysis was then performed in triplo, using an exhaustiveness of 30 on the center location of the active site. Residues comprising the active site were made flexible, with the exception of residues 200 and 221 in chain B, which were kept rigid. All other settings were kept at their defaults. Generated LNT conformations yielded an average RMSD of 1.7 in the top scoring positions compared to the crystal structure LNT

#### *In silico* docking of LNT and LNnT to *Bi. infantis* and *Bi. breve β*-galactosidase

The three-dimensional structure of *Bi. breve β*-galactosidase (UniProt D4BMY8) was predicted using AlphaFold3, while the LNnT ligand structure was extracted from PDB entry 5UF4 using PyMOL. Similarly, the crystal structures of *Bi. infantis β*-galactosidase in complex with LNT were obtained directly from PDB entry 8IBT. For molecular docking, residues comprising the active site were made flexible, with the exception of residues 200 and 221 in chain B, which were kept rigid. The docking grid was defined to encompass the entire active site, using a box size of 25 × 30 × 24 Å centered at coordinates (x = 73.000, y = −27.800, z = −61.900). Docking was performed using AutoDock Vina using an exhaustiveness value of 30, while other settings were kept at default values. Hydrogen bonds between the ligand and protein were identified in PyMol using the ‘Find Polar Contacts’ tool. Distance between the ligand and catalytic residues were calculated using heavy atoms.

### Data and script availability

The LC-MS datasets presented in this study are deposited in the ProteomeXchange repository accession number: PXD076309. The raw sequencing data generated in this study have been deposited in the European Nucleotide Archive (ENA) under the accession number PRJEB111408. The AIMS bifidobacterial MAGs are deposited in Zenodo (DOI: <u>10.5281/zenodo.17206992</u>). The modified bifidoAnnotator script underlying data, enzyme cost analysis script and parameters, and script for molecular docking used in this study are available at https://github.com/nicholaspucci/Shan_et_al_HMO_SynCom. The *bifidoAnnotator*’s source code and documentation are available at https://github.com/nicholaspucci/bifidoAnnotator.

### Usage of Large Language Models

ChatGPT (GPT-5.3) and UvA AI Chat (based on GPT-5.1) were used for manuscript polishing.

## Results

### HMOs differentially alter the composition and metabolic function of infant gut bacterial community iBaCo

To study the effects of individual HMOs on the ecology and function of infant gut microbiota, we assembled an infant <u>Ba</u>cterial <u>Co</u>mmunity (iBaCo) using eight representative infant gut species: *Bifidobacterium pseudocatenulatum*, *Bifidobacterium bifidum*, *Bifidobacterium longum subsp. infantis*, *Bifidobacterium breve* JCM1192/DSM20213, *Collinsella aerofaciens*, *Anaerostipes caccae*, *Blautia luti*, and *Bacteroides fragilis*, and allowed the iBaCo inoculum to self-assemble in YCFA media containing individual HMOs, with No-sugar as a negative control (**Figure 1A**). After 24 h, we observed a baseline growth in the absence of carbohydrate (No-sugar), with a slightly increased OD_600_ (0.10 ± 0.06). In contrast, iBaCo clearly grew as a community in the presence of each HMO, with OD_600_ values increased to 0.37 ± 0.16 (2’-FL), 0.40 ± 0.15 (3-FL), 0.31 ± 0.13 (DFL), 0.38 ± 0.07 (3’- SL), 0.40 ± 0.04 (6’-SL), 0.49 ± 0.09 (LNT), and 0.47 ± 0.08 (LNnT), respectively (**Figure 1B**) and each HMO was completely consumed by iBaCo (**Supplementary Figure 1**). Correspondingly, we found that the total live bacterial density was higher in all HMOs compared to that in the absence of carbohydrate, albeit similar among HMOs (**Figure 1C**). For example, in the presence of LNT or LNnT, the live bacterial density increased from 1.0 × 10^7^ CFU/mL to 6.2 × 10^8^ and 4.7 × 10^8^ CFU/mL, respectively, significantly higher than that in the absence of carbohydrate (No-sugar) (**Figure 1C**).

**Figure 1.**
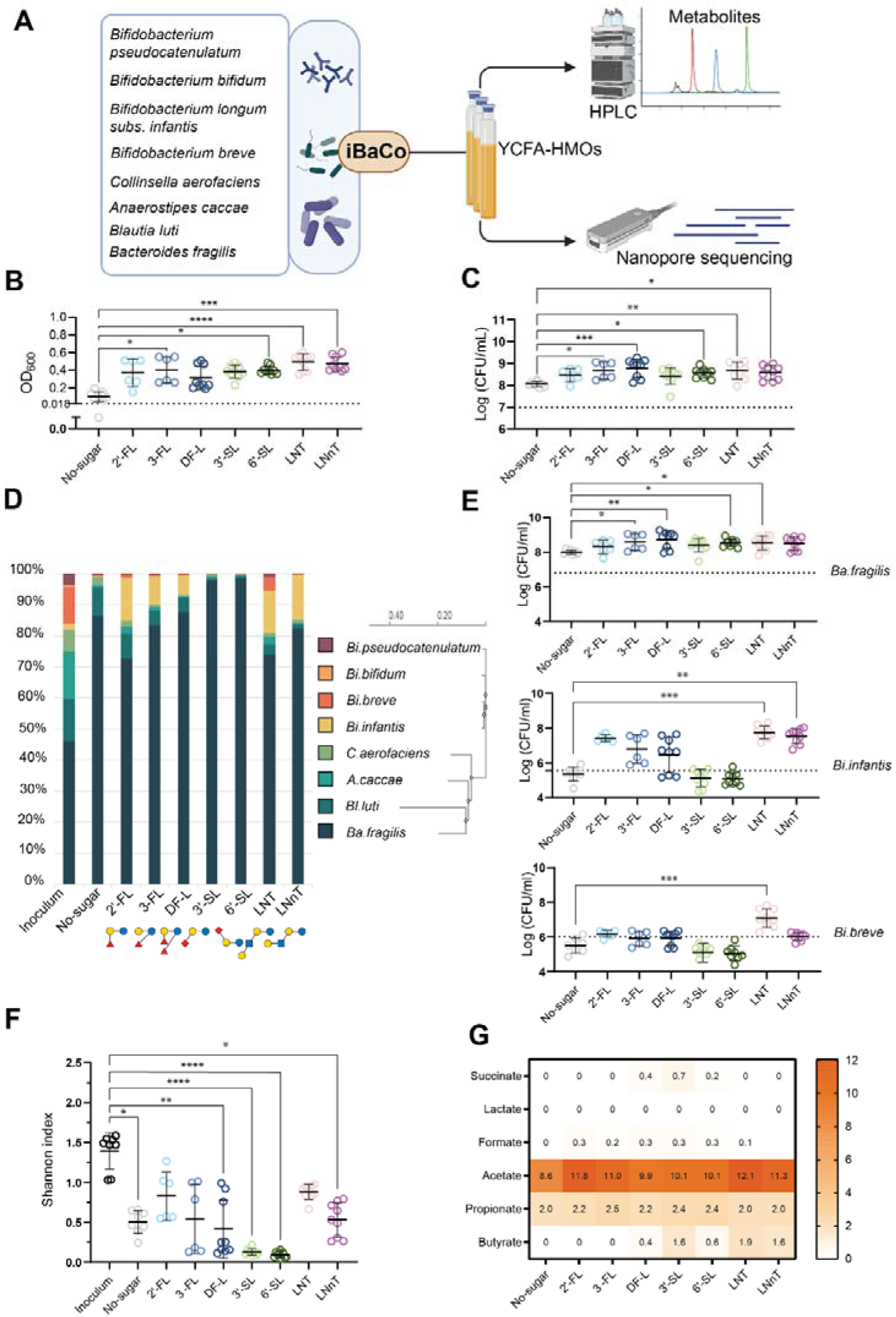
Assembly, compositional and functional characterization of iBaCo in the presence of HMOs. (A) Schematic workflow of iBaCo assembly and characterization. (B) The OD_600_ values of iBaCo after 24 h of culturing. Dashed lines indicate the starting OD_600_ after inoculation. (C) Total live cell density (bacterial colony forming unit [CFU]/mL) of iBaCo after 24 h of culturing. The dash line indicates the cell density at t = 0 h. (D) Relative abundance of each species in iBaCo. (E) Absolute bacterial density (CFU/mL) of *Ba. fragilis*, *Bi. infantis* and *Bi. breve* in iBaCo. The dash line indicates the cell density at t = 0 h. (F) Shannon index of iBaCo grown in different HMOs. (G) Net production of short chain fatty acids and organic acids by iBaCo in the presence of different HMOs. Error bars in B-C and E-F represent mean ± standard deviation. The Kruskal-Wallis test with Dunn’s multiple comparisons test was performed. *p < 0.05; **p < 0.01; ***p < 0.001; ****p < 0.0001. Data in (B-G) are based on three independent experiments with at least 3 technical replicates in each experiment.

At the community level, HMOs differentially impact the diversity and composition of iBaCo. Among all HMOs, *Ba. fragilis* is a dominant species in iBaCo. Compared to the inoculum, the relative abundance of *Ba. fragilis* was 73-88% in the presence of fucosylated HMOs, i.e. 2’-FL, 3-FL DFL, and neutral HMOs, i.e. LNT and LNnT, (**Figure 1D**), which is lower than that in the presence of sialylated HMOs, accounting for >97% relative abundance. This higher dominance of *Ba. fragilis* in sialylated HMOs is likely not due to its higher competence under these conditions, evidenced by the similar absolute *Ba. fragilis* density among all HMOs (**Figure 1E**), suggesting that the higher relative abundance of *Ba. fragilis* is caused by the decrease of other species in iBaCo. Indeed, we observed that the cell density of *Bi. infantis* and *Bi. breve* in sialylated HMOs were similar or even lower than that in the absence of carbohydrate, and much lower compared to other HMOs (**Figure 1E**). The loss of these species in iBaCo is also reflected in the Shannon diversity (0.13 ± 0.04 of 3’-SL and 0.10 ± 0.04 of 6’-SL, **Figure 1F**), suggesting that both sialylated HMOs do not support these bifidobacterial species and can not maintain a diverse community. It is worth noting that the combination of relative abundance and absolute live cell density data, compared to relative abundance alone, offers a more complete picture of compositional changes in a SynCom.

*Bi. infantis* is one of the most abundant bifidobacterium species in the infant gut. Despite the dominance of *Ba. fragilis* in the iBaCo communities under all cultured conditions, Bifidobacterium species were maintained within iBaCo, accounting for up to 20% of the community (**Figure 1D**). Therefore, iBaCo remains effective to investigate the impact of HMOs on bifidobacterial function in a community. Here, we found HMO- and species-dependent effects in iBaCo. For example, both fucosylated and neutral HMOs maintained *Bi. infantis* at proportionally high relative abundance ( 6.4-13.9% and 13.7-14.4%, **Figure 1D**), cell density (1.30-2.90 × 10^7^ and 4.80-7.74 × 10^7^ CFU/mL, **Figure 1E**), and higher level of acetate (**Figure 1G**), compared to sialylated HMOs. HMOs did not significantly affect the formate and propionate production by iBaCo. However, butyrate production, albeit low, was observed at a significantly higher level in LNT and LNnT (**Figure 1G** and **Supplementary Figure 3**). The effect on the other two Bifidobacterium species, i.e. *Bi. breve* and *Bi. pseudocatenulatum*, is highly HMO-specific. Among all HMOs, only LNT was able to maintain the relative abundance and high cell density of *Bi. breve* and *Bi. pseudocatenulatum* in iBaCo (**Figure 1D, 1E** and **Supplementary Figure 2**). This is in agreement with the clinical observation that both *Bi. breve* and *Bi. pseudocatenulatum* have higher relative abundance in infants with 3-6 or >6-month breast milk feeding compared to those with <3 month breast milk feeding^43^. Together, these results suggest that fucosylated and neutral HMOs but not sialylated HMOs can effectively maintain *Bi. infantis*, whereas only LNT can maintain *Bi. breve* and *Bi. pseudocatenulatum* in iBaCo.

### Substrate preference on the monoculture growth largely explains species behavior in iBaCo community

To investigate the contribution of individual species to the community and metabolic response to HMOs, we characterize the growth and metabolite profile of the eight species in the presence of HMOs as the sole carbohydrate and found their growth and utilization pattern of HMOs can largely explain their behavior in iBaCo. For example, *Ba. fragilis* grew to a similar yield among all HMOs with similar maximum growth rates (0.12-0.28 h^-1^) but different lag phases (**Supplementary Figure 8**). Thin layer chromatography confirmed that *Ba. fragilis* individually consumed all HMOs, including sialylated HMOs (**Supplementary Figure 10**). This is in agreement with the previous observation that *Ba. fragilis* grew well in a mixture of milk oligosaccharides using the mucus glycan-degrading enzymatic machinery^44^ and a sialidase is essential for its colonization and dominance in the gut microbiota of mice in early life^45^.

For all Bifidobacterium species, i.e. *Bi. infantis*, *Bi. bifidum*, *Bi. breve*, and *Bi. pseudocatenulatum*, the growth pattern and maximum growth rate (**Supplementary Figure 8**) highly matches with their cell density in iBaCo (**Figure 1E**). Interestingly, *Bi. infantis* exhibited a broad capacity to utilize HMOs, although utilization of 3′-SL and 6′-SL remained incomplete and supported only limited growth. In contrast, *Bi. breve* and *Bi. pseudocatenulatum* behaved as more specialized HMO utilizers, displaying a pronounced growth preference for LNT over LNnT (**Figure 2A** and **Supplementary Figure 6**). The maximum growth rate of *Bi. breve* was 0.24 ± 0.12 vs. 0.08 ± 0.04 h^-1^ in LNT vs LNnT, while the lag phase 4.63 ± 2.15 h vs 33.25 ± 23.66 h (**Supplementary Figure 8**). In agreement, for *Bi. Breve*, both LNT and LNnT were consumed, whereas for *Bi. pseudocatenulatum*, only LNT was consumed (**Supplementary Figure 10**). This preferential utilization of LNT and fast growth in LNT largely explain the higher cell density of *Bi. breve* and *Bi. pseudocatenulatum* in iBaCo grown in LNT rather than LNnT and other HMOs.

**Figure 2.**
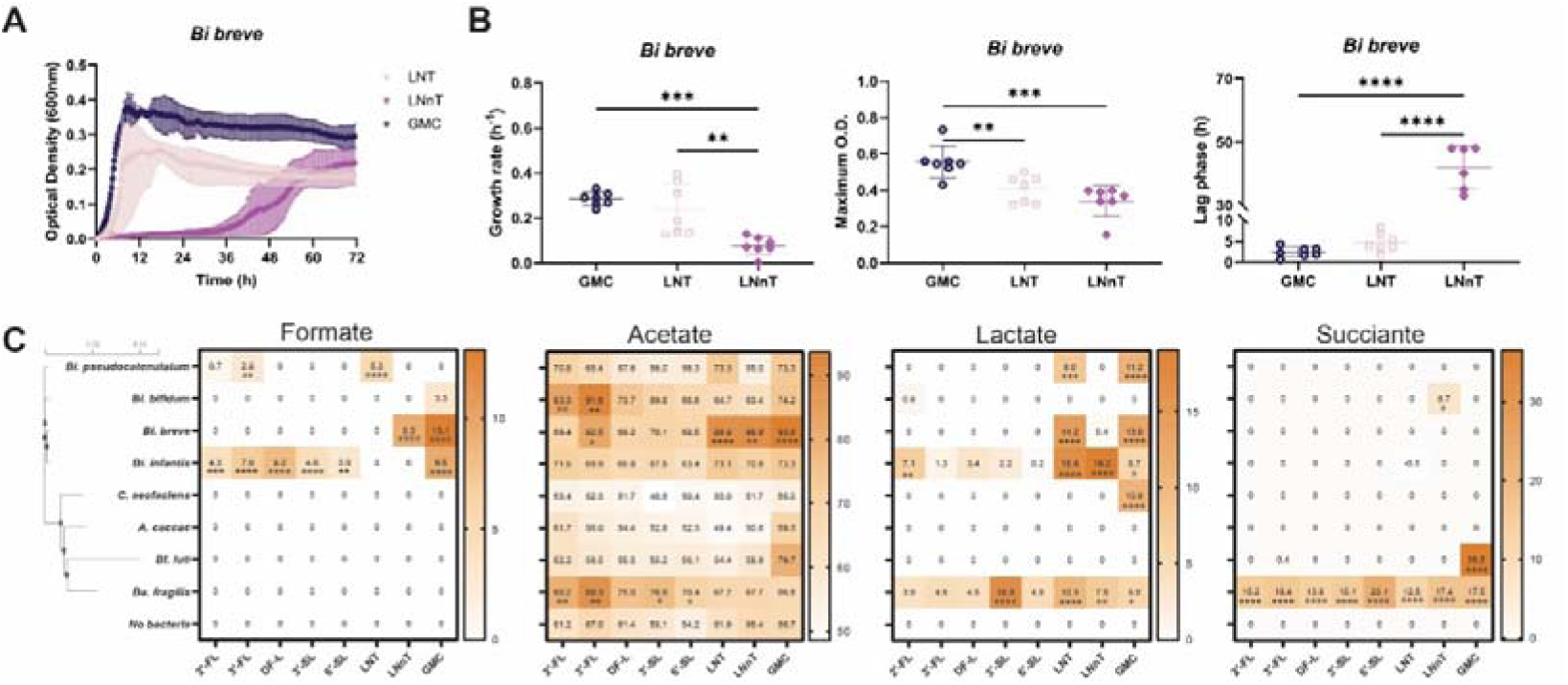
Growth pattern and production of organic acids by individual iBaCo species in monoculture. (A) Growth curves of *Bi. breve* in the presence of LNT, LNnT, and GMC. (B) The growth rate, maximum OD, and lag phase of *Bi. breve* in the presence of LNT, LNnT, and GMC. Statistical analysis was performed using one-way ANOVA. (C) Concentration of formate, acetate, lactate, and succinate in the presence of different HMOs. Glucose (GMC) as the positive control. Data represent the mean of three independent experiments, with individual replicates shown as points (mM). Statistical analysis was performed by two-way ANOVA followed by Dunnett’s test, comparing each bacterial condition to the no-bacteria control for each HMO. * p < 0.05; ** p < 0.01; *** p < 0.001; **** p < 0.0001.

The substrate preference on growth is coupled with distinct pattern of short chain fatty acids and organic acid production by these infant gut bacterial species. For example, *Bi. breve* produced high levels of formate, lactate and acetate in glucose (**Figure 2C**). However, it preferred to accumulate lactate (14.2 mM) and acetate (26.9 mM) but not formate after consuming LNT, whereas it produced formate (9.3 mM) and acetate (26.9 mM) but not lactate in LNnT (**Figure 2C**). Similar preference on the metabolite production was observed for *Bi. pseudocatenulatum* (**Figure 2C**). Interestingly, this substrate preference is also apparent for *Ba. fragilis*. For instance, *Ba. fragilis* produced higher levels of lactate (16.8 vs. 4.9 mM) and succinate (15.1 vs. 23.1 mM) in 3′-SL than 6′-SL. These results suggest that the small difference on the HMO structures, e.g. *β*1-3 versus *β*1-4 galactosyl linkage between LNT and LNnT, could lead to distinct utilization patterns by specific infant bacteria.

### LNT-inducible ***β***-galactosidase D4BMY8 has substrate preference on LNT over LNnT

Next, we sought to investigate which molecular machinery contributes to the distinctive metabolic output by these infant gut bacteria. We leveraged the most striking difference between LNT and LNnT for *Bi. breve* and compared the proteome of *Bi. breve* at its late exponential growth in LNT versus LNnT. To utilize LNT or LNnT, *β*-galactosidases are required to initiate the degradation and release galactose, followed by *β*-N-acetylhexosaminidase to release N-acetylglucosamine, and finally degrade lactose (**Figure 3A**). Genome mining or functional annotation of the *Bi. breve* genome (GenBank assembly: ACCG02000012, JCM1192/DSM20213) identified putative genes that are potentially involved in the transport and hydrolysis of LNT or LNnT (**Figure 3B**). In particular, a GH42 *β*-galactosidase gene (BIFBRE_03435) is located next to three ABC transporter genes (BIFBRE_03432, BIFBRE_03433, BIFBRE_03436) (**Figure 3B**). Proteomic analysis confirmed a high-level expression of these genes in *Bi. breve* grown in LNT and LNnT, whereas little expression in glucose (**Figure 3C**, **Supplementary Table 8 and 9**), suggesting that these proteins are involved in LNT and LNnT utilization and inducible to a similar extent by LNT and LNnT. Interestingly, among four *β*-galactosidases in the genome, only *β*-galactosidase D4BMY8, but not D4BMM7 and D4BR09, was significantly higher in LNT versus LNnT (**Supplementary Figure 12**), hinted that D4BMY8 is essential for LNT utilization. In addition, the *β*-N-acetylhexosaminidase (D4BR02, BIFBRE_04532) was expressed at slightly higher level in both LNT and LNnT compared to glucose. These data suggest that the proteins that are involved in the release of galactose and N-acetylglucosamine from LNT or LNnT are present in *Bi. breve*. Consistently, the gene clusters using galactose and N-acetylglucosamine are also highly and inducibly expressed in *Bi. breve* grown in LNT and LNnT (**Figure 3D**). For example, in the Leloir pathway of galactose utilization^46,47^, GalK (D4BMU0, phosphorylation of galactose), GalT (D4BL97, D4BMT9) and GalE (D4BQW0, formation of Glc-1-P) were expressed at significantly higher level in LNT/LNnT versus glucose (**Figure 3D**). Interestingly, the enzymes involved in the downstream utilization of Glc-1-P, known as Embden-Meyerhof pathway, i.e. Pgm (D4BQV9, Glc-1-P to Glc-6-P), Pgi (D4BMK3, Glc-6-P to Fruc-6-P), PfkB (D4BS00, Fruc-6-P to Fruc-1,6-P2) and FbaA (D4BLN9, Fruc-1,6-P2 to glyceraldehyde-3-P), are expressed at similar level among LNT, LNnT and glucose (**Figure 3D**). Similarly, most enzymes involved in GlcNAc metabolism such as NahK (D4BQW9, GlcNAc phosphorylation) deacetylated by NagA (D4BS05, GlcNAc-6-P deacetylation), and NagB (D4BS04, GlcN-6-P aldo-keto isomerization) were highly expressed only in LNT and LNnT but not detectable in glucose. Taken together, these results suggest that the utilization of LNT/LNnT and their hydrolyzed monosaccharides galactose and GlcNAc are highly inducible in *Bi. breve*, and that the enzyme levels alone can not explain the differences in metabolic output between LNT and LNnT.

**Figure 3.**
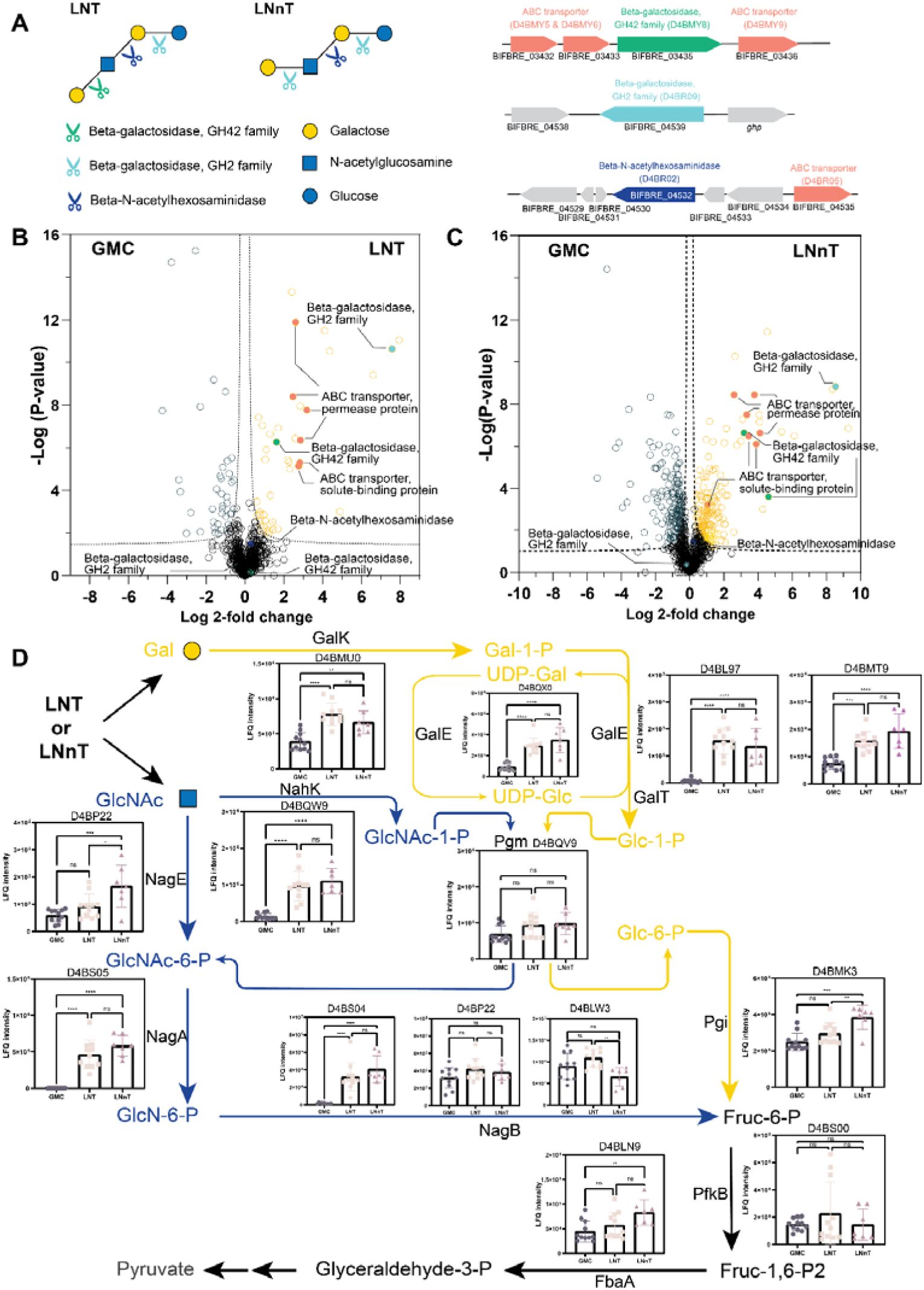
Proteomic and genomic insights into *Bi. breve* metabolism of LNT and LNnT. (A) Cleavage pathways of LNT and LNnT with annotated putative gene clusters. (B–C Volcano plots showing differential protein expression under LNT (B) and LNnT (C) versus GMC conditions. Proteins significantly upregulated in LNT/LNnT are outlined in gold, whereas those enriched in GMC are outlined in teal. Functional classes are highlighted: ABC transporters (coral), GH2 *β*-galactosidases (mint), GH42 *β*-galactosidases (green), and *β*-N-acetylhexosaminidases (blue). *Dashed lines indicate significance thresholds*. (D) Expression profiles of proteins involved in downstream galactose and GlcNAc metabolism. LFQ intensities were analyzed by one-way ANOVA. * p < 0.05, ** p < 0.01, *** p < 0.001; **** p < 0.0001.

The surprisingly similar level of LNT/LNnT utilization enzymes in *Bi. breve* hinted that the intracellular level of these proteins is not a major contributor of LNT/LNnT’s different effects on *Bi. breve* growth and metabolic patterns. We hypothesized that the *Bi. breve β*-galactosidases have substrate preference on LNT over LNnT. To test this, we overexpressed BIFBRE_03435, BIFBRE_04539 and BIFBRE_03324 genes in IPTG-inducible plasmid in *E. coli* BL21 and purified the protein. The corresponding purified *β*-galactosidases (D4BMY8, D4BR09 and D4BMM7) were incubated with LNT or LNnT at 37 □ for 2 hours, and the concentrations of LNT or LNnT and their hydrolysis product galactose were quantified every 15 minutes (see Methods). For D4BMY8, LNT was degraded faster than LNnT at 1 mM. Correspondingly, the concentration of galactose in LNT increased faster than that in LNnT (**Figure 4A**), confirmed that D4BMY8 has a clear substrate preference. In contrast, D4BR09 exhibited a substrate preference for LNnT over LNT with a much slower hydrolysis (**Supplementary Figure 15**), while the other GH42 family *β*-galactosidase D4BMM7 was unable to hydrolyze either LNT or LNnT (**Supplementary Figure 15**). Finally, knockout of D4BMY8 ortholog *lacZ5*/*lntA* in *Bi. breve* UCC2003 completely abolished its ability to grow in LNT but not in LNnT, while knockout of other *β*-galactosidases did not affect the growth (**Supplementary Figure 16**), confirming indispensable and specific role of this gene in LNT utilization *in vivo* in *Bi. breve* cells.

**Figure 4.**
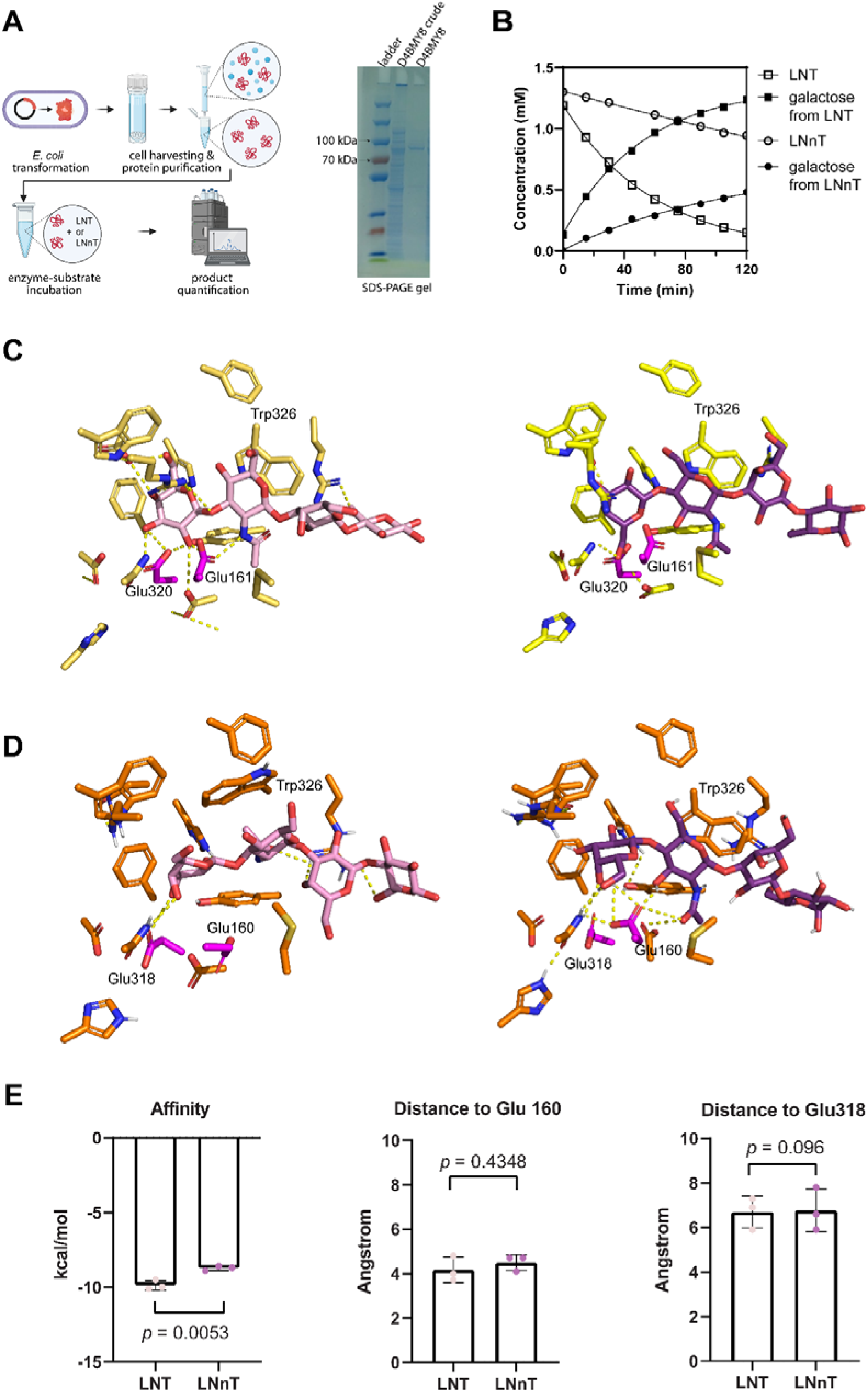
Enzyme-substrate kinetics and molecular docking for LNT or LNnT with *Bi. breve β*-galactosidase. (A) schematic illustration of *β*-galactosidase purification and enzymatic assay. (B) concentrations of substrate (LNT or LNnT) and its hydrolysis product galactose over 2 hour incubation with the purified *β*-galactosidase D4BMY8. (C) LNT (yellow protein, light pink LNT, as positive control) and LNnT (yellow protein, purple LNnT) in complex with *Bi. infantis β*-galactosidase (PDB: 8ibt). (D) *Bi. breve* D4BMY8 with docked LNT (orange protein, light pink LNT) or LNnT (orange protein, purple LNnT). Magenta sticks represent catalytic residues. (E) Binding energy (ΔG, kcal/mol) of LNT and LNnT with D4BMY8, and their distance (Angstrom) to catalytic amino acid residues in D4BMY8. Data represent three docking runs. Statistics were performed using Student’s t-test.

Next, we sought to investigate the biochemical basis of the substrate specificity of D4BMY8. Since LNT and LNnT have the same composition of monosaccharides and only one difference on the beta-galactosyl linkage, we hypothesize that this conformational difference interferes with their interaction with the amino acids in the catalytic pocket of D4BMY8. To test this, we performed molecular docking of D4BMY8 with LNT or LNnT as ligand molecules, and evaluated their biochemical interactions by calculating the binding energy and the number of hydrogen bonds (see Methods). For molecular docking, we first used the experimentally confirmed crystal structure of *Bi. infantis β*-galactosidase (PDB: 8IBT, **Figure 4C** and **Supplementary Table 7**)^42^, and confirmed that the affinity between LNT and 8IBT is higher than LNnT. Between LNT and 8IBT, 12 hydrogen bonds could be formed, but only 4 between LNnT and 8IBT. Similarly, for *Bi. breve β*-galactosidase D4BMY8, its predicted affinity with LNT is higher than LNnT, (−9.9 ± 0.26 kcal/mol v.s. −8.7 ± 0.12 kcal/mol, *p* = 0.005) (**Figure 4E**). Interestingly, both substrates have a similar number of hydrogen bonds with residues in the active site and similar distance to the catalytic glutamate residues (**Figure 4D**). Notably, the residue Trp326 exhibited a pronounced vertical inversion in the presence of LNnT compared with LNT (**Figure 4D**), while other residues in the catalytic pocket showed similar conformations for both substrates, indicating this residue might be essential for stabilizing the substrate. Together, these results give insight into the key role of *β*-galactosidase D4BMY8 in preferential utilization of LNT over LNnT.

### Substrate preference and enzyme costs explain the preferential production of lactate or formate

Together with the proteome and substrate preference of *β*-galactosidase observed above, a shift in metabolic end products was observed: lactate was accumulated when *Bi. breve* cells were grown in LNT, whereas formate was accumulated in LNnT. Since the protein levels of the metabolic pathways are not substantially different, we hypothesize that the fast and slow release of galactose from LNT or LNnT differentially drive the metabolic flux in *Bi. breve* cells. Here, we calculated and compared the enzyme costs per ATP flux on LNT and LNnT. We found that the net ATP gain per unit of enzyme for a flux unit of HMO differs for the two pathways for growth on LNT versus LNnT. If *Bi. breve* produces acetyl-CoA and formate from pyruvate (using Pfl), it generates an additional two ATP through phosphorylation of acetyl-phosphate (**Figure 5A**). However, this comes at the cost of using five enzymes (PflB, Pta, AckA, Aldh, Adh) after the pyruvate (switching) node, instead of a single enzyme (Ldh) producing lactate. The higher cost associated with the degradation of LNnT, because of the lower *k*_cat_ and affinity, pushes the total enzyme costs of both branches closer together (**Figure 5B**). We hypothesized that the net ATP gain (formate branch: 9, lactate branch: 7) at the cost of more enzymes could be the reason *Bi. breve* switches strategy in a less favorable environment, as LNnT is harder to degrade. Therefore, we quantified this by calculating *G_P_*, the net ATP yield over the enzyme cost of the pathway (*C_P_*) using 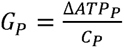, where *p* refers to the set of reactions in the pathway (formate/lactate refers to the complete set of reactions from the HMO to the product).

**Figure 5.**
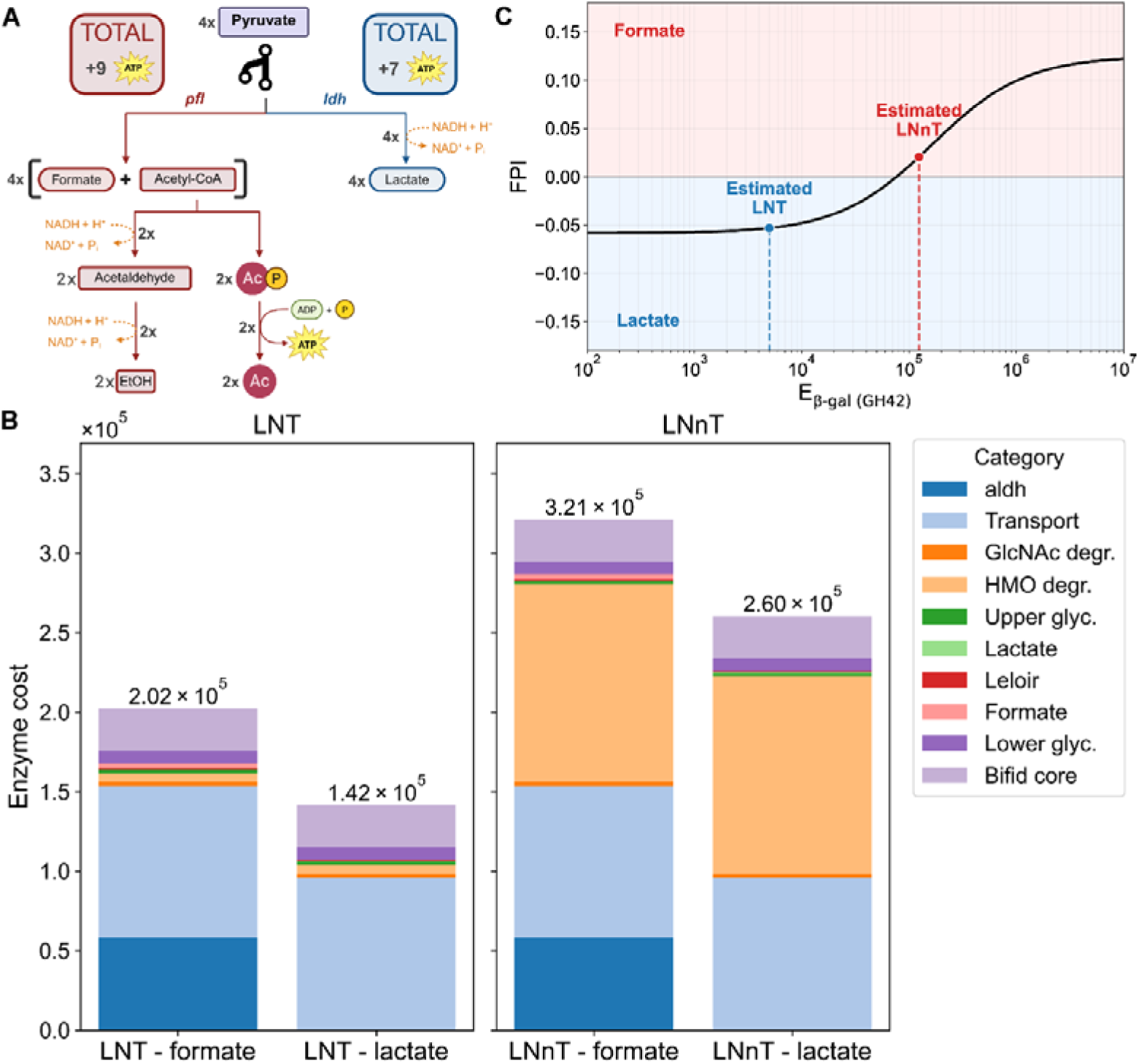
Enzyme cost analysis and pathway preference during the LNT and LNnT fermentation by *Bi. breve*. (A) Illustration of the metabolic network after the branching node pyruvate and downstream by-products. (B) The total enzyme costs for the pathways related to the specific substrate (LNT vs LNnT) and the branch chosen. The enzyme cost of the Aldh enzyme, which is a step downstream of the formate branch, is shown as a different category, because *k_cat, Aldh_* has a significantly lower value compared to other values in the complete pathway. (C) The Formate Preference Index (FPI) was calculated as the normalized difference 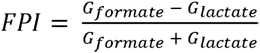 and shown here as a function of the enzyme cost for *β*-galactosidase (E_*β*-gal_). Estimated costs for *β*-galactosidase cleaving LNT and LNnT are shown with the dashed lines and dots.

To understand the metabolic preference for one product over the other, we calculated the normalized difference between the values obtained for *G*_formate_ and *G*_lactate_ as the formate preference index 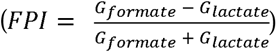. This index is above zero when using the formate branch leads to a lower pathway cost per ATP and below zero otherwise. We analyzed this index for a range of enzyme costs of *β*-galactosidase (**Figure 5C**). We observed that increasing the enzyme cost of *β*-galactosidase can indeed drive a metabolic switch to the formate branch (FPI > 0), and that the estimated enzyme cost of LNnT, *E*_*β*-gal, LNnT_, resides in this domain. Consistently, we observed a higher protein level of PflB in LNnT than LNT. These results indicate that the formate branch gains more ATP but has a higher enzyme cost. The higher enzyme cost associated with LNnT degradation increases the total pathway cost for both the formate and lactate branches. Consequently, while the absolute cost difference between the two branches remains unchanged, their relative difference in total enzyme cost is reduced compared to LNT utilization. This compression of relative costs shifts the benefit, defined as ATP yield relative to total pathway enzyme investment, in favor of the formate branch.

This analysis is intentionally coarse-grained and local. It ignores regulatory effects, changes in global proteome allocation such as ribosome investment, and metabolite channeling through multi-enzyme complexes. Taken together, these results support a scenario in which *Bi. breve* prefers the higher-yield, formate-producing branch to compensate for the high enzymatic cost of slowly hydrolyzed substrates such as LNnT, whereas the kinetically cheaper LNT can be directed through a faster but lower-ATP-yield lactate fermentation.

### LNT-preferential ***β***-galactosidase is divergently distributed in infants

To investigate the broader distribution of *β*-galactosidase genes across the *Bi. breve* species, we conducted a comprehensive analysis of 147 *Bi. breve* isolate genomes and metagenome-assembled genomes (MAGs). Using a targeted annotation approach to identify distinct *β*-galactosidase genes, we found all investigated *Bi. breve* genomes and MAGs (hereafter referred to as genomes) carrying at least one *β*-galactosidase gene, with an average 5.3 ± 1.2 (mean ± SD) *β*-galactosidase genes, specifically 2.9 ± 0.9 GH2 and 2.4 ± 0.5 GH42. We identified a total of 8 *β*-galactosidase variants across all genomes, 5 belonging to the GH2 family (*lacZ2*, *lacZ3*, *lacZ6*, *lacZ7a* and *Bga2D*) and 3 to the GH42 family (*lacZ4*, *lacZ5* and *Bga42C*; **Figure 6A**). Notably, *lacZ5* and *lacZ6* homologs were identified in 146 (99%) and 145 (99%) of investigated *Bi. breve* genomes (**Figure 6A**). The reference strain *Bi. breve* JCM1192/DSM20213 carried a total of 4 *β*-galactosidase genes, with 2 belonging to GH2 (*lacZ6* and *lacZ2*) and 2 to GH42 (*lacZ4* and *lacZ5/lntA* D4BMY8; **Figure 6**). This gene profile suggests that *Bi. breve* JCM1192/DSM20213 possesses a comprehensive enzymatic toolkit for the utilization of a large portion of HMOs.

**Figure 6.**
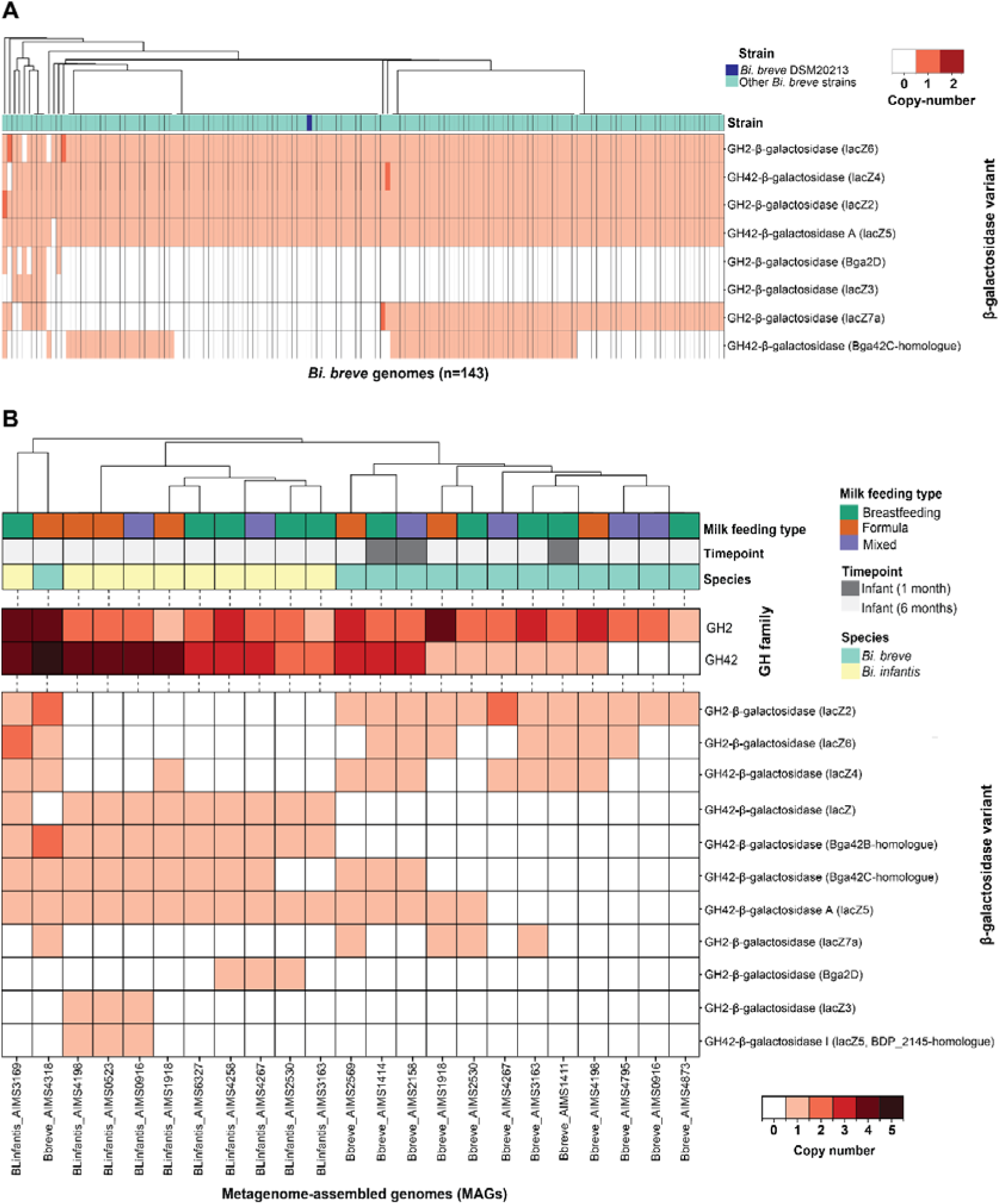
Intra-specific variation in *β*-galactosidase variants reveals different HMO-utilization potential in *Bi. breve*. (A) Heatmap showing the copy-number of genes encoding for different *β*-galactosidase variants across 143 publicly available *Bi. breve* genomes. The annotation bar shows the placement of the focal *Bi. breve* JCM1192/DSM20213 (blue) relative to other *Bi. breve* genomes (light green). Similarly, the heatmap (B) shows the copy-number of genes encoding for *β*-galactosidase variants for *Bi. breve* (light green) and *Bi. infantis* (light yellow). MAGs from AIMS infants (sampling time point: 1-month infant = dark gray, 6-month infant = light gray, milk feeding type: breastfeeding = green, formula = orange, mixed = purple). *β*-galactosidase copy number is shown grouped by GH family (A) and per enzyme variant (B). Genomes are hierarchically clustered based on *β*-galactosidase copy-number (method: average linkage, distance: euclidean)

To investigate the *in vivo* distribution of distinct *β*-galactosidase genes, we retrieved high-quality MAGs of *Bi. breve* and *Bi. infantis* from fecal and oral microbiomes of one- and six-months-old infants of the Amsterdam Infant Microbiome Study (AIMS)^12^. Including *Bi. infantis* MAGs allowed us to place *Bi. breve* variation in an inter-species context, as both species co-occur in the infant gut and share the capacity to utilize HMOs. Examining *β*-galactosidase gene profiles, we detected notable inter- and intra-species variation in the presence and copy numbers of GH2 and GH42 family enzymes (**Figure 6B**), suggesting distinct HMO-utilization potential among infant-associated strains. The *lacZ2* (BBR_RS10300) gene was found across all investigated *Bi. breve* MAGs (**Figure 6B**). The D4BLG2 protein encoded by *lacZ2* was expressed to a similar level in the presence of glucose, LNT or LNnT, suggesting that this protein is constitutively present in *Bi. breve* and may not contribute to the different utilization patterns of LNT and LNnT (**Figure 3**). Notably, *lacZ6* (BBR_RS18475) homologs were present in 61% (8/13) of the infant-derived MAGs (Figure 6B), suggesting that the capacity for *β*-galactosidase activity on LNT and LNnT is variably distributed among infant-associated strains. The enzyme D4BR09 encoded by *lacZ6* is inducibly and highly expressed in LNT and LNnT versus glucose (**Figure 3**), with broad substrate capacity associated with the utilization of HMO-structures such as galactobiose Gal*β*1-4Gal, LNT/LNB and LNnT. Among the GH42 genes, *lntA/lacZ5* (BBR_RS13095), present in *Bi. breve* JCM1192/DSM20213 (D4BMY8) and 46% (6/13) of the MAGs, has been specifically associated with the degradation of the type 1 HMO structures (e.g. LNT) that are predominant in human milk. This aligns with its high protein expression in the presence of LNT or LNnT (**Figure 3**). Compared to *Bi. breve*, *Bi. infantis* MAGs displayed a markedly different *β*-galactosidase gene profile, characterised by low prevalence of *lacZ4* (20%, 2/10) and *lacZ6* (10%, 1/10), but presence of *lacZ5* and *Bga2A* in all investigated MAGs (100%, 10/10; **Figure 6B**). Bga2A has been specifically associated with LNnT and lac degradation^48,49^. Together, the distribution patterns of *β*-galactosidase variants across infant-derived *Bi. breve* and *Bi. infantis* MAGs highlight the divergent HMO-utilisation potential both within and between these two dominant infant gut bifidobacteria.

## Discussion

We established an 8-species infant gut SynCom iBaCo and investigated how individual HMOs influence its composition and metabolic function. We found that neutral HMOs can maintain a higher community diversity and bifidobacterial abundance than fucosyl and sialyl HMOs. In particular, LNT but not LNnT can specifically promote *Bi. breve* in iBaCo. Using proteomics, enzymatic kinetics, and molecular docking, we identified a *Bi. breve* GH42 family *β*-galactosidase LacZ5 D4BMY8 with the key preferential utilization of LNT over LNnT that largely explain the higher abundance of *Bi. breve* in iBaCo. *Bi. breve* harboring lacZ5-encoding gene is divergently distributed in human infant stools. Since the introduction of defined bacterial community in germ-free mice by Russell W. Schaedler in the mid-60s^50^, SynCom has been an important tool to study microbial ecology, microbiota-host interaction, and beyond. To date, much progress has been made on human adult gut microbiota, with over 30 reported SynCom consisting of 2-119 bacteria species^18^, but progress on infant gut microbiota using SynComs is lagging behind. An early effort by Jeffery Gordon’s group^51^ sought to identify determinants of bacterial succession during the early colonization period using SynCom and gnotobiotic mice. They found that the SynCom S2 representing 2-6-month infant outcompeted S1, representing 0-2-month infant, in mice. Among S2, two *Bifidobacterium* species, *Bi. breve* and *Bi. adolescentis* were maintained. Using an infant SynCom of four *Bifidobacterium* species, Ojima et al.^20^ demonstrated the priority effects could shape the structure of the community even in the presence of the same HMO. The strength of the observed priority effects can be strongly enhanced by a mixture of HMOs, i.e. equal mixture of 2’-FL, 3-FL, LNT, LNnT, 3’-SL and 6’-SL^52^. Using individual HMOs, we demonstrated that HMOs have deterministic but divergent effects on the iBaCo at the community and metabolic level. In particular, SL seems to have a strong inhibitory effect on bifidobacteria, while LNT and FL tends to promote bifidobacteria in iBaCo. However, in other infant synthetic communities (e.g. 7-strain infant bacterial community BabyBac and 13-strain BIG-Syc), *Bi. infantis* was outcompeted by other species in the presence of 4HMO or 5HMO mix (2′-FL, 3-FL, LNT, 3′-SL, and 6′-SL) and was undetectable or negligible in the community at the end of cultivation^53^, despite that *Bi. infantis* is an efficient utilizer of these HMOs in monoculture^54–59^. This discrepancy may be due to the different experimental settings such as cultivation models (static tubes vs chemotaxis bioreactor) and HMO exposure (individual vs mixed HMO). Nonetheless, these studies demonstrated the strong influence of HMOs in the bacterial community structure.

Beyond HMO-community level mechanisms, we demonstrated the key role of single enzyme-substrate specificity in orchestrating the abundance of bifidobacteria in an infant gut bacterial community iBaCo. This has important conceptual consequences as it underlines a single gene/enzyme as a strong driving force of microbial ecology and community function. Previously, Foster and co-workers developed a body of ecological theory and predicted that microbial competition but not cooperation increases community stability^60^. Among many factors, utilization of microbiota-accessible dietary compounds is an important driving force of microbial abundance in a community^61^. Our data fit this theory and illustrate that lacZ5-mediated privileged utilization of LNT over LNnT enables *Bi. breve* to maintain its abundance in iBaCo. The small conformational difference (*β*1-3 in LNT versus *β*1-4 in LNnT) strongly contributes to their distinct interaction with lacZ5, which ultimately lead to pronounced differences at *Bi. breve* growth and metabolite level, as well as ecosystem-level changes in iBaCo. Furthermore, the metabolic flux analysis illustrates that lacZ5 kinetics between LNT and LNnT can bias the relative ATP-per-enzyme-cost efficiency of downstream pathways. Nonetheless, this enzyme in *Bi. infantis* (Bga42A, Blon_2016) also possesses substrate preference on LNT over LNnT, but this substrate preference did not lead to its different abundance in iBaCo, possibly due to the presence of more GH42 and GH2 glycan hydrolases ^56,62–65^. Similarly, for another *Bi. breve* strain UCC2003, multiple *β*-galactosidases enable a robust utilization of LNT and LNnT^66,67^, potentially due to the presence of six *β*-galactosidases in its genome while only four in *Bi. breve* JCM1192/DSM20213. The divergent distribution of lacZ5 homologs across infant-derived Bi. breve MAGs indicates a substantial intra-specific variation in *β*-galactosidase repertoires and subsequent HMO-utilization potential in vivo in infants. This suggests that the LNT-utilization strategy observed in *Bi. breve* JCM1192/DSM20213 potentially contributing to the distinct *Bi. breve* colonization patterns observed across infants^68^.

While iBaCo was useful to unravel the key molecular mechanisms, as a simplified infant gut microbiota, it shares the limitations of other SynCom. First, it does not fully capture the complexity of the infant gut microbiota. An infant gut typically harbors up to 50 species per infant at age of 1-6 month^68^. Although this number is much lower than the number of taxa in adult gut microbiota (typically at least 160 species per individual)^69^, it is still challenging to mimic in the laboratory. Cheng et al.^70^ recently demonstrated that it is feasible to establish a highly complex adult gut SynCom hCom2 consisting of 119 gut bacterial species, calling for an effort to construct an infant SynCom of high complexity. Second, iBaCo is dominated by *Ba. fragilis*, followed by *Bifidobacterium*. The dominance of Bacteroides implies that iBaCo is closer to Bacteroidetes-dominant cluster of infants of 12 months age (31.6% of the 405 infants), rather than Firmicutes-dominant (46.0%) or Proteobacteria-dominant cluster (22.4%)^71^. At 2 weeks to 6 months after birth, the dominance of bifidobacteria is even more prominent, accounting for up to 75% of the infants^43^. Future work is warranted to finetune the communities to represent the infant gut microbiota at different life stages and living conditions. In spite of these limitations, our findings provide a concrete example where a single enzyme-level trait drives the species behavior in a community and the community-level function, thus linking trait-based ecological theory with a infant-associated microbiome.

## Supporting information

Supplementary information_iBaCo

## Acknowledgements

We thank J. Bleeker for technical assistance on LC-MS analysis of the samples, S. Brul for helpful discussion and proof reading, F. Branco dos Santos for technical support on HPLC, M. Schwalbe for technical assistance on HPAEC analysis, A. A. Antoniou for visualizing HPAEC chromatographs and growth parameters, and S. Syeda for technical assistance on the plasmid construction. The HMOs were kindly provided by dsm-firmenich via the HMO donation program. The *Bifidobacterium breve* UCC 2003 and its mutants were kindly provided by D. van Sinderen at the University College Cork.

## Authors’ Contribution

YS: Conceptualization, Methodology, Validation, Investigation, Writing, Visualization.

NP: Methodology, Software, Investigation, Formal analysis, Writing, Visualization.

CB: Methodology, Investigation, Writing, Formal analysis, Visualization.

RH: Methodology, Investigation, Formal analysis, Visualization.

MB: Validation, Investigation.

SL: Methodology, Software, Formal analysis, Visualization.

ASC: Methodology, Formal analysis.

GK: Methodology, Resource, Data curation.

WD: Methodology.

ADJvD: Supervision.

DRM: Supervision.

MW: Supervision, Formal analysis, Funding acquisition.

JZ: Conceptualization, Resource, Writing, Visualization, Supervision, Project administration, Funding acquisition.

