## Supplementary information_iBaCo for "A bifidobacterial enzyme orchestrates ecology and function of infant gut bacterial community": Supplementary information_20260417.docx

Supplementary Tables

Supplementary Table 1: Bacteria strains used in this study

| **Bacterial strains** | | |
| --- | --- | --- |
| *Species name* | Feature | Strain designation/source |
| *Bifidobacterium breve* | wild-type strain obtained from DSMZ | JCM1192/DSM 20213 |
| *Bifidobacterium longum subs. infantis* | wild-type strain obtained from DSMZ | DSM 20088 |
| *Bifidobacterium bifidum* | wild-type strain obtained from DSMZ | DSM 20456 |
| *Bifidobacterium pseudocatenulatum* | wild-type strain obtained from DSMZ | DSM 20438 |
| *Bacteroides fragilis* | wild-type strain obtained from DSMZ | DSM 24895 |
| *Blautia luti* | wild-type strain obtained from DSMZ | DSM 14534 |
| *Anaerostipes caccae* | wild-type strain obtained from DSMZ | DSM 14662 |
| *Collinsella aerofaciens* | wild-type strain obtained from DSMZ | DSM 3979 |
| *Bifidobacterium breve* | Isolate from nursling stool | UCC 2003, James et al., (2016), Sci. Rep *6*, 38560 |
| *Bifidobacterium breve* | pORI19-tet-bbr_1552 insertion mutant of UCC2003 | UCC 2003-ΔlacZ6, James et al., (2016), Sci. Rep *6*, 38560 |
| *Bifidobacterium breve* | Tn5-transposon mutant in Bbr_0010 of UCC2003 | UCC2003-ΔlacZ2, James et al., (2016), Sci. Rep *6*, 38560 |
| *Bifidobacterium breve* | pORI19-tet-bbr_0529 insertion mutant of UCC2003 | UCC 2003-ΔlntA , James et al., (2016), Sci. Rep *6*, 38560 |
| *Bifidobacterium breve* | Insertion mutant of UCC2003 | UCC 2003-ΔlacZ4, James et al., (2016), Sci. Rep *6*, 38560 |
| *Escherichia coli* DH5α | Commercial *E. coli* strain for plasmid construction | Molecular Microbial Physiology Group, University of Amsterdam |
| *Escherichia coli* BL21 | Commercial *E. coli* strain for protein expression | Molecular Microbial Physiology Group, University of Amsterdam |
| *E. coli* BL21-pYS01 | BL21 carrying pYS01 | This study |
| *E. coli* BL21-pYS02 | BL21 carrying pYS02 | This study |
| *E. coli* BL21-pYS03 | BL21 carrying pYS03 | This study |

Supplementary Table 2: Consumables used in this study

| Chemical/reagents | Source | Identifier |
| --- | --- | --- |
| **Human Milk Oligosaccharides** | | |
| 2’-FL–2´Fucosyllactose (purity: 98.2%) | dsm-firmenich | CAS#: 41263-94-9 |
| 3-FL – 3-Fucosyllactose (Purity: 92,40 %) | dsm-firmenich | CAS#: 41312-47-4 |
| DFL – Difucosyllactose (Purity: 80,00%) | dsm-firmenich | CAS#: **20768-11-0** |
| 3’SL – 3’-Sialyllactose (Purity: 95,20%) | dsm-firmenich | CAS#: 35890-38-1 |
| 6’SL – 6’sialyllactose sodium salt (Purity: 98,80%) | dsm-firmenich | CAS#: 157574-76-0 |
| Lnt – Lacto-N-Tetraose (Purity: 91,92%) | dsm-firmenich | CAS#: 14116-68-8 |
| LNnt – Lacto-N-Neotetraose (Purity: 99,40%) | dsm-firmenich | CAS#: 3007-32-4 |
| **Chemicals and consumables** | | |
| Casitone | Gibco | Cat#: 225930 |
| Yeast extract | Duchefa Biochemie | CAS#: 8013-01-2 |
| [NaHCO_3_](https://bacmedia.dsmz.de/ingredients/55) | VWR International B.V. | CAS: [6363-53-7](https://www.sigmaaldrich.com/NL/en/product/mm/106329) |
| Glucose | Carl Roth | CAS#: 50-99-7 |
| Maltose | Thermo Fisher Scientific | CAS#: 6365-53-7 |
| Cellobiose | Sigma-Aldrich (Merck) | CAS#: 528-50-7 |
| [L-Cysteine HCl x H_2_O](https://bacmedia.dsmz.de/ingredients/770) | Merck | CAS#: 52-90-4 |
| [Resazurin](https://bacmedia.dsmz.de/ingredients/25) | Thermo Fisher Scientific | CAS#: 62758-13-8 |
| [K_2_HPO_4_](https://bacmedia.dsmz.de/ingredients/10) | Merck | CAS#: 7758-11-4 |
| [KH_2_PO_4_](https://bacmedia.dsmz.de/ingredients/11) | Merck | CAS#: 7778-77-0 |
| [(NH_4_)_2_SO_4_](https://bacmedia.dsmz.de/ingredients/27) | Merck | CAS#: 7783-20-2 |
| [NaCl](https://bacmedia.dsmz.de/ingredients/43) | Thermo Fisher Scientific | CAS#: 7647-14-5 |
| [MgSO_4_ x 7 H_2_O](https://bacmedia.dsmz.de/ingredients/8) | VWR International B. V. | CAS#: 10034-99-8 |
| [CaCl_2_ x 2 H_2_O](https://bacmedia.dsmz.de/ingredients/7) | Merck | CAS#: 10035-04-8 |
| [Acetic acid](https://bacmedia.dsmz.de/ingredients/380) | VWR International B. V. | CAS#: 64-19-7 |
| [Propionic acid](https://bacmedia.dsmz.de/ingredients/381) | VWR International B. V. | CAS#: 79-09-4 |
| [n-Valeric acid](https://bacmedia.dsmz.de/ingredients/288) | Merck | CAS#: 109-52-4 |
| [iso-Valeric acid](https://bacmedia.dsmz.de/ingredients/378) | Merck | CAS#: 503-74-2 |
| [iso-Butyric acid](https://bacmedia.dsmz.de/ingredients/377) | Merck | CAS#: 79-31-2 |
| [KOH](https://bacmedia.dsmz.de/ingredients/126) | VWR International B. V. | CAS#: 1310-58-3 |
| [Ethanol](https://bacmedia.dsmz.de/ingredients/201) | VWR International B. V. | CAS#: 64-17-5 |
| [Hemin](https://bacmedia.dsmz.de/ingredients/704) | Merck | CAS#: 10009-13-5 |
| [Biotin](https://bacmedia.dsmz.de/ingredients/46) | Merck | CAS#: 58-85-5 |
| [Vitamin B12](https://bacmedia.dsmz.de/ingredients/18) | Merck | CAS#: 68-19-9 |
| [p-Aminobenzoic acid](https://bacmedia.dsmz.de/ingredients/47) | Merck | CAS#: 150-13-0 |
| [Folic acid](https://bacmedia.dsmz.de/ingredients/138) | Thermo Fisher Scientific | CAS#: 59-30-3 |
| [Pyridoxine hydrochloride](https://bacmedia.dsmz.de/ingredients/519) | Merck | CAS#: 65-23-6 |
| [Thiamine HCl](https://bacmedia.dsmz.de/ingredients/1207) | Merck | CAS#: 67-03-8 |
| [Riboflavin](https://bacmedia.dsmz.de/ingredients/136) | Merck | CAS#: 83-88-5 |
| DPBS, calcium, magnesium | Thermo Fisher Scientific | Cat#:14040141 |
| Verex vials | Phenomenex | Cat#: AR0-39P0-13 |
| Formic acid | Biosolve | CAS#: 64-18-6 |
| DL-Sodium lactate (50% in aqueous phase) | VWR chemicals | CAS#: 75-17-3 |
| Sodium succinate | Sigma-Aldrich (Merck) | CAS#: 6106-21-4 |
| TLC Silica gel 60 F24 (5 x 7.5 cm) plates | Sigma-Aldrich (Merck) | Cat#: [1.05549](https://www.sigmaaldrich.com/NL/en/product/mm/105549) |
| 1-Butanol | Merck | CAS#: 71-36-3 |
| Sulfuric acid | Merck | CAS#: 7664-93-9 |
| 2 × Rapid Taq Master Mix | Vazyme | Cat#: P222 & 223 |
| Phusion Flash High-Fidelity PCR Master Mix | Thermo Scientific™ | Cat#: F548S |
| DNA purification SPRI Magnetic beads | Applied Biological Materials (abm) | Cat#: G951 |
| Agarose | Thermo Fisher Scientific | Cat#: 16500500 |
| 1 Kb Plus DNA Ladder | Thermo Fisher Scientific | Cat#: 10787018 |
| Midori Green Advance | NIPPON Genetics | Cat#: MG04 |
| Tris | BIO-RAD | Cat#: 610716 |
| Na_2_EDTA | Duchefa Biochemie | Cat#: E0511.0250 |
| Glacial acetic acid | Merck | Cat#: 695092 |
| NEB Blunt/TA ligase Master mix | New England Biology (NEB) | Cat#: M0367 |
| NEBNext FFPE Repair Mix | NEB | Cat#: M6630 |
| NEBNext Ultra II End repair/dA-tailing Module | NEB | Cat#: E7546 |
| NEB Quick Ligation Module | NEB | Cat#: E6056 |
| Flongle Flow Cell | Oxford Nanopore technology | Cat#: FLO-MIN114 |
| MinION Flow Cell | Oxford Nanopore technology | Cat#: FLO-FLG114 |
| cOmplete™ Protease Inhibitor Cocktail | Roche | Cat#: 11873580001 |
| Ammonium bicarbonate (Ambic) | Merck | CAS#: [1066-33-7](https://www.sigmaaldrich.com/NL/en/search/1066-33-7?focus=products&page=1&perpage=30&sort=relevance&term=1066-33-7&type=cas_number) |
| sodium dodecyl sulfate | Merck | CAS#: [151-21-3](https://www.sigmaaldrich.com/NL/en/search/1066-33-7?focus=products&page=1&perpage=30&sort=relevance&term=1066-33-7&type=cas_number) |
| Tris(2-carboxyethyl) phosphine hydrochloride | Sigma-Aldrich (Merck) | Cat#: C4706-2G |
| 2-Chloroacetamide | Sigma-Aldrich (Merck) | Cat#: C0267-100G |
| Ammonium bicarbonate | Sigma-Aldrich (Merck) | Cat#: 11213-1KG-R |
| Sequencing Grade Modified Trypsin | Promega Corporation | Cat#: V5111 |
| Native Barcoding Kit 24 V14 | Oxford nanopore Technologies | Cat#: SQK-NBD114.24 |
| Qubit™ 1X dsDNA High Sensitivity (HS) and Broad Range (BR) Assay Kits | Invitrogen by Thermo Fisher scientific | Cat#: Q33230 |
| PureLink™ Genomic DNA Mini Kit | Thermo Fisher Scientific | Cat#: K182002 |
| Pierce™ BCA Protein Assay Kits | Thermo Fisher Scientific | Cat#: A55864 |
| Monarch^®^ Spin PCR & DNA Cleanup Kit (5 μg) | NEB | Cat#: T1130S |
| Monarch^®^ Spin DNA gel Extraction Kit | NEB | Cat#: T1120S |
| Monarch^®^ Spin Plasmid Miniprep Kit | NEB | Cat#: T1110S |
| Mix & Go! E.coli Transformation Kit | ZYMO RESEARCH | Cat#: T3001 |
| His-Spin Protein Miniprep | ZYMO RESEARCH | Cat#: P2001 |
| Kanamycin Sulphate Monohydrate | Duchefa Biochemie | Cat#: K0126.0001 |
| T4 DNA ligase | NEB | Cat#: M0202S |
| BamHI | Thermo Fisher Scientific | Cat#: ER0055 |
| NdeI | Thermo Fisher Scientific | Cat#: ER0581 |
| [HindIII](https://www.neb.com/en/products/r0104-hindiii) | Thermo Fisher Scientific | Cat#: ER0501 |
| FastDigest Green Buffer (10 ×) | Thermo Fisher Scientific | Cat#: B72 |
| 2x Laemmli Sample Buffer | BIO-RAD | Cat#: 1610737 |
| X-Gal | Duchefa Biochemie | Cas#: 7200-90-6 |
| Dimethyl sulfoxide (DMSO) | Merck | Cat#: 472301 |
| SurePAGE™, Bis-Tris, 10x8, 4-20%, 12 wells | GeneScript | Cat#: M00656 |
| InstantBlue^®^Coomassie Protein Stain | abcam | Cat#: ISB1L |

Supplementary Table 3: Hardwares used in this study

| Hardwares | Manufacturer | Type code |
| --- | --- | --- |
| High-Performance liquid chromatography | Shimadzu Corporation | LC-20AT |
| Refractive index detector | Shimadzu Corporation | RID 20A |
| Ion exclusion Rezex ROA-Organic Acid H+(8%) column (300 × 7.8 mm) | Phenomenex | 00H-0138-KO |
| Nanopore Minion | Oxford Nanopore technology | MIN-101B |
| Qubit™ 4 Fluorometer | Invitrogen by Thermo Fisher scientific | Q33226 |
| Hungate tubes | Chemglass | Cat#: CLS-4208-11 |
| High-performance Aion-Exchange Chromatography (Dionex™ ICS-6000 Standard Bore and Microbore HPIC™ Systems) | Thermo Fisher scientific | ICS-6000 |
| HPAEC guard column (Dionex™ CarboPac™ PA1 IC Columns, 4 × 50 mm) | Thermo Fisher scientific | 43096 |
| HPAEC analytical column (Dionex™ CarboPac™ PA1 IC Columns, 4 × 250 mm) | Thermo Fisher scientific | Cat#: 057178 |
| Plate reader | BYONOY | Absorbance-96 |
| Spectrophotometer | DeNovix | DS-11+ |
| Thermocycler | Bionetra | Biometra Tone 96 |
| Electrophoresis power supply | BIO-RAD | Powerpack 300 |
| Gel Imaging System | NIPPON Genitics | FAS-V |
| Ultrasonicator | Branson | CT 06810, 250 power supply 100-132-134 |
| UHPLC system | Thermo Fisher scientific | Ultimate 3000 RSLCnano |
| C18, 1.6 μm particle size, 75 μm × 250 mm analytical column | Ionopticks | AUR4-25075C18 |
| TIMS-TOF Pro mass spectrometer | Bruker | timsTOF Pro |
| Bead beater | Mmr | HOG-024 |

Supplementary Table 4: Primers used in this study

| Name of primers | Sequence (5’ - 3’) |
| --- | --- |
| 8f | AGTTTGATCCTGGCTCAG |
| 1492r | TACGGYTACCTTGTTACGACTT |
| HE_D4BMY8_f | TCGCATATGGAACATCGCGAATTCAA |
| HE_D4BMY8_r | ACTGGATCCTTACAGCTTTACCACCAG |
| HE_D4BMY8_seq1 | TCAGACAGGATGGTCAG |
| HE_D4BR09_f | TCGCATATGAACACAACCGACGATCA |
| HE_D4BR09_r | ATGAAGCTTCAGATGAGTTCGAGGTTCA |
| HE_D4BR09_seq1 | GCGAAATTAATACGACTCAC |
| HE_D4BR09_seq2 | AACTCAACGCGAGGCCTTCCG |
| HE_D4BR09_seq3 | ATCAGCGATTTCGAGAGCCG |
| HE_D4BR09_seq4 | GCAATCGCTCCGGATACGA |
| HE_D4BMM7_f | TCGCATATGACTACTCGTAGAGCATTT |
| HE_D4BMM7_r | ATGAAGCTTTTAGCAGGACGTTTTAGC |
| HE_D4BMM7_seq1 | CGCGAAATTAATACGACTCAC |
| HE_D4BMM7_seq2 | GACATGCTGCTCGACTTCT |
| HE_D4BMM7_seq3 | ATCGGGCTCGGCGGATACCCAG |

Supplementary Table 5: Plasmids and recombinant proteins used in this study

| Name of Plasmids/protein | Feature | Source |
| --- | --- | --- |
| pYS01 | Km^r^, pET28a plasmid containing BIFBRE_03435 from *Bi. breve* DSM 20213 | This study |
| pYS02 | Km^r^, pET28a plasmid containing BIFBRE_04539 from *Bi. breve* DSM 20213 | This study |
| pYS03 | Km^r^, pET28a plasmid containing BIFBRE_03324 from *Bi. breve* DSM 20213 | This study |
| His₆-tagged recombinant D4BMY8 | Putative GH42 family β-galactosidase, expressed by *E. coli* BL21-pYS01 | This study |
| His₆-tagged recombinant D4BR09 | Putative GH2 family β-galactosidase, expressed by *E. coli* BL21-pYS02 | This study |
| His₆-tagged recombinant D4BMM7 | Putative GH42 family β-galactosidase, expressed by *E. coli* BL21-pYS03 | This study |

Supplementary Table 6: Softwares used in this study

| Software | Source | Application |
| --- | --- | --- |
| GraphPad Prism (V9.0) | https://www.graphpad.com | Statistics and plotting |
| Chromeleon (7.2.10 ES MUj) | https://www.thermofisher.com | HPAEC data analysis |
| Microsoft Excel (Office 16) | https://www.microsoft.com | Data processing |
| Adobe Illustrator 2023 | https://www.adobe.com | Potting |
| EPI2ME (5.2.5) | https://epi2me.nanoporetech.com | Nanopore sequencing data analysis |
| dbCAN3 | https://bcb.unl.edu/dbCAN3 | β-galactosidase genes profiling |
| R (V4.4.0) | https://www.graphpad.com | Growth parameter analysis and plotting |
| R studio(V4.4.0) | https://www.thermofisher.com | Growth parameter analysis and plotting |
| Sanpgene (V6.2.1) | https://snapgene.com | Gene and plasmid visualization |
| BBDuk & BBMap | <https://bbmap.org> | β-galactosidase genes profiling |
| Python (V3.10) | https://www.python.org | Metabolite flux analysis |
| Biorender | https://biorender.com | Schematic plot |
| Pandas | https://pandas.pydata.org | Metabolite flux analysis |
| R package growthrates | https://tpetoldt.github.io/growthrates | Growth parameter analysis |
| Perseus (V2.1.3.0) | https://maxquant.net/perseus | Proteomics analysis |
| PyMol | https://pymol.org | Molecular docking |
| ChatGPT (gpt-5.4) | https://chatgpt.com | Manuscript polishing |
| UvA AI Chat (based on gpt-5.1) | https://aichat.uva.nl/chat | Manuscript polishing |

Supplementary Table 7. The predicted binding affinities and number of hydrogen bonds for each enzyme-ligand interaction.

| **Sample** | **Mean affinity** ± **SD (kcal/mol)** | **Hydrogen bonds with ligand** |
| --- | --- | --- |
| *Bi. infantis* + LNT | −9.5 ± 0.36 | 12 |
| *Bi. infantis* + LNnT | −8.3 ± 0.05 | 4 |
| *Bi. breve* D4BMY8 + LNT | −9.9 ± 0.26 | 6 |
| *Bi. breve* D4BMY8 + LNnT | −8.7 ± 0.12 | 7 |

Supplementary Table 8. Significantly changed protein in LNT versus GMC

| significance | log (P-value) | log 2-fold change | protein IDs | Protein names | Gene Names |
| --- | --- | --- | --- | --- | --- |
| + | 9.190026 | -1.62087 | D4BQ33 | Beta-carotene 15,15'-monooxygenase | BIFBRE_04187 |
| + | 9.412514 | 6.599269 | D4BR10 | Glycoside/pentoside/hexuronide transporter | gph BIFBRE_04540 |
| + | 4.496684 | -3.41162 | D4BNY7 | Efflux ABC transporter, permease protein | BIFBRE_03795 |
| + | 14.69311 | -3.79114 | D4BQW0 | Phosphotransferase system, EIIC | BIFBRE_04488 |
| + | 2.042394 | -2.30278 | D4BQD7 | Efflux ABC transporter, permease protein | BIFBRE_04306 |
| + | 2.943712 | -2.80611 | D4BQ14 | Xylulose kinase (Xylulokinase) (EC 2.7.1.17) | xylB BIFBRE_04242 |
| + | 4.061941 | -1.4668 | D4BQ42 | Fatty acid ABC transporter ATP-binding/permease protein | BIFBRE_04196 |
| + | 7.425742 | 0.694372 | D4BMT8 | Transcriptional regulator, DeoR family | BIFBRE_03385 |
| + | 8.628599 | -1.00794 | D4BQ32 | ABC transporter, permease protein | BIFBRE_04186 |
| + | 5.077841 | -1.14247 | D4BRR7 | Penicillin-binding protein, transpeptidase domain protein | BIFBRE_04807 |
| + | 6.4455 | 2.59016 | D4BS02 | ROK family protein | BIFBRE_04893 |
| + | 2.014071 | -1.54459 | D4BLQ4 | Alpha amylase, catalytic domain protein | BIFBRE_02991 |
| + | 2.575076 | 1.0484 | D4BQC2 | Pyridine nucleotide-disulfide oxidoreductase | BIFBRE_04291 |
| + | 3.76041 | 1.3015 | D4BR08 | Transcriptional regulator, LacI family | BIFBRE_04538 |
| + | 6.265045 | 1.620415 | D4BMY8 | Beta-galactosidase (Beta-gal) (EC 3.2.1.23) | BIFBRE_03435 |
| + | 2.388814 | 0.813058 | D4BMN6 | Phosphotyrosine-protein phosphatase family protein | BIFBRE_03333 |
| + | 2.856604 | 0.780695 | D4BNI9 | Thiazole synthase (EC 2.8.1.10) | thiG BIFBRE_03639 |
| + | 1.976408 | -3.02523 | D4BQD6 | ABC transporter, ATP-binding protein | BIFBRE_04305 |
| + | 5.433454 | 2.19372 | D4BMY3 | Sugar-binding domain protein | BIFBRE_03431 |
| + | 7.053148 | 0.892626 | D4BNQ8 | Pyruvate kinase (EC 2.7.1.40) | pyk BIFBRE_03709 |
| + | 13.31358 | 2.412152 | D4BQW7 | 1,3-beta-galactosyl-N-acetylhexosamine phosphorylase (EC 2.4.1.211) | gnpA BIFBRE_04496 |
| + | 6.359408 | 2.849343 | D4BMY6;D4BMM6;D4BME5 | ABC transporter, permease protein | BIFBRE_03433, BIFBRE_03323, BIFBRE_03239 |
| + | 3.140313 | -0.58128 | D4BQ30 | Transcriptional regulator, LacI family | BIFBRE_04184 |
| + | 8.387686 | 2.469437 | D4BMY5 | ABC transporter, permease protein | BIFBRE_03432 |
| + | 15.24226 | -2.55596 | D4BLY3 | Phosphoenolpyruvate-protein phosphotransferase (EC 2.7.3.9) (Phosphotransferase system, enzyme I) | ptsP BIFBRE_03070 |
| + | 3.276358 | -1.14291 | D4BNZ2 | ABC transporter, substrate-binding protein, family 3 | BIFBRE_03800 |
| + | 5.009064 | 2.299838 | D4BL96 | Putative glucose-6-phosphate 1-epimerase (EC 5.1.3.15) | BIFBRE_02828 |
| + | 4.677898 | -1.2794 | D4BQ41 | ABC transporter, ATP-binding protein | BIFBRE_04195 |
| + | 3.412467 | -0.92331 | D4BM68 | Copper-exporting ATPase (EC 3.6.3.4) | BIFBRE_03160 |
| + | 10.54474 | 4.366953 | D4BL97 | Galactose-1-phosphate uridylyltransferase (Gal-1-P uridylyltransferase) (EC 2.7.7.12) (UDP-glucose--hexose-1-phosphate uridylyltransferase) | galT BIFBRE_02829 |
| + | 5.379602 | -0.89703 | D4BR97 | Fatty acid ABC transporter ATP-binding/permease protein | BIFBRE_04633 |
| + | 3.004781 | 4.892037 | D4BMM4 | ABC transporter, solute-binding protein | BIFBRE_03321 |
| + | 2.103641 | -0.94196 | D4BQW1 | PRD domain protein | BIFBRE_04489 |
| + | 1.950652 | 1.231227 | D4BN85 | GroES-like protein | BIFBRE_03534 |
| + | 2.398008 | 1.297612 | D4BQN6 | Glycosyl hydrolase, family 1 | BIFBRE_04413 |
| + | 2.791101 | -0.98396 | D4BLD1 | Thioredoxin-like fold domain-containing protein | BIFBRE_02863 |
| + | 6.925427 | 1.702578 | D4BQX0 | UDP-glucose 4-epimerase (EC 5.1.3.2) | galE BIFBRE_04499 |
| + | 11.893 | 2.602228 | D4BQW6 | ABC transporter, permease protein | BIFBRE_04495 |
| + | 2.152874 | 1.448602 | D4BQ70 | ErfK/YbiS/YcfS/YnhG | BIFBRE_04225 |
| + | 2.295071 | 0.95601 | D4BSE6 | protein adenylyltransferase (EC 2.7.7.108) | BIFBRE_05038 |
| + | 2.414308 | -1.40946 | D4BM51 | Export protein | BIFBRE_03143 |
| + | 2.236373 | -1.57671 | D4BM52 | ABC transporter, ATP-binding protein | BIFBRE_03144 |
| + | 2.553113 | 0.746065 | D4BQ27 | Phosphoglycerate mutase family protein | BIFBRE_04181 |
| + | 2.10687 | 1.252719 | D4BL77 | ABC transporter, solute-binding protein | BIFBRE_02810 |
| + | 5.142692 | 2.765847 | D4BMY9 | ABC transporter, solute-binding protein | BIFBRE_03436 |
| + | 2.052151 | 1.094611 | D4BLA5 | Sugar-binding domain protein | BIFBRE_02837 |
| + | 4.557308 | -1.22727 | D4BLR1 | 4-alpha-glucanotransferase (EC 2.4.1.25) (Amylomaltase) (Disproportionating enzyme) | malQ BIFBRE_02998 |
| + | 7.967366 | 2.838959 | D4BQW9 | Phosphotransferase enzyme family | BIFBRE_04498 |
| + | 7.932428 | -2.30536 | D4BLY0 | Transcriptional regulator, LacI family | BIFBRE_03067 |
| + | 4.484847 | -0.5701 | D4BLI9 | Phosphoenolpyruvate carboxylase | ppc BIFBRE_02924 |
| + | 3.121723 | -1.67595 | D4BLC9 | Glycosyl hydrolase family 3 N-terminal domain protein | BIFBRE_02861 |
| + | 3.141365 | -1.17668 | D4BLQ5 | Transcriptional regulator, LacI family | BIFBRE_02992 |
| + | 3.944406 | -3.3322 | D4BNY6 | ABC transporter, ATP-binding protein | BIFBRE_03794 |
| + | 2.846286 | 0.974486 | D4BR15 | Lactaldehyde reductase (EC 1.1.1.77) | fucO BIFBRE_04548 |
| + | 5.448004 | 1.035984 | D4BMU0 | Galactokinase (EC 2.7.1.6) | galK BIFBRE_03387 |
| + | 2.880496 | -0.70256 | D4BRI4 | Polyphosphate:nucleotide phosphotransferase, PPK2 family (EC 2.7.4.-) | BIFBRE_04720 |
| + | 2.910597 | -1.19121 | D4BLR2 | Alpha amylase, catalytic domain protein | BIFBRE_02999 |
| + | 11.04567 | 7.940842 | D4BS05 | N-acetylglucosamine-6-phosphate deacetylase (EC 3.5.1.25) | nagA BIFBRE_04896 |
| + | 5.334621 | 2.261845 | D4BS03 | ROK family protein | BIFBRE_04894 |
| + | 5.526592 | -0.68031 | D4BR24 | ATP-dependent zinc metalloprotease FtsH (EC 3.4.24.-) | hflB ftsH BIFBRE_04557 |
| + | 2.531384 | -1.63325 | D4BQX3 | Uncharacterized protein | BIFBRE_04502 |
| + | 3.702533 | -0.93379 | D4BPK8 | Multifunctional fusion protein [Includes: Cytidylate kinase (CK) (EC 2.7.4.25) (Cytidine monophosphate kinase) (CMP kinase); GTPase Der (GTP-binding protein EngA)] | der cmk BIFBRE_04022 |
| + | 11.50952 | 4.099134 | D4BS04 | Glucosamine-6-phosphate deaminase (EC 3.5.99.6) (GlcN6P deaminase) (GNPDA) (Glucosamine-6-phosphate isomerase) | nagB BIFBRE_04895 |
| + | 3.478085 | 0.622911 | D4BNW1 | Formate acetyltransferase (EC 2.3.1.54) (Pyruvate formate-lyase) | pflB BIFBRE_03767 |
| + | 3.836755 | -1.20288 | D4BMU7 | Adenine DNA glycosylase (EC 3.2.2.31) | BIFBRE_03394 |
| + | 2.145109 | -1.21117 | D4BQ15 | Sugar-binding domain protein | BIFBRE_04243 |
| + | 2.97616 | 0.762422 | D4BLA6 | ABC transporter, ATP-binding protein | BIFBRE_02838 |
| + | 2.388641 | -0.85999 | D4BPK7 | Pseudouridine synthase (EC 5.4.99.-) | BIFBRE_04021 |
| + | 2.513775 | 1.42626 | D4BL72 | Alpha amylase, catalytic domain protein | BIFBRE_02804 |
| + | 2.898365 | -0.74763 | D4BR64 | HdeD family acid-resistance protein | BIFBRE_04598 |
| + | 2.178203 | 1.693359 | D4BM96 | Chorismate mutase (EC 5.4.99.5) | BIFBRE_03188 |
| + | 2.102194 | -0.93512 | D4BMX6 | ABC transporter, substrate-binding protein, family 3 | BIFBRE_03423 |
| + | 10.64426 | 7.575546 | D4BR09 | Beta-galactosidase (EC 3.2.1.23) (Lactase) | BIFBRE_04539 |
| + | 5.90541 | 1.097814 | D4BMT9 | Galactose-1-phosphate uridylyltransferase (EC 2.7.7.12) | galT BIFBRE_03386 |
| + | 4.381601 | -0.69557 | D4BR94 | Exo-alpha-(1->6)-L-arabinopyranosidase | BIFBRE_04630 |
| + | 6.64734 | 1.28851 | D4BLB1 | ABC transporter, ATP-binding protein | BIFBRE_02843 |
| + | 7.765561 | 3.180457 | D4BQW5 | ABC transporter, permease protein | BIFBRE_04494 |
| + | 4.778406 | -0.80551 | D4BR96 | ABC transporter, ATP-binding protein | BIFBRE_04632 |
| + | 7.736379 | -4.26133 | D4BR88 | beta-glucosidase (EC 3.2.1.21) | BIFBRE_04624 |
| + | 2.826952 | 0.629173 | D4BP86 | L,D-TPase catalytic domain-containing protein | BIFBRE_03894 |
| + | 1.957344 | -1.38415 | D4BQ19 | Putative 6-phospho 3-hexuloisomerase | BIFBRE_04247 |
| + | 5.295478 | 2.817584 | D4BQW4 | ABC transporter, solute-binding protein | BIFBRE_04493 |

- Pink indicate the proteins significantly over expressed in LNT.

Supplementary Table 9. Significantly changed protein in LNnT versus GMC

| significance | log (P-value) | log 2-fold change | protein IDs | Protein names | Gene Names |
| --- | --- | --- | --- | --- | --- |
| + | 2.7 | -0.7 | D4BS85 | DNA topoisomerase (ATP-hydrolyzing) (EC 5.6.2.2) | BIFBRE_04977 |
| + | 6.1 | 3.9 | D4BMY9 | ABC transporter, solute-binding protein | BIFBRE_03436 |
| + | 1.6 | -1.1 | D4BNZ5 | ABC transporter, permease protein | BIFBRE_03803 |
| + | 1.8 | 1.7 | D4BLB5 | PTS system, Lactose/Cellobiose specific IIB subunit | BIFBRE_02847 |
| + | 2.7 | -1.1 | D4BRC4 | HAD hydrolase, family IA, variant 3 | BIFBRE_04660 |
| + | 1.9 | -1.1 | D4BQX1 | Transcriptional regulator, LuxR family | BIFBRE_04500 |
| + | 3.6 | -0.6 | D4BQB0 | Transcriptional regulator, IclR family, C-terminal domain protein | BIFBRE_04279 |
| + | 5.2 | -1.6 | D4BQ41 | ABC transporter, ATP-binding protein | BIFBRE_04195 |
| + | 2.9 | -0.6 | D4BQK6 | Putative permease | BIFBRE_04379 |
| + | 1.7 | -0.9 | D4BS19 | 4-hydroxy-3-methylbut-2-enyl diphosphate reductase (HMBPP reductase) (EC 1.17.7.4) | ispH BIFBRE_04910 |
| + | 3.9 | -1.4 | D4BR83 | Anaerobic ribonucleoside-triphosphate reductase (EC 1.17.4.2) | nrdD BIFBRE_04619 |
| + | 1.8 | 0.8 | D4BP74 | Phosphoglycerate mutase family protein | BIFBRE_03882 |
| + | 6.0 | -1.2 | D4BRH7 | ABC-type polar-amino-acid transporter (EC 7.4.2.1) | BIFBRE_04713 |
| + | 6.1 | -1.8 | D4BNZ2 | ABC transporter, substrate-binding protein, family 3 | BIFBRE_03800 |
| + | 6.2 | -4.1 | D4BNY7 | Efflux ABC transporter, permease protein | BIFBRE_03795 |
| + | 2.3 | 1.2 | D4BQZ0 | NlpC/P60 family protein | BIFBRE_04520 |
| + | 3.5 | 1.4 | D4BPI9 | Fructosamine kinase | BIFBRE_03998 |
| + | 4.5 | -1.1 | D4BMX1 | Peptide chain release factor 2 (RF-2) | prfB BIFBRE_03418 |
| + | 3.4 | 1.4 | D4BQM6 | DNA polymerase IV (Pol IV) (EC 2.7.7.7) | dinB BIFBRE_04402 |
| + | 3.8 | 1.0 | D4BLY7 | Raf-like protein | BIFBRE_03074 |
| + | 1.4 | 1.2 | D4BLV2 | UDP-N-acetylglucosamine diphosphorylase | BIFBRE_03039 |
| + | 2.9 | 0.8 | D4BMU0 | Galactokinase (EC 2.7.1.6) | galK BIFBRE_03387 |
| + | 3.2 | 1.2 | D4BR08 | Transcriptional regulator, LacI family | BIFBRE_04538 |
| + | 3.5 | -1.9 | D4BMZ8 | Putative cystathionine beta-synthase | BIFBRE_03445 |
| + | 2.7 | 0.5 | D4BLD0 | non-specific serine/threonine protein kinase (EC 2.7.11.1) | BIFBRE_02862 |
| + | 3.7 | 2.1 | D4BN85 | GroES-like protein | BIFBRE_03534 |
| + | 1.3 | 1.5 | D4BN09 | Pyridoxal 5'-phosphate synthase subunit PdxS (PLP synthase subunit PdxS) (EC 4.3.3.6) (Pdx1) | pdxS BIFBRE_03456 |
| + | 1.5 | 0.7 | D4BLJ0 | Threonine/serine exporter-like N-terminal domain-containing protein | BIFBRE_02925 |
| + | 4.7 | 1.0 | D4BLV0 | Aspartate-semialdehyde dehydrogenase (EC 1.2.1.11) | asd BIFBRE_03037 |
| + | 3.2 | 1.1 | D4BR05 | ABC transporter, solute-binding protein | BIFBRE_04535 |
| + | 2.7 | 1.1 | D4BMS2 | G5 domain protein | BIFBRE_03369 |
| + | 1.6 | 1.3 | D4BQ25 | Nitroreductase family protein | BIFBRE_04179 |
| + | 4.9 | 0.6 | D4BNK5 | NAD kinase (EC 2.7.1.23) (ATP-dependent NAD kinase) | nadK BIFBRE_03655 |
| + | 1.8 | -0.6 | D4BRR5 | Ribosomal RNA small subunit methyltransferase H (EC 2.1.1.199) (16S rRNA m(4)C1402 methyltransferase) (rRNA (cytosine-N(4)-)-methyltransferase RsmH) | mraW rsmH BIFBRE_04805 |
| + | 2.1 | -0.7 | D4BNB3 | ErfK/YbiS/YcfS/YnhG | BIFBRE_03562 |
| + | 1.7 | 0.6 | D4BLP8 | Transporter, major facilitator family protein | BIFBRE_02985 |
| + | 4.6 | 0.8 | D4BRP6 | Uncharacterized protein | BIFBRE_04786 |
| + | 3.2 | 0.5 | D4BPC7 | Glycogen synthase, Corynebacterium family | glgA BIFBRE_03936 |
| + | 2.1 | 0.5 | D4BPK4 | Bifunctional purine biosynthesis protein PurH [Includes: Phosphoribosylaminoimidazolecarboxamide formyltransferase (EC 2.1.2.3) (AICAR transformylase); IMP cyclohydrolase (EC 3.5.4.10) (ATIC) (IMP synthase) (Inosinicase)] | purH BIFBRE_04018 |
| + | 1.7 | -1.0 | D4BLI6 | Diguanylate cyclase (GGDEF) domain protein | BIFBRE_02921 |
| + | 3.4 | 1.5 | D4BMX4 | CHAP domain protein | BIFBRE_03421 |
| + | 1.7 | 0.9 | D4BNY5 | O-acetylhomoserine aminocarboxypropyltransferase/cysteine synthase | BIFBRE_03793 |
| + | 3.8 | -1.0 | D4BRL2 | ATPase, AAA family | BIFBRE_04747 |
| + | 8.4 | 2.6 | D4BQW6 | ABC transporter, permease protein | BIFBRE_04495 |
| + | 3.0 | 0.9 | D4BN18 | ABC transporter, solute-binding protein | BIFBRE_03466 |
| + | 1.5 | 1.8 | D4BRJ6 | ECF transporter S component | BIFBRE_04732 |
| + | 2.1 | -0.5 | D4BN00 | DNA helicase RecQ (EC 5.6.2.4) | recQ BIFBRE_03447 |
| + | 6.7 | 5.4 | D4BNP7 | 1,4-alpha-glucan branching enzyme GlgB (EC 2.4.1.18) (1,4-alpha-D-glucan:1,4-alpha-D-glucan 6-glucosyl-transferase) (Alpha-(1->4)-glucan branching enzyme) (Glycogen branching enzyme) (BE) | glgB BIFBRE_03698 |
| + | 6.8 | 9.3 | D4BMM4 | ABC transporter, solute-binding protein | BIFBRE_03321 |
| + | 3.0 | 1.3 | D4BP67 | Peptidase C26 | BIFBRE_03875 |
| + | 4.5 | -5.4 | D4BNY6 | ABC transporter, ATP-binding protein | BIFBRE_03794 |
| + | 1.9 | 1.2 | D4BMT2 | 3-isopropylmalate dehydrogenase (EC 1.1.1.85) (3-IPM-DH) (Beta-IPM dehydrogenase) (IMDH) | leuB BIFBRE_03379 |
| + | 3.0 | 4.3 | D4BMF7 | LPXTG-motif cell wall anchor domain protein | BIFBRE_03252 |
| + | 3.8 | -0.9 | D4BN46 | Arylsulfatase (EC 3.1.6.-) | BIFBRE_03494 |
| + | 1.9 | 2.6 | D4BPT9 | Late embryogenesis abundant protein | BIFBRE_04104 |
| + | 2.7 | 1.5 | D4BL93 | PTS system, N-acetylglucosamine-specific IIBC component (EC 2.7.1.69) | nagE BIFBRE_02825 |
| + | 3.5 | -0.7 | D4BQZ3 | Thymidylate synthase (TS) (TSase) (EC 2.1.1.45) | thyA BIFBRE_04523 |
| + | 3.8 | 0.5 | D4BS56 | TRAM domain protein | BIFBRE_04947 |
| + | 1.6 | -1.1 | D4BMN5 | Exopolysaccharide biosynthesis polyprenyl glycosylphosphotransferase | BIFBRE_03332 |
| + | 8.7 | 8.4 | D4BS05 | N-acetylglucosamine-6-phosphate deacetylase (EC 3.5.1.25) | nagA BIFBRE_04896 |
| + | 2.0 | 0.7 | D4BLD2 | G5 domain protein | BIFBRE_02865 |
| + | 4.4 | 0.5 | D4BMT8 | Transcriptional regulator, DeoR family | BIFBRE_03385 |
| + | 5.5 | 1.1 | D4BNC0 | Orotidine-5'-phosphate decarboxylase (EC 4.1.1.23) | pyrF BIFBRE_03569 |
| + | 3.6 | -0.6 | D4BRF1 | ATPase family associated with various cellular activities (AAA) | BIFBRE_04687 |
| + | 3.6 | -0.4 | D4BMH6 | Signal recognition particle protein (EC 3.6.5.4) (Fifty-four homolog) | ffh BIFBRE_03271 |
| + | 2.2 | 0.8 | D4BNV8 | Amidohydrolase | BIFBRE_03764 |
| + | 1.9 | 0.6 | D4BMG5 | LemA family protein | BIFBRE_03260 |
| + | 3.6 | -0.4 | D4BRJ7 | Glycosyltransferase, group 2 family protein (EC 2.4.-.-) | BIFBRE_04733 |
| + | 3.0 | 0.4 | D4BLK1 | Alpha-1,4 glucan phosphorylase (EC 2.4.1.1) | glgP BIFBRE_02936 |
| + | 1.6 | 0.9 | D4BMC4 | Putative endoribonuclease L-PSP | BIFBRE_03218 |
| + | 4.1 | 0.8 | D4BPN1 | Multifunctional fusion protein [Includes: Indole-3-glycerol phosphate synthase (IGPS) (EC 4.1.1.48); Tryptophan synthase beta chain (EC 4.2.1.20)] | trpB trpC BIFBRE_04046 |
| + | 1.8 | 0.6 | D4BLC1 | Ribonucleoside-diphosphate reductase (EC 1.17.4.1) | BIFBRE_02853 |
| + | 5.7 | -1.4 | D4BPK8 | Multifunctional fusion protein [Includes: Cytidylate kinase (CK) (EC 2.7.4.25) (Cytidine monophosphate kinase) (CMP kinase); GTPase Der (GTP-binding protein EngA)] | der cmk BIFBRE_04022 |
| + | 1.4 | -0.8 | D4BNX3 | Na(+)/H(+) antiporter NhaA (Sodium/proton antiporter NhaA) | nhaA BIFBRE_03781 |
| + | 3.0 | -0.6 | D4BLN5 | ferredoxin--NADP(+) reductase (EC 1.18.1.2) | BIFBRE_02971 |
| + | 2.5 | -1.3 | D4BQ37 | Amino acid permease | BIFBRE_04191 |
| + | 2.8 | 0.5 | D4BQD3 | Nucleoside triphosphate pyrophosphatase (EC 3.6.1.9) (Nucleotide pyrophosphatase) (Nucleotide PPase) | maf BIFBRE_04302 |
| + | 1.7 | 0.9 | D4BQ23 | YhgE/Pip domain protein | BIFBRE_04177 |
| + | 2.3 | -0.5 | D4BLZ2 | GtrA-like protein | BIFBRE_03079 |
| + | 5.8 | -0.8 | D4BQE9 | Ribonuclease E (EC 3.1.26.12) | BIFBRE_04317 |
| + | 2.1 | -0.7 | D4BQU8 | tRNA pseudouridine synthase B (EC 5.4.99.25) (tRNA pseudouridine(55) synthase) (Psi55 synthase) (tRNA pseudouridylate synthase) (tRNA-uridine isomerase) | truB BIFBRE_04476 |
| + | 4.9 | 1.5 | D4BR15 | Lactaldehyde reductase (EC 1.1.1.77) | fucO BIFBRE_04548 |
| + | 2.6 | 1.0 | D4BQL8 | Magnesium transporter | BIFBRE_04391 |
| + | 2.7 | -0.7 | D4BNV0 | Bifunctional protein GlmU [Includes: UDP-N-acetylglucosamine pyrophosphorylase (EC 2.7.7.23) (N-acetylglucosamine-1-phosphate uridyltransferase); Glucosamine-1-phosphate N-acetyltransferase (EC 2.3.1.157)] | glmU BIFBRE_03754 |
| + | 4.8 | -1.4 | D4BM27 | Ammonium transporter | amt BIFBRE_03118 |
| + | 6.3 | 1.0 | D4BMR4 | AAA domain-containing protein | BIFBRE_03361 |
| + | 2.3 | 0.8 | D4BPG9 | Aminotransferase, class I/II (EC 2.6.1.-) | BIFBRE_03979 |
| + | 3.5 | -0.5 | D4BR82 | Exodeoxyribonuclease 7 large subunit (EC 3.1.11.6) (Exodeoxyribonuclease VII large subunit) (Exonuclease VII large subunit) | xseA BIFBRE_04618 |
| + | 2.8 | 1.1 | D4BSE6 | protein adenylyltransferase (EC 2.7.7.108) | BIFBRE_05038 |
| + | 4.6 | 1.8 | D4BQ70 | ErfK/YbiS/YcfS/YnhG | BIFBRE_04225 |
| + | 3.7 | -1.3 | D4BPK7 | Pseudouridine synthase (EC 5.4.99.-) | BIFBRE_04021 |
| + | 2.2 | 1.2 | D4BQC2 | Pyridine nucleotide-disulfide oxidoreductase | BIFBRE_04291 |
| + | 2.2 | 0.8 | D4BLM8 | Helicase C-terminal domain protein | BIFBRE_02963 |
| + | 2.7 | -1.3 | D4BQW1 | PRD domain protein | BIFBRE_04489 |
| + | 3.6 | 1.0 | D4BMJ2 | Cell division protein | BIFBRE_03287 |
| + | 8.3 | -2.3 | D4BLY3 | Phosphoenolpyruvate-protein phosphotransferase (EC 2.7.3.9) (Phosphotransferase system, enzyme I) | ptsP BIFBRE_03070 |
| + | 3.6 | 4.6 | D4BMM7 | Beta-galactosidase (Beta-gal) (EC 3.2.1.23) | BIFBRE_03324 |
| + | 1.6 | -1.0 | D4BP06 | Inner membrane component domain-containing protein | BIFBRE_03814 |
| + | 2.6 | 0.4 | D4BMZ4 | ABC transporter, permease protein | BIFBRE_03441 |
| + | 2.1 | 0.6 | D4BN05 | Putative dGTPase (EC 3.1.5.1) | BIFBRE_03452 |
| + | 1.6 | 0.7 | D4BN67 | Formate-dependent phosphoribosylglycinamide formyltransferase (EC 6.3.1.21) (5'-phosphoribosylglycinamide transformylase 2) (Formate-dependent GAR transformylase) (GAR transformylase 2) (GART 2) (Non-folate glycinamide ribonucleotide transformylase) (Phosphoribosylglycinamide formyltransferase 2) | purT BIFBRE_03515 |
| + | 7.5 | -1.3 | D4BR97 | Fatty acid ABC transporter ATP-binding/permease protein | BIFBRE_04633 |
| + | 2.7 | -0.6 | D4BNP8 | CarD-like protein | BIFBRE_03699 |
| + | 3.5 | -0.7 | D4BS24 | Translation initiation factor IF-3 | infC BIFBRE_04915 |
| + | 2.0 | 0.7 | D4BNI9 | Thiazole synthase (EC 2.8.1.10) | thiG BIFBRE_03639 |
| + | 1.4 | 1.0 | D4BLZ1 | Macrophage migration inhibitory factor (MIF) | BIFBRE_03078 |
| + | 7.5 | 3.3 | D4BQW5 | ABC transporter, permease protein | BIFBRE_04494 |
| + | 3.2 | -0.8 | D4BQ68 | Cell envelope-like function transcriptional attenuator common domain protein | BIFBRE_04223 |
| + | 5.9 | -4.3 | D4BR88 | beta-glucosidase (EC 3.2.1.21) | BIFBRE_04624 |
| + | 2.1 | 0.6 | D4BP86 | L,D-TPase catalytic domain-containing protein | BIFBRE_03894 |
| + | 2.5 | -0.6 | D4BPI8 | Chaperone protein DnaJ | dnaJ BIFBRE_03997 |
| + | 5.0 | -2.0 | D4BNV6 | ABC transporter, ATP-binding protein | BIFBRE_03762 |
| + | 3.8 | 1.1 | D4BRM8 | Oleate hydratase | BIFBRE_04764 |
| + | 2.6 | -1.2 | D4BLD1 | Thioredoxin-like fold domain-containing protein | BIFBRE_02863 |
| + | 6.7 | -0.8 | D4BNG5 | DUF3043 domain-containing protein | BIFBRE_03615 |
| + | 2.3 | -1.6 | D4BNY8 | Response regulator receiver domain protein | BIFBRE_03796 |
| + | 1.9 | 0.7 | D4BMJ8 | Nicotinate phosphoribosyltransferase (EC 6.3.4.21) | pncB BIFBRE_03293 |
| + | 2.2 | 1.4 | D4BQ91 | FAH family protein | BIFBRE_04260 |
| + | 1.4 | -1.3 | D4BPE3 | tRNA (adenine(58)-N(1))-methyltransferase TrmI (EC 2.1.1.220) | BIFBRE_03952 |
| + | 2.2 | 0.8 | D4BQN2 | Transcriptional regulator, LacI family | BIFBRE_04409 |
| + | 4.9 | 1.2 | D4BRP7 | Imidazole glycerol phosphate synthase subunit HisH (EC 4.3.2.10) (IGP synthase glutaminase subunit) (EC 3.5.1.2) (IGP synthase subunit HisH) (ImGP synthase subunit HisH) (IGPS subunit HisH) | hisH BIFBRE_04787 |
| + | 1.5 | 1.4 | D4BNC1 | Guanylate kinase (EC 2.7.4.8) (GMP kinase) | gmk BIFBRE_03570 |
| + | 2.9 | -1.5 | D4BNV5 | NLPA lipoprotein | BIFBRE_03761 |
| + | 1.9 | -0.5 | D4BLQ3 | Ketol-acid reductoisomerase (NADP(+)) (KARI) (EC 1.1.1.86) (Acetohydroxy-acid isomeroreductase) (AHIR) (Alpha-keto-beta-hydroxylacyl reductoisomerase) | ilvC BIFBRE_02990 |
| + | 3.1 | -0.9 | D4BLW1 | ABC transporter, ATP-binding protein | BIFBRE_03048 |
| + | 3.7 | -2.1 | D4BQX3 | Uncharacterized protein | BIFBRE_04502 |
| + | 1.7 | 0.8 | D4BPG1 | ADP-dependent (S)-NAD(P)H-hydrate dehydratase (EC 4.2.1.136) (ADP-dependent NAD(P)HX dehydratase) | nnrD BIFBRE_03970 |
| + | 2.5 | 2.0 | D4BS02 | ROK family protein | BIFBRE_04893 |
| + | 8.8 | 8.5 | D4BR09 | Beta-galactosidase (EC 3.2.1.23) (Lactase) | BIFBRE_04539 |
| + | 2.9 | 1.7 | D4BLZ3 | Transporter, major facilitator family protein | BIFBRE_03080 |
| + | 2.4 | 0.7 | D4BNQ2 | ABC transporter, substrate-binding protein | BIFBRE_03703 |
| + | 6.5 | 6.1 | D4BR10 | Glycoside/pentoside/hexuronide transporter | gph BIFBRE_04540 |
| + | 3.5 | 0.7 | D4BLA6 | ABC transporter, ATP-binding protein | BIFBRE_02838 |
| + | 1.8 | 0.9 | D4BNU2 | Uncharacterized protein | BIFBRE_03746 |
| + | 3.7 | 2.2 | D4BMA5 | Alpha-1,4-glucan:maltose-1-phosphate maltosyltransferase (GMPMT) (EC 2.4.99.16) ((1->4)-alpha-D-glucan:maltose-1-phosphate alpha-D-maltosyltransferase) | glgE BIFBRE_03197 |
| + | 1.7 | 0.8 | D4BRJ9 | ROK family protein | BIFBRE_04735 |
| + | 2.2 | -0.7 | D4BRF0 | DUF58 domain-containing protein | BIFBRE_04686 |
| + | 2.2 | 0.5 | D4BR21 | D-alanyl-D-alanine carboxypeptidase/D-alanyl-D-alanine-endopeptidase (EC 3.4.16.4) | dacB BIFBRE_04554 |
| + | 2.9 | -0.5 | D4BS73 | Protein RecA (Recombinase A) | recA BIFBRE_04964 |
| + | 3.4 | -1.9 | D4BLC9 | Glycosyl hydrolase family 3 N-terminal domain protein | BIFBRE_02861 |
| + | 1.6 | 0.8 | D4BMC2 | Adenylate cyclase | BIFBRE_03215 |
| + | 3.1 | 1.2 | D4BQ27 | Phosphoglycerate mutase family protein | BIFBRE_04181 |
| + | 2.8 | 1.6 | D4BM36 | Poly(Hydroxyalcanoate) granule associated protein (Phasin) | BIFBRE_03127 |
| + | 7.5 | 4.1 | D4BRL3 | Multifunctional fusion protein [Includes: Phosphomethylpyrimidine synthase (EC 4.1.99.17) (Hydroxymethylpyrimidine phosphate synthase) (Thiamine biosynthesis protein ThiC) (HMP-P synthase) (HMP-phosphate synthase) (HMPP synthase); Thiamine-phosphate synthase (TP synthase) (TPS) (EC 2.5.1.3) (Thiamine-phosphate pyrophosphorylase) (TMP pyrophosphorylase) (TMP-PPase)] | thiC thiE BIFBRE_04749 |
| + | 4.1 | 1.8 | D4BNG7 | Cobalt ABC transporter, permease protein | BIFBRE_03617 |
| + | 3.7 | 1.7 | D4BP40 | Alpha amylase, catalytic domain protein | BIFBRE_03848 |
| + | 1.4 | 2.1 | D4BMW3 | 4-hydroxy-tetrahydrodipicolinate synthase (HTPA synthase) (EC 4.3.3.7) | dapA BIFBRE_03410 |
| + | 3.1 | -1.0 | D4BLP3 | Fluoride-specific ion channel FluC | fluC crcB BIFBRE_02980 |
| + | 1.6 | 0.8 | D4BMK0 | dITP/XTP pyrophosphatase (EC 3.6.1.66) (Non-canonical purine NTP pyrophosphatase) (Non-standard purine NTP pyrophosphatase) (Nucleoside-triphosphate diphosphatase) (Nucleoside-triphosphate pyrophosphatase) (NTPase) | BIFBRE_03295 |
| + | 3.3 | -0.5 | D4BS12 | BioF2-like acetyltransferase domain-containing protein | BIFBRE_04903 |
| + | 1.6 | -1.1 | D4BRG3 | Universal stress family protein | BIFBRE_04699 |
| + | 6.1 | 2.8 | D4BS03 | ROK family protein | BIFBRE_04894 |
| + | 6.6 | 3.2 | D4BMY8 | Beta-galactosidase (Beta-gal) (EC 3.2.1.23) | BIFBRE_03435 |
| + | 3.0 | -0.4 | D4BPB4 | Elongation factor 4 (EF-4) (EC 3.6.5.n1) (Ribosomal back-translocase LepA) | lepA BIFBRE_03923 |
| + | 3.6 | -1.0 | D4BPK5 | Transporter, major intrinsic protein (MIP) family protein | BIFBRE_04019 |
| + | 2.1 | 0.7 | D4BLN7 | Protease HtpX homolog (EC 3.4.24.-) | htpX BIFBRE_02973 |
| + | 1.4 | 0.9 | D4BRP1 | Oxidoreductase, aldo/keto reductase family protein | BIFBRE_04781 |
| + | 2.3 | 0.9 | D4BLB8 | LytTr DNA-binding domain protein | BIFBRE_02849 |
| + | 2.1 | -0.7 | D4BQ32 | ABC transporter, permease protein | BIFBRE_04186 |
| + | 2.4 | 1.8 | D4BQ36 | Efflux ABC transporter, permease protein | BIFBRE_04190 |
| + | 1.7 | -0.6 | D4BQC3 | UDP-N-acetylglucosamine 1-carboxyvinyltransferase (EC 2.5.1.7) (Enoylpyruvate transferase) (UDP-N-acetylglucosamine enolpyruvyl transferase) (EPT) | murA BIFBRE_04292 |
| + | 2.9 | 1.1 | D4BR73 | Aminotransferase (EC 2.6.1.-) | BIFBRE_04608 |
| + | 3.9 | -1.6 | D4BS46 | Efflux ABC transporter, permease protein | BIFBRE_04937 |
| + | 2.6 | 1.7 | D4BPF2 | Amidohydrolase family protein | BIFBRE_03961 |
| + | 3.0 | -0.4 | D4BLD9 | Putative integral membrane protein MviN | BIFBRE_02872 |
| + | 3.1 | -3.9 | D4BMK5 | ABC transporter, permease protein | BIFBRE_03302 |
| + | 1.7 | -1.3 | D4BN45 | ABC transporter, substrate-binding protein | BIFBRE_03493 |
| + | 2.2 | -0.5 | D4BQK4 | Phosphate transport system permease protein PstA | pstA BIFBRE_04377 |
| + | 1.7 | -1.0 | D4BR51 | Branched-chain amino acid ABC transporter, permease protein | BIFBRE_04586 |
| + | 3.4 | 0.9 | D4BPC4 | ABC transporter, substrate-binding protein, family 5 | BIFBRE_03933 |
| + | 3.0 | 0.5 | D4BML1 | Transcriptional regulator, LacI family | BIFBRE_03308 |
| + | 3.4 | -1.2 | D4BQU7 | Ribosome-binding factor A | rbfA BIFBRE_04475 |
| + | 3.5 | -2.5 | D4BMK4 | ABC transporter, ATP-binding protein | BIFBRE_03301 |
| + | 5.0 | -0.8 | D4BN22 | ABC transporter, substrate-binding protein, family 5 | BIFBRE_03470 |
| + | 1.7 | 0.8 | D4BNH7 | Acetylornithine aminotransferase (ACOAT) (EC 2.6.1.11) | argD BIFBRE_03627 |
| + | 4.1 | 1.5 | D4BN20 | Alpha amylase, catalytic domain protein | BIFBRE_03468 |
| + | 1.7 | 0.6 | D4BPF0 | Aspartate carbamoyltransferase (EC 2.1.3.2) (Aspartate transcarbamylase) (ATCase) | pyrB BIFBRE_03959 |
| + | 2.4 | 0.9 | D4BMG6 | DUF2207 domain-containing protein | BIFBRE_03261 |
| + | 1.6 | 0.9 | D4BQB6 | Dihydroorotate oxidase (EC 1.3.98.1) | BIFBRE_04285 |
| + | 2.2 | 1.1 | D4BPL9 | FHA domain protein | BIFBRE_04034 |
| + | 4.2 | -0.6 | D4BRQ2 | ATP-dependent helicase HrpA | hrpA BIFBRE_04792 |
| + | 7.7 | -3.6 | D4BQ14 | Xylulose kinase (Xylulokinase) (EC 2.7.1.17) | xylB BIFBRE_04242 |
| + | 2.3 | 0.8 | D4BLM5 | Pyridoxamine 5'-phosphate oxidase family protein | BIFBRE_02960 |
| + | 2.8 | -2.1 | D4BR53 | ABC transporter, ATP-binding protein | BIFBRE_04588 |
| + | 2.3 | -1.0 | D4BMV9 | PD-(D/E)XK endonuclease-like domain-containing protein | BIFBRE_03406 |
| + | 4.9 | 1.4 | D4BLM9 | 8-oxo-dGTP diphosphatase (EC 3.6.1.55) | BIFBRE_02965 |
| + | 2.8 | 1.2 | D4BNH5 | Arginine biosynthesis bifunctional protein ArgJ [Cleaved into: Arginine biosynthesis bifunctional protein ArgJ alpha chain; Arginine biosynthesis bifunctional protein ArgJ beta chain] [Includes: Glutamate N-acetyltransferase (EC 2.3.1.35) (Ornithine acetyltransferase) (OATase) (Ornithine transacetylase); Amino-acid acetyltransferase (EC 2.3.1.1) (N-acetylglutamate synthase) (AGSase)] | argJ BIFBRE_03625 |
| + | 2.5 | 0.9 | D4BR66 | Methionine aminopeptidase (MAP) (MetAP) (EC 3.4.11.18) (Peptidase M) | map BIFBRE_04601 |
| + | 2.8 | 1.0 | D4BS18 | Aldose 1-epimerase | BIFBRE_04909 |
| + | 2.0 | 1.3 | D4BMF5 | Nudix hydrolase domain-containing protein | BIFBRE_03250 |
| + | 2.0 | 0.6 | D4BS01 | ROK family protein | BIFBRE_04892 |
| + | 2.7 | -1.0 | D4BQ33 | Beta-carotene 15,15'-monooxygenase | BIFBRE_04187 |
| + | 1.7 | -1.8 | D4BNZ3 | ABC transporter, ATP-binding protein | BIFBRE_03801 |
| + | 3.7 | 0.5 | D4BP82 | Multifunctional fusion protein [Includes: Shikimate kinase (SK) (EC 2.7.1.71); 3-dehydroquinate synthase (DHQS) (EC 4.2.3.4)] | aroB aroK BIFBRE_03890 |
| + | 2.6 | 0.5 | D4BPV9 | IMP dehydrogenase family protein | BIFBRE_04124 |
| + | 7.1 | 4.1 | D4BL97 | Galactose-1-phosphate uridylyltransferase (Gal-1-P uridylyltransferase) (EC 2.7.7.12) (UDP-glucose--hexose-1-phosphate uridylyltransferase) | galT BIFBRE_02829 |
| + | 1.7 | -0.9 | D4BR52 | ABC transporter, ATP-binding protein | BIFBRE_04587 |
| + | 4.6 | -1.5 | D4BM70 | RmuC domain protein | BIFBRE_03162 |
| + | 2.7 | -0.7 | D4BMX3 | Cell division protein FtsX | BIFBRE_03420 |
| + | 3.0 | -0.6 | D4BRR2 | UvrD-like helicase ATP-binding domain-containing protein | BIFBRE_04803 |
| + | 3.7 | -1.0 | D4BLE0 | Thioredoxin-disulfide reductase (EC 1.8.1.9) | trxB BIFBRE_02873 |
| + | 1.7 | 1.2 | D4BQP8 | Glyoxalase family protein | BIFBRE_04425 |
| + | 2.1 | -0.5 | D4BMB8 | Peptidyl-prolyl cis-trans isomerase (EC 5.2.1.8) | BIFBRE_03211 |
| + | 3.6 | -0.6 | D4BM26 | Signal recognition particle receptor FtsY (SRP receptor) (EC 3.6.5.4) | ftsY BIFBRE_03117 |
| + | 2.0 | 1.1 | D4BQI5 | Kinase, PfkB family | BIFBRE_04358 |
| + | 1.4 | 0.9 | D4BN88 | Glycerophosphodiester phosphodiesterase family protein | BIFBRE_03537 |
| + | 2.6 | 1.0 | D4BLN9 | Fructose-bisphosphate aldolase (FBP aldolase) (EC 4.1.2.13) | fbaA BIFBRE_02976 |
| + | 1.6 | 0.9 | D4BM31 | MATE efflux family protein | BIFBRE_03122 |
| + | 6.6 | 4.1 | D4BMY6;D4BMM6;D4BME5 | ABC transporter, permease protein | BIFBRE_03433, BIFBRE_03323, BIFBRE_03239 |
| + | 3.1 | 1.9 | D4BP97 | Histidine triad domain protein | BIFBRE_03906 |
| + | 8.4 | 3.8 | D4BMY5 | ABC transporter, permease protein | BIFBRE_03432 |
| + | 3.8 | 1.3 | D4BLP7 | Sucrose phosphorylase (EC 2.4.1.7) | gtfA BIFBRE_02984 |
| + | 4.6 | -0.7 | D4BP66 | Small ribosomal subunit protein uS4 | rpsD BIFBRE_03874 |
| + | 3.6 | 0.7 | D4BNE0 | Aminoglycoside phosphotransferase domain-containing protein | BIFBRE_03589 |
| + | 3.3 | -0.4 | D4BMV0 | DNA-directed RNA polymerase subunit beta (RNAP subunit beta) (EC 2.7.7.6) (RNA polymerase subunit beta) (Transcriptase subunit beta) | rpoB BIFBRE_03397 |
| + | 6.5 | 3.4 | D4BQW4 | ABC transporter, solute-binding protein | BIFBRE_04493 |
| + | 2.0 | 0.5 | D4BMZ7 | ABC transporter, ATP-binding protein | BIFBRE_03444 |
| + | 14.4 | -4.8 | D4BQW0 | Phosphotransferase system, EIIC | BIFBRE_04488 |
| + | 2.3 | 2.1 | D4BMY3 | Sugar-binding domain protein | BIFBRE_03431 |
| + | 2.7 | 1.7 | D4BLH4 | carbonic anhydrase (EC 4.2.1.1) | cah BIFBRE_02909 |
| + | 6.2 | 2.0 | D4BLB1 | ABC transporter, ATP-binding protein | BIFBRE_02843 |
| + | 2.2 | -1.1 | D4BN47 | Aldehyde dehydrogenase | BIFBRE_03495 |
| + | 6.2 | 3.0 | D4BL96 | Putative glucose-6-phosphate 1-epimerase (EC 5.1.3.15) | BIFBRE_02828 |
| + | 2.3 | -0.6 | D4BMG2 | Polyribonucleotide nucleotidyltransferase (EC 2.7.7.8) (Polynucleotide phosphorylase) (PNPase) | pnp gpsI BIFBRE_03257 |
| + | 2.1 | -0.7 | D4BMU4 | UPF0210 protein BIFBRE_03391 | BIFBRE_03391 |
| + | 3.6 | 0.6 | D4BMK3 | Glucose-6-phosphate isomerase (GPI) (EC 5.3.1.9) (Phosphoglucose isomerase) (PGI) (Phosphohexose isomerase) (PHI) | pgi BIFBRE_03300 |
| + | 10.3 | 2.6 | D4BQW7 | 1,3-beta-galactosyl-N-acetylhexosamine phosphorylase (EC 2.4.1.211) | gnpA BIFBRE_04496 |
| + | 3.9 | -1.0 | D4BQF2 | GTPase Obg (EC 3.6.5.-) (GTP-binding protein Obg) | cgtA obg BIFBRE_04321 |
| + | 2.7 | 1.1 | D4BPI6 | Transketolase (EC 2.2.1.1) | tkt BIFBRE_03995 |
| + | 2.8 | 1.6 | D4BS00 | Kinase, PfkB family | BIFBRE_04891 |
| + | 2.1 | 0.7 | D4BR22 | tRNA(Ile)-lysidine synthase (EC 6.3.4.19) (tRNA(Ile)-2-lysyl-cytidine synthase) (tRNA(Ile)-lysidine synthetase) | tilS BIFBRE_04555 |
| + | 2.1 | -0.5 | D4BMB9 | Tat pathway signal sequence domain protein | BIFBRE_03212 |
| + | 4.6 | -1.0 | D4BM83 | Uncharacterized protein | BIFBRE_03175 |
| + | 3.5 | -0.6 | D4BS49 | ABC transporter, substrate-binding protein, family 5 | BIFBRE_04940 |
| + | 2.1 | -0.5 | D4BQG8 | biotin carboxylase (EC 6.3.4.14) | BIFBRE_04338 |
| + | 2.4 | 0.8 | D4BS21 | Glyceraldehyde-3-phosphate dehydrogenase, type I (EC 1.2.1.-) | gap BIFBRE_04912 |
| + | 2.6 | -0.8 | D4BS93 | DNA topoisomerase (ATP-hydrolyzing) (EC 5.6.2.2) | BIFBRE_04985 |
| + | 2.4 | -0.6 | D4BQR7 | Large ribosomal subunit protein uL16 | rplP BIFBRE_04444 |
| + | 2.5 | -0.4 | D4BP77 | Alanine--tRNA ligase (EC 6.1.1.7) (Alanyl-tRNA synthetase) (AlaRS) | alaS BIFBRE_03885 |
| + | 3.1 | -0.6 | D4BMS0 | non-specific protein-tyrosine kinase (EC 2.7.10.2) | BIFBRE_03367 |
| + | 3.0 | -1.1 | D4BNM0 | UPF0182 protein BIFBRE_03670 | BIFBRE_03670 |
| + | 2.4 | -0.5 | D4BPV6 | Peptide deformylase (PDF) (EC 3.5.1.88) (Polypeptide deformylase) | def BIFBRE_04121 |
| + | 1.8 | 0.5 | D4BRE0 | DNA-binding protein HB1 | hup BIFBRE_04676 |
| + | 5.6 | -2.3 | D4BLQ5 | Transcriptional regulator, LacI family | BIFBRE_02992 |
| + | 3.9 | -0.6 | D4BQR6 | Small ribosomal subunit protein uS3 | rpsC BIFBRE_04443 |
| + | 4.2 | -0.7 | D4BMK8 | Large ribosomal subunit protein bL19 | rplS BIFBRE_03304 |
| + | 1.9 | 1.4 | D4BQV3 | Ribose-5-phosphate isomerase A (EC 5.3.1.6) (Phosphoriboisomerase A) (PRI) | rpiA BIFBRE_04481 |
| + | 5.2 | -0.6 | D4BQR3 | Large ribosomal subunit protein uL2 | rplB BIFBRE_04440 |
| + | 7.5 | -1.1 | D4BR96 | ABC transporter, ATP-binding protein | BIFBRE_04632 |
| + | 1.4 | 1.0 | D4BP45 | ABC transporter, ATP-binding protein | BIFBRE_03853 |
| + | 3.3 | -3.7 | D4BLQ4 | Alpha amylase, catalytic domain protein | BIFBRE_02991 |
| + | 1.9 | -1.2 | D4BS47 | ABC transporter, ATP-binding protein | BIFBRE_04938 |
| + | 2.3 | -0.6 | D4BN73 | Small ribosomal subunit protein uS7 | rpsG BIFBRE_03522 |
| + | 1.5 | 0.8 | D4BNQ9 | Hydrolase, NUDIX family | BIFBRE_03710 |
| + | 2.6 | -1.1 | D4BM24 | Glycosyltransferase, group 2 family protein (EC 2.4.-.-) | BIFBRE_03115 |
| + | 3.1 | -1.7 | D4BQK2 | Phosphate-binding protein | pstS BIFBRE_04375 |
| + | 2.3 | 1.1 | D4BQY2 | 1,4-dihydroxy-2-naphthoate octaprenyltransferase (DHNA-octaprenyltransferase) (EC 2.5.1.74) | menA BIFBRE_04512 |
| + | 1.3 | 1.4 | D4BS66 | tRNA dimethylallyltransferase (EC 2.5.1.75) (Dimethylallyl diphosphate:tRNA dimethylallyltransferase) (DMAPP:tRNA dimethylallyltransferase) (DMATase) (Isopentenyl-diphosphate:tRNA isopentenyltransferase) (IPP transferase) (IPPT) (IPTase) | miaA BIFBRE_04957 |
| + | 1.2 | -1.4 | D4BRP0 | Uncharacterized protein | BIFBRE_04780 |
| + | 2.4 | -1.2 | D4BMM0 | ABC transporter, ATP-binding protein | BIFBRE_03317 |
| + | 1.8 | -1.1 | D4BR49 | Receptor family ligand-binding protein | BIFBRE_04584 |
| + | 1.9 | -0.6 | D4BMP0 | Glycosyltransferase, group 1 family protein (EC 2.4.-.-) | BIFBRE_03337 |
| + | 1.6 | 1.2 | D4BRS7 | Lipoprotein | BIFBRE_04818 |
| + | 6.2 | 1.4 | D4BNW1 | Formate acetyltransferase (EC 2.3.1.54) (Pyruvate formate-lyase) | pflB BIFBRE_03767 |
| + | 1.9 | -0.5 | D4BLE9 | Large ribosomal subunit protein bL34 | rpmH BIFBRE_02882 |
| + | 4.2 | -3.5 | D4BP39 | DEAD/DEAH box helicase | BIFBRE_03847 |
| + | 1.6 | 1.0 | D4BLK8 | Glutamine amidotransferase, class I | BIFBRE_02943 |
| + | 2.7 | -0.5 | D4BQR9 | Small ribosomal subunit protein uS17 | rpsQ BIFBRE_04446 |
| + | 1.8 | 0.7 | D4BP11 | HipA-like C-terminal domain-containing protein | BIFBRE_03819 |
| + | 7.2 | 1.2 | D4BNQ8 | Pyruvate kinase (EC 2.7.1.40) | pyk BIFBRE_03709 |
| + | 3.1 | -0.6 | D4BQF0 | Large ribosomal subunit protein bL21 | rplU BIFBRE_04319 |
| + | 2.9 | 1.0 | D4BLG5 | Glutamate dehydrogenase | BIFBRE_02898 |
| + | 2.4 | -0.6 | D4BQG0 | Large ribosomal subunit protein uL11 | rplK BIFBRE_04330 |
| + | 2.6 | -0.5 | D4BQR1 | Large ribosomal subunit protein uL4 | rplD BIFBRE_04438 |
| + | 2.3 | 2.0 | D4BPD2 | Uncharacterized protein | BIFBRE_03941 |
| + | 2.3 | -0.4 | D4BMN9 | Glycosyltransferase, group 1 family protein (EC 2.4.-.-) | BIFBRE_03336 |
| + | 4.4 | -0.6 | D4BQR2 | Large ribosomal subunit protein uL23 | rplW BIFBRE_04439 |
| + | 2.4 | -0.5 | D4BN74 | Elongation factor G (EF-G) | fusA BIFBRE_03523 |
| + | 2.0 | 0.9 | D4BRM7 | Oxidoreductase, aldo/keto reductase family protein | BIFBRE_04763 |
| + | 2.6 | -1.0 | D4BN03 | Amino acid transporter | BIFBRE_03450 |
| + | 6.8 | 2.1 | D4BQX0 | UDP-glucose 4-epimerase (EC 5.1.3.2) | galE BIFBRE_04499 |
| + | 1.7 | 0.6 | D4BQL1 | FHA domain protein | BIFBRE_04384 |
| + | 6.0 | -2.2 | D4BMZ9 | Cys/Met metabolism PLP-dependent enzyme | BIFBRE_03446 |
| + | 3.5 | 1.1 | D4BNU9 | D-xylulose 5-phosphate/D-fructose 6-phosphate phosphoketolase | xfp BIFBRE_03753 |
| + | 1.8 | 2.0 | D4BME8 | Transcriptional regulator, TetR family | BIFBRE_03242 |
| + | 3.5 | -0.9 | D4BP62 | DNA 3'-5' helicase (EC 5.6.2.4) | BIFBRE_03870 |
| + | 2.4 | -0.6 | D4BQS4 | Small ribosomal subunit protein uS8 | rpsH BIFBRE_04451 |
| + | 3.0 | -0.7 | D4BQF4 | Polysaccharide deacetylase | BIFBRE_04323 |
| + | 3.2 | -0.6 | D4BLD8 | Secreted protein | BIFBRE_02871 |
| + | 2.7 | -0.8 | D4BL90 | Putative potassium/sodium efflux P-type ATPase, fungal-type | BIFBRE_02822 |
| + | 4.0 | -0.7 | D4BQS0 | Large ribosomal subunit protein uL14 | rplN BIFBRE_04447 |
| + | 4.3 | 2.1 | D4BQ60 | ABC transporter, ATP-binding protein | BIFBRE_04215 |
| + | 2.8 | -0.7 | D4BN72 | Small ribosomal subunit protein uS12 | rpsL BIFBRE_03521 |
| + | 4.2 | -0.7 | D4BR24 | ATP-dependent zinc metalloprotease FtsH (EC 3.4.24.-) | hflB ftsH BIFBRE_04557 |
| + | 11.4 | 4.5 | D4BS04 | Glucosamine-6-phosphate deaminase (EC 3.5.99.6) (GlcN6P deaminase) (GNPDA) (Glucosamine-6-phosphate isomerase) | nagB BIFBRE_04895 |
| + | 3.4 | 0.3 | D4BS99 | RelA/SpoT family protein | BIFBRE_04991 |
| + | 6.2 | -2.3 | D4BLY0 | Transcriptional regulator, LacI family | BIFBRE_03067 |
| + | 3.6 | -1.3 | D4BLR2 | Alpha amylase, catalytic domain protein | BIFBRE_02999 |
| + | 2.7 | 1.0 | D4BLB4 | Ascorbate-specific PTS system EIIC component (Ascorbate-specific permease IIC component UlaA) | BIFBRE_02846 |
| + | 2.4 | -0.8 | D4BMX8 | ABC transporter, substrate-binding protein, family 3 | BIFBRE_03425 |
| + | 1.7 | 0.9 | D4BS14 | Sigma-70 region 2 | BIFBRE_04905 |
| + | 2.8 | -0.5 | D4BQS9 | Large ribosomal subunit protein uL15 | rplO BIFBRE_04456 |
| + | 2.0 | -0.5 | D4BP79 | Endolytic murein transglycosylase (EC 4.2.2.29) (Peptidoglycan lytic transglycosylase) (Peptidoglycan polymerization terminase) | mltG BIFBRE_03887 |
| + | 3.8 | 1.9 | D4BP48 | Divergent AAA domain protein | BIFBRE_03856 |
| + | 2.7 | 0.5 | D4BNS4 | Phosphate acetyltransferase (EC 2.3.1.8) (Phosphotransacetylase) | pta BIFBRE_03727 |
| + | 1.7 | 0.8 | D4BRD9 | Pup--protein ligase (EC 6.3.1.19) (Proteasome accessory factor A) (Pup-conjugating enzyme) | pafA BIFBRE_04675 |
| + | 4.9 | 1.4 | D4BMT9 | Galactose-1-phosphate uridylyltransferase (EC 2.7.7.12) | galT BIFBRE_03386 |
| + | 3.4 | -0.6 | D4BRG0 | Cold-shock DNA-binding domain protein | BIFBRE_04696 |
| + | 3.7 | -0.5 | D4BQT5 | Small ribosomal subunit protein uS13 | rpsM BIFBRE_04462 |
| + | 2.4 | 0.5 | D4BMZ6 | ABC transporter, ATP-binding protein | BIFBRE_03443 |
| + | 2.9 | -0.9 | D4BRH8 | ABC transporter, substrate-binding protein, family 3 | BIFBRE_04714 |
| + | 3.4 | -0.5 | D4BM45 | Small ribosomal subunit protein bS18 | rpsR BIFBRE_03136 |
| + | 2.9 | 1.3 | D4BM62 | Major facilitator superfamily (MFS) profile domain-containing protein | BIFBRE_03154 |
| + | 1.5 | 1.2 | D4BRF9 | histidine kinase (EC 2.7.13.3) | BIFBRE_04695 |
| + | 2.4 | 0.5 | D4BN93 | Transcriptional regulator, LacI family | BIFBRE_03542 |
| + | 2.5 | -0.6 | D4BMW4 | Ribonuclease J (RNase J) (EC 3.1.-.-) | rnj BIFBRE_03411 |
| + | 2.9 | -0.5 | D4BQP3 | Small ribosomal subunit protein uS9 | rpsI BIFBRE_04420 |
| + | 3.5 | 1.0 | D4BS67 | protein adenylyltransferase (EC 2.7.7.108) | BIFBRE_04958 |
| + | 3.3 | 1.6 | D4BMC1 | Thioredoxin | BIFBRE_03214 |
| + | 2.9 | -1.0 | D4BM68 | Copper-exporting ATPase (EC 3.6.3.4) | BIFBRE_03160 |
| + | 5.2 | -1.7 | D4BLR1 | 4-alpha-glucanotransferase (EC 2.4.1.25) (Amylomaltase) (Disproportionating enzyme) | malQ BIFBRE_02998 |
| + | 1.3 | 1.1 | D4BS98 | Deoxyuridine 5'-triphosphate nucleotidohydrolase (dUTPase) (EC 3.6.1.23) (dUTP pyrophosphatase) | dut BIFBRE_04990 |
| + | 2.4 | -0.5 | D4BQU9 | Bifunctional riboflavin kinase/FMN adenylyltransferase (EC 2.7.1.26) (EC 2.7.7.2) (Riboflavin biosynthesis protein RibF) | BIFBRE_04477 |
| + | 4.3 | -0.7 | D4BMH4 | Small ribosomal subunit protein bS16 | rpsP BIFBRE_03269 |
| + | 2.9 | 0.9 | D4BP08 | Cupin domain protein | BIFBRE_03816 |
| + | 1.9 | -2.7 | D4BQ16 | ABC transporter, ATP-binding protein | BIFBRE_04244 |
| + | 3.2 | -0.6 | D4BPG3 | ABC transporter, ATP-binding protein | BIFBRE_03972 |
| + | 2.8 | -0.5 | D4BQR5 | Large ribosomal subunit protein uL22 | rplV BIFBRE_04442 |
| + | 4.6 | -0.6 | D4BLI9 | Phosphoenolpyruvate carboxylase | ppc BIFBRE_02924 |
| + | 3.5 | -0.4 | D4BS68 | FtsK/SpoIIIE family protein | BIFBRE_04959 |
| + | 2.2 | 0.9 | D4BQ66 | ABC transporter, ATP-binding protein | BIFBRE_04221 |
| + | 3.8 | -2.1 | D4BQ15 | Sugar-binding domain protein | BIFBRE_04243 |
| + | 7.3 | 3.1 | D4BQW9 | Phosphotransferase enzyme family | BIFBRE_04498 |
| + | 2.2 | -1.2 | D4BMU7 | Adenine DNA glycosylase (EC 3.2.2.31) | BIFBRE_03394 |
| + | 1.9 | -0.8 | D4BRC7 | SNARE-like domain protein | BIFBRE_04663 |
| + | 2.9 | 1.0 | D4BPJ8 | Holliday junction branch migration complex subunit RuvB (EC 3.6.4.-) | ruvB BIFBRE_04012 |
| + | 3.0 | -0.6 | D4BMC0 | Lipoprotein | BIFBRE_03213 |
| + | 2.3 | -0.8 | D4BRI0 | ABC transporter, permease protein | BIFBRE_04716 |
| + | 2.9 | -0.7 | D4BRH9 | ABC transporter, permease protein | BIFBRE_04715 |
| + | 3.8 | -0.6 | D4BQT6 | Small ribosomal subunit protein uS11 | rpsK BIFBRE_04463 |
| + | 1.9 | 0.7 | D4BPF8 | DUF2974 domain-containing protein | BIFBRE_03967 |
| + | 3.2 | -0.6 | D4BNG3 | DUF4191 domain-containing protein | BIFBRE_03613 |
| + | 3.1 | 1.3 | D4BRP4 | Histidinol-phosphate aminotransferase (EC 2.6.1.9) (Imidazole acetol-phosphate transaminase) | hisC BIFBRE_04784 |
| + | 1.8 | 0.6 | D4BPD1 | thioredoxin-dependent peroxiredoxin (EC 1.11.1.24) (Bacterioferritin comigratory protein) (Thioredoxin peroxidase) | BIFBRE_03940 |
| + | 1.6 | -0.9 | D4BPU6 | Phosphoribosyl-AMP cyclohydrolase (PRA-CH) (EC 3.5.4.19) | hisI BIFBRE_04111 |
| + | 1.4 | 1.2 | D4BNM5 | Enolase (EC 4.2.1.11) (2-phospho-D-glycerate hydro-lyase) (2-phosphoglycerate dehydratase) | eno BIFBRE_03675 |
| + | 4.2 | -1.2 | D4BN26 | CARDB domain-containing protein | BIFBRE_03474 |
| + | 3.0 | -0.6 | D4BQS1 | Large ribosomal subunit protein uL24 | rplX BIFBRE_04448 |
| + | 2.3 | -0.4 | D4BQS7 | Small ribosomal subunit protein uS5 | rpsE BIFBRE_04454 |
| + | 2.7 | -0.5 | D4BL95 | RNA methyltransferase, TrmH family, group 3 | BIFBRE_02827 |
| + | 1.9 | -0.7 | D4BPU3 | Uncharacterized protein | BIFBRE_04108 |
| + | 2.5 | -1.1 | D4BNZ4 | ABC transporter, permease protein | BIFBRE_03802 |
| + | 1.8 | 1.5 | D4BN30 | N5-carboxyaminoimidazole ribonucleotide mutase (N5-CAIR mutase) (EC 5.4.99.18) (5-(carboxyamino)imidazole ribonucleotide mutase) | purE BIFBRE_03478 |
| + | 5.5 | -0.9 | D4BM85 | Transcription termination factor Rho (EC 3.6.4.-) (ATP-dependent helicase Rho) | rho BIFBRE_03176 |
| + | 2.1 | 0.6 | D4BS89 | DEAD/DEAH box helicase | BIFBRE_04980 |
| + | 2.5 | -0.4 | D4BNQ4 | Small ribosomal subunit protein bS1 (30S ribosomal protein S1) | rpsA BIFBRE_03705 |
| + | 3.6 | -1.7 | D4BRR7 | Penicillin-binding protein, transpeptidase domain protein | BIFBRE_04807 |
| + | 4.5 | -1.1 | D4BS41 | Large ribosomal subunit assembly factor BipA (EC 3.6.5.-) (GTP-binding protein BipA) | typA bipA BIFBRE_04932 |
| + | 4.0 | -0.8 | D4BMN4 | Chemotaxis protein | BIFBRE_03331 |
| + | 4.9 | -0.8 | D4BR31 | Energy-dependent translational throttle protein EttA (EC 3.6.1.-) (Translational regulatory factor EttA) | ettA BIFBRE_04564 |
| + | 2.5 | -2.0 | D4BQ18 | Branched-chain amino acid ABC transporter, permease protein | BIFBRE_04246 |
| + | 1.2 | -2.5 | D4BM91 | ABC transporter, solute-binding protein | BIFBRE_03183 |
| + | 3.6 | 1.4 | D4BS70 | Competence/damage-inducible domain protein CinA | BIFBRE_04961 |
| + | 4.9 | -1.9 | D4BQ42 | Fatty acid ABC transporter ATP-binding/permease protein | BIFBRE_04196 |
| + | 2.5 | -1.0 | D4BLW2 | Dihydrodipicolinate synthetase family | BIFBRE_03049 |
| + | 5.2 | -0.7 | D4BQU6 | Translation initiation factor IF-2 | infB BIFBRE_04473 |
| + | 2.2 | 1.2 | D4BP53 | nicotinamidase (EC 3.5.1.19) (Nicotinamide deamidase) | BIFBRE_03861 |
| + | 2.3 | 1.3 | D4BPA9 | branched-chain-amino-acid transaminase (EC 2.6.1.42) | ilvE BIFBRE_03918 |
| + | 2.6 | -0.5 | D4BQV1 | Secreted protein | BIFBRE_04479 |
| + | 4.4 | -0.7 | D4BS26 | Large ribosomal subunit protein bL20 | rplT BIFBRE_04917 |
| + | 4.6 | 1.0 | D4BQP4 | Glycogen debranching enzyme GlgX (EC 3.2.1.-) | glgX BIFBRE_04421 |
| + | 5.0 | -2.0 | D4BP18 | Drug resistance MFS transporter, drug:H+ antiporter-2 family | BIFBRE_03826 |
| + | 3.8 | -0.6 | D4BQS5 | Large ribosomal subunit protein uL6 | rplF BIFBRE_04452 |
| + | 2.3 | 1.0 | D4BLA8 | Uncharacterized protein | BIFBRE_02840 |

- Purple indicate the proteins significantly over expressed in LNnT.

Supplementary Table 10. Significantly changed protein in LNT versus LNnT

| significance | log (P-value) | log 2-fold change | protein IDs | Protein names | Gene Names |
| --- | --- | --- | --- | --- | --- |
| + | 3.05952 | -1.55454 | D4BMY8 | Beta-galactosidase (Beta-gal) (EC 3.2.1.23) | BIFBRE_03435 |
| + | 3.905127 | 1.120227 | D4BNW5 | 3'-5' exonuclease (EC 3.6.1.-) | BIFBRE_03771 |
| + | 4.023011 | 1.051355 | D4BQW0 | Phosphotransferase system, EIIC | BIFBRE_04488 |
| + | 2.778071 | 1.928979 | D4BQN6 | Glycosyl hydrolase, family 1 | BIFBRE_04413 |
| + | 3.656212 | 1.278145 | D4BMM0 | ABC transporter, ATP-binding protein | BIFBRE_03317 |
| + | 2.605432 | -3.78248 | D4BNP7 | 1,4-alpha-glucan branching enzyme GlgB (EC 2.4.1.18) (1,4-alpha-D-glucan:1,4-alpha-D-glucan 6-glucosyl-transferase) (Alpha-(1->4)-glucan branching enzyme) (Glycogen branching enzyme) (BE) | glgB BIFBRE_03698 |
| + | 3.790146 | 1.927346 | D4BMZ9 | Cys/Met metabolism PLP-dependent enzyme | BIFBRE_03446 |

- Pink indicate the proteins significantly over expressed in LNT, while purple indicate the proteins significantly over expressed in LNnT.

Supplementary Figures

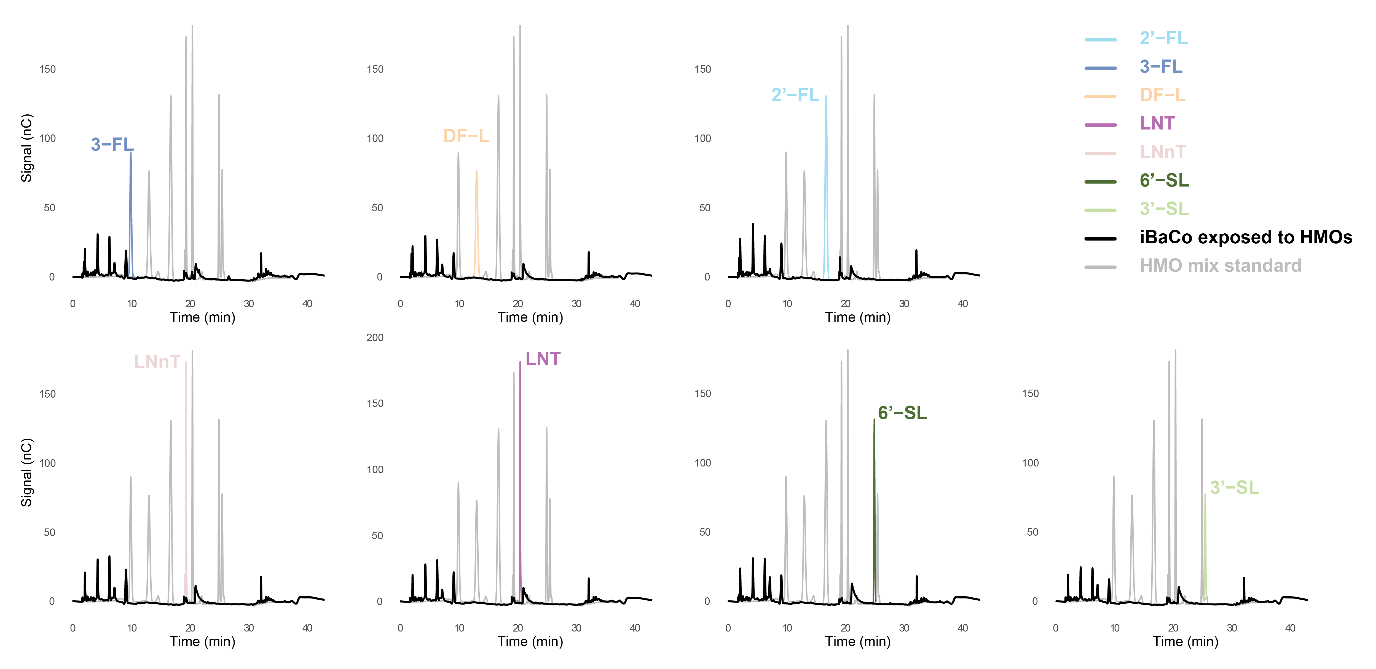

**Supplementary Figure 1.** **HMO consumption by iBaCo.** Supernatants were collected after 24 h culture and analyzed by HPAEC. Gray chromatographs indicate HMO standards, black chromatographs indicate iBaCo cultured in HMOs. HMO disappearances showed complete depletion of HMOs without monosaccharide release.

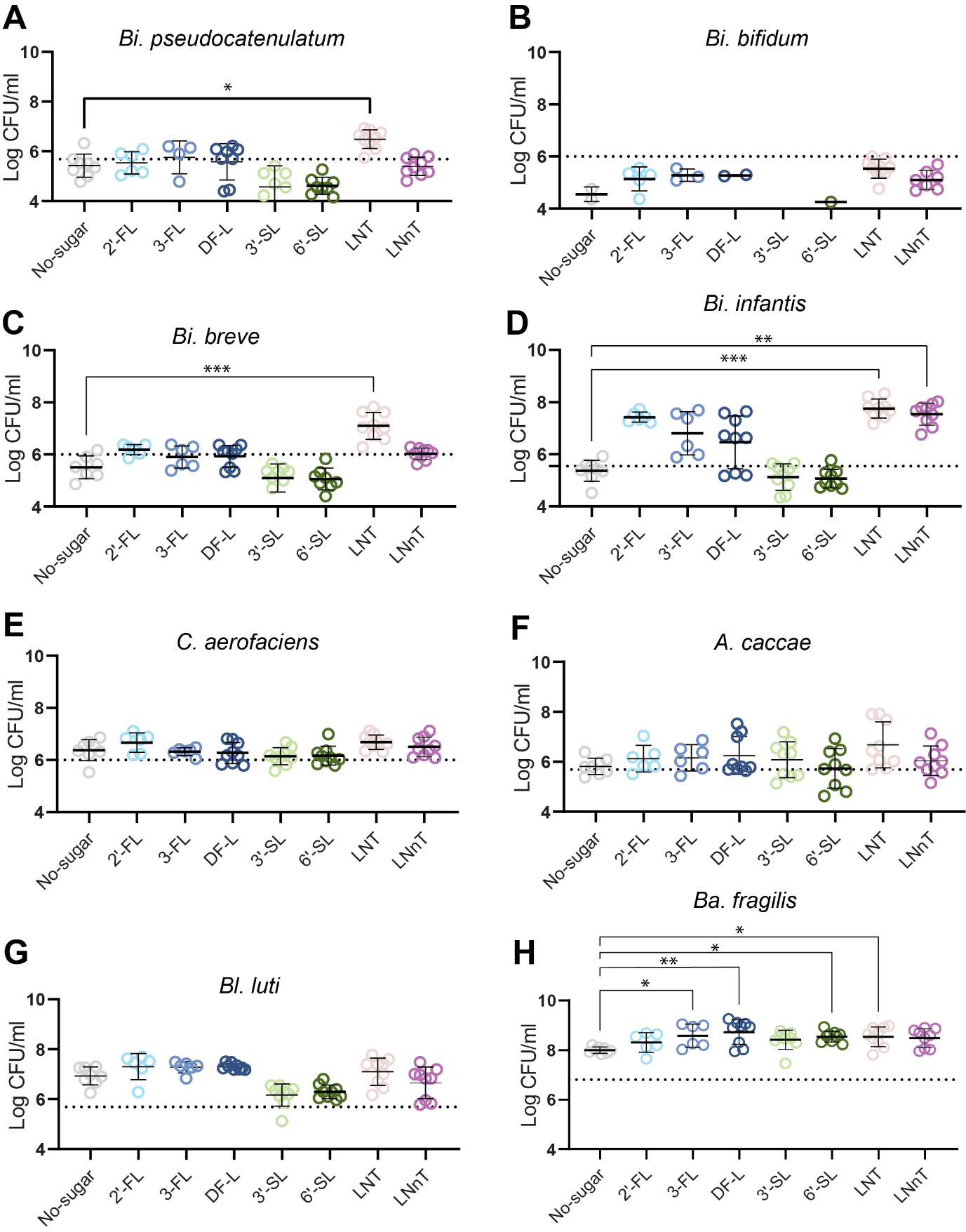

**Supplementary Figure 2.** **Absolute growth of iBaCo species.** Growth of (A) Bi. pseudocatenulatum, (B) Bi. bifidum, (C) Bi. breve, (D) bi. infantis, (E) C. aerofaciens, (F) A. caccae, (G) Bl. luti, and (H) Ba. fragilis. Data are pooled from three independent experiments; each point represents a biological replicate, bars indicate mean ± SD. Dashed lines denote inoculum levels at 0 h. Statistical significance was assessed by Kruskal–Wallis with Dunn’s test; *p < 0.05, *****p*** < 0.01, ******p*** < 0.001, *******p*** < 0.0001.

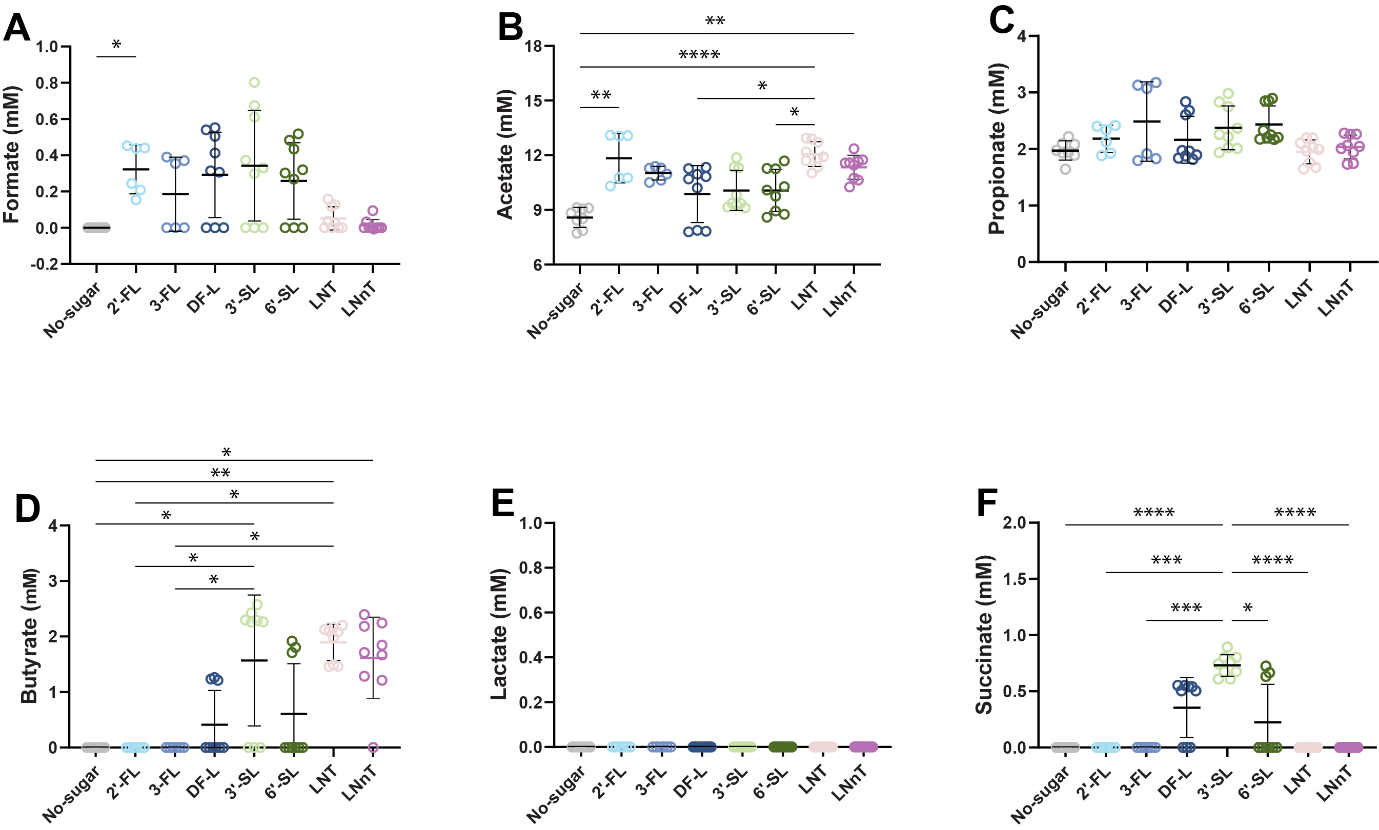
**Supplementary Figure 3.** **Organic acid profiles of iBaCo.** Concentrations of (A) formate, (B) acetate, (C) propionate, (D) butyrate, (E) lactate, and (F) succinate after 24 h culture in HMO-supplemented media, measured by HPLC. Data are from three independent experiments; points represent biological replicates, bars indicate mean ± SD. Dashed lines mark baseline concentrations in media. Kruskal–Wallis with Dunn’s test; *p < 0.05, *****p*** < 0.01, ******p*** < 0.001, *******p*** < 0.0001. Butyrate values <1.25 mM and succinate <0.5 mM were set to 0.

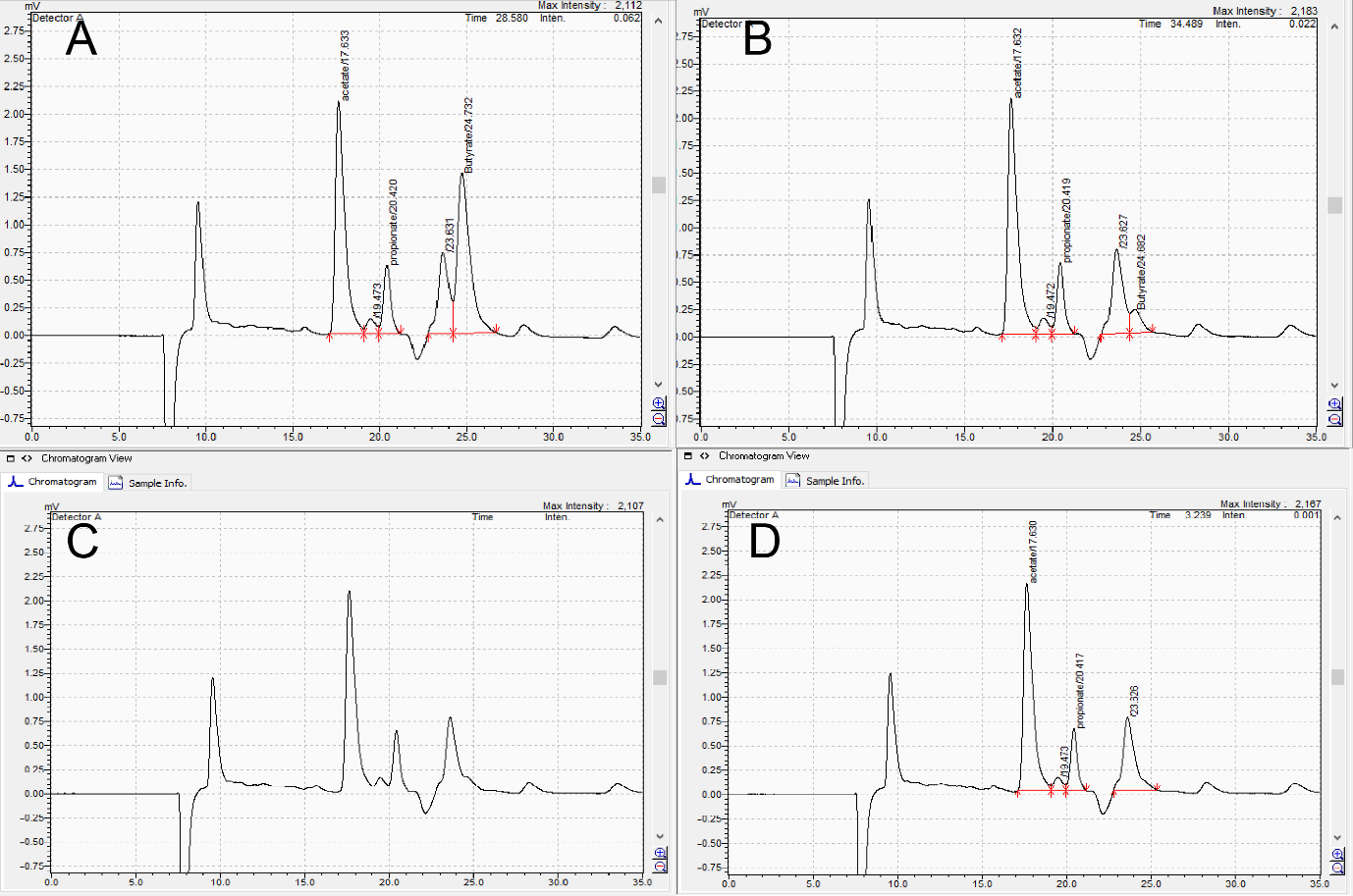

**Supplementary Figure 4.** **Detection limit of butyrate.** mYCFA medium was spiked with butyrate at (A) 10 mM, (B) 1.25 mM, (C) 0.625 mM, and (D) 0.3125 mM and analyzed by HPLC. No peak was observed below 0.625 mM.

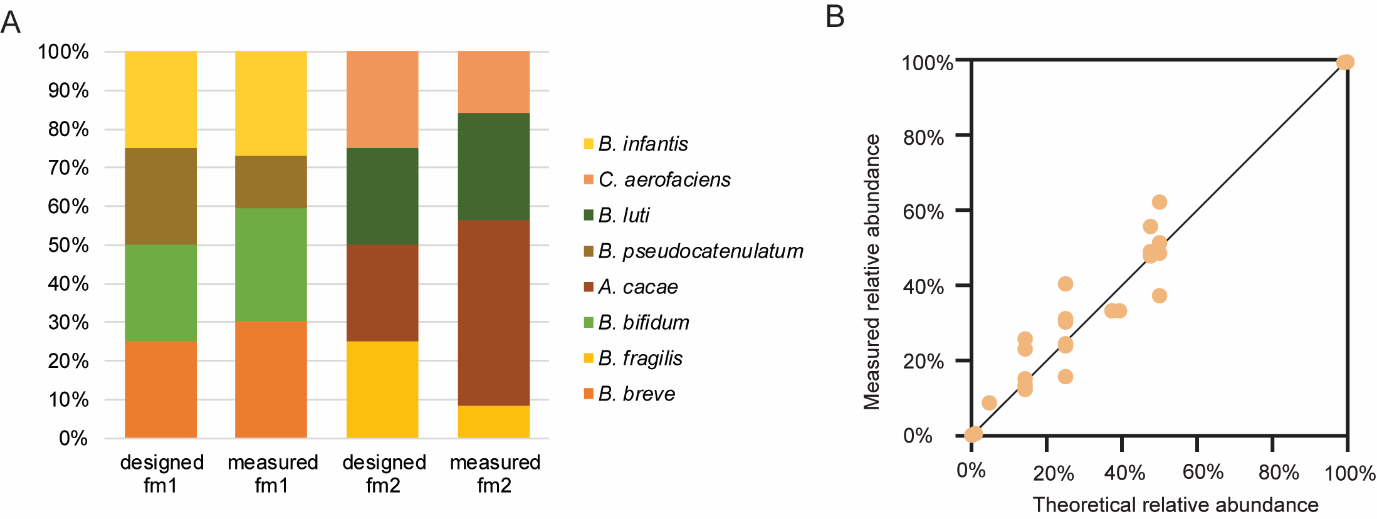

**Supplementary Figure 5: Mock community validation for Nanopore sequencing.** (A) Designed versus measured relative abundances of two mock communities. (B) Comparison of theoretical versus measured values.

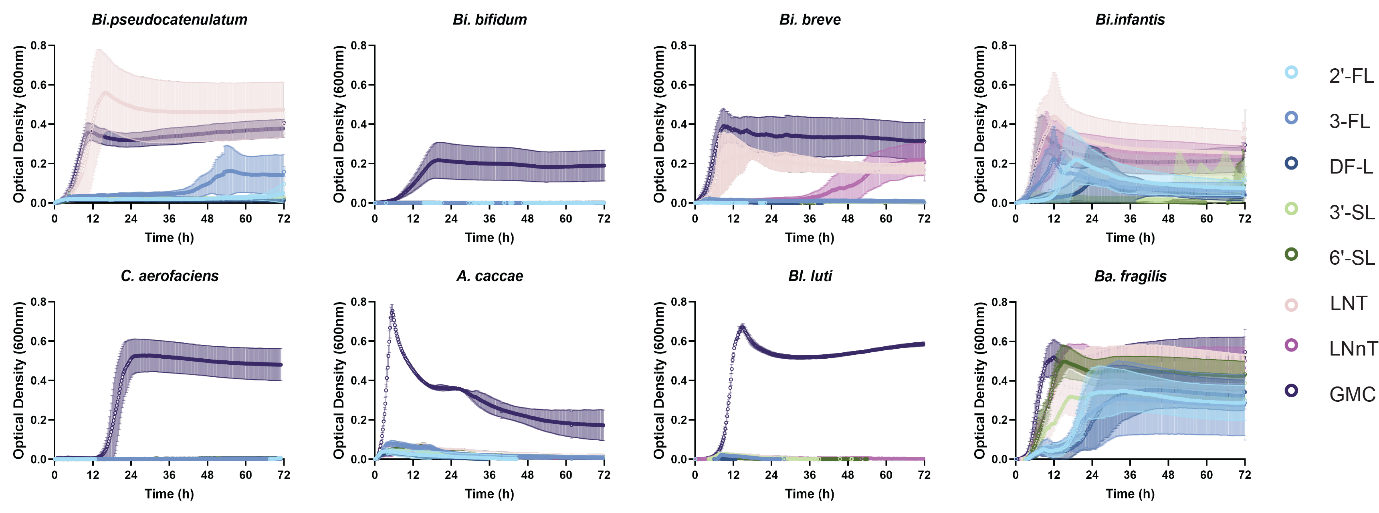

**Supplementary Figure 6.** **Growth curves of iBaCo species in HMO-supplemented YCFA**. Data are shown as mean ± SD. GMC: glucose, maltose, and cellobiose.

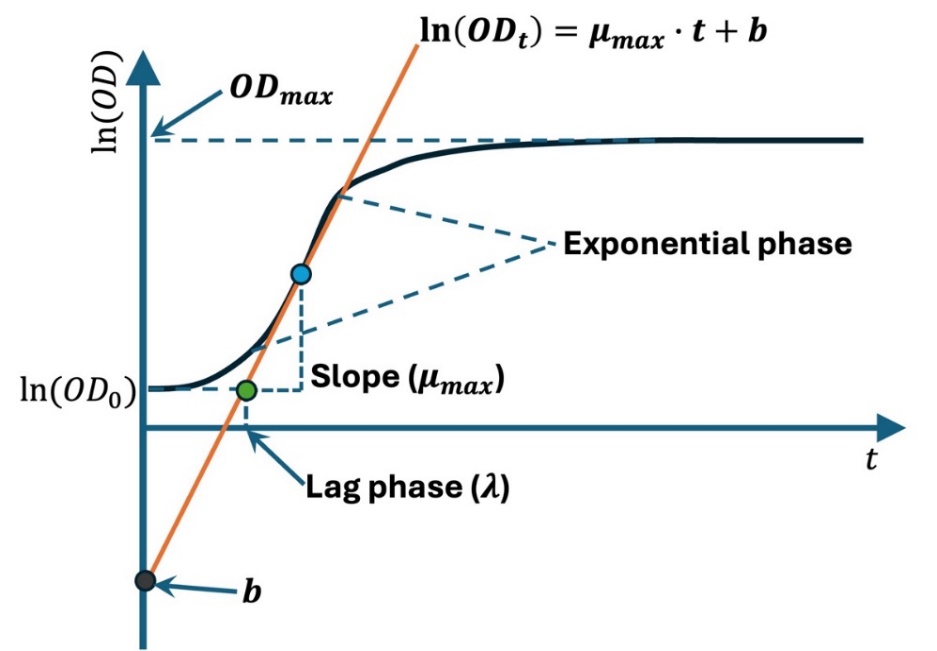

**Supplementary Figure 7 Schematic illustration of the growth curve analysis for estimating kinetic parameters.** The maximum specific growth rate ($\mu_{max}$) was calculated as the slope of the regression line (solid orange) fitted to the exponential phase of the $\ln(OD)$-time curve. The lag phase ($\lambda$) was determined by extrapolating this line to the initial $OD$ baseline. $b$ is the y-intercept of the regression line. The maximum optical density (${OD}_{max}$) is the highest $OD$ value reached during the cultivation period.

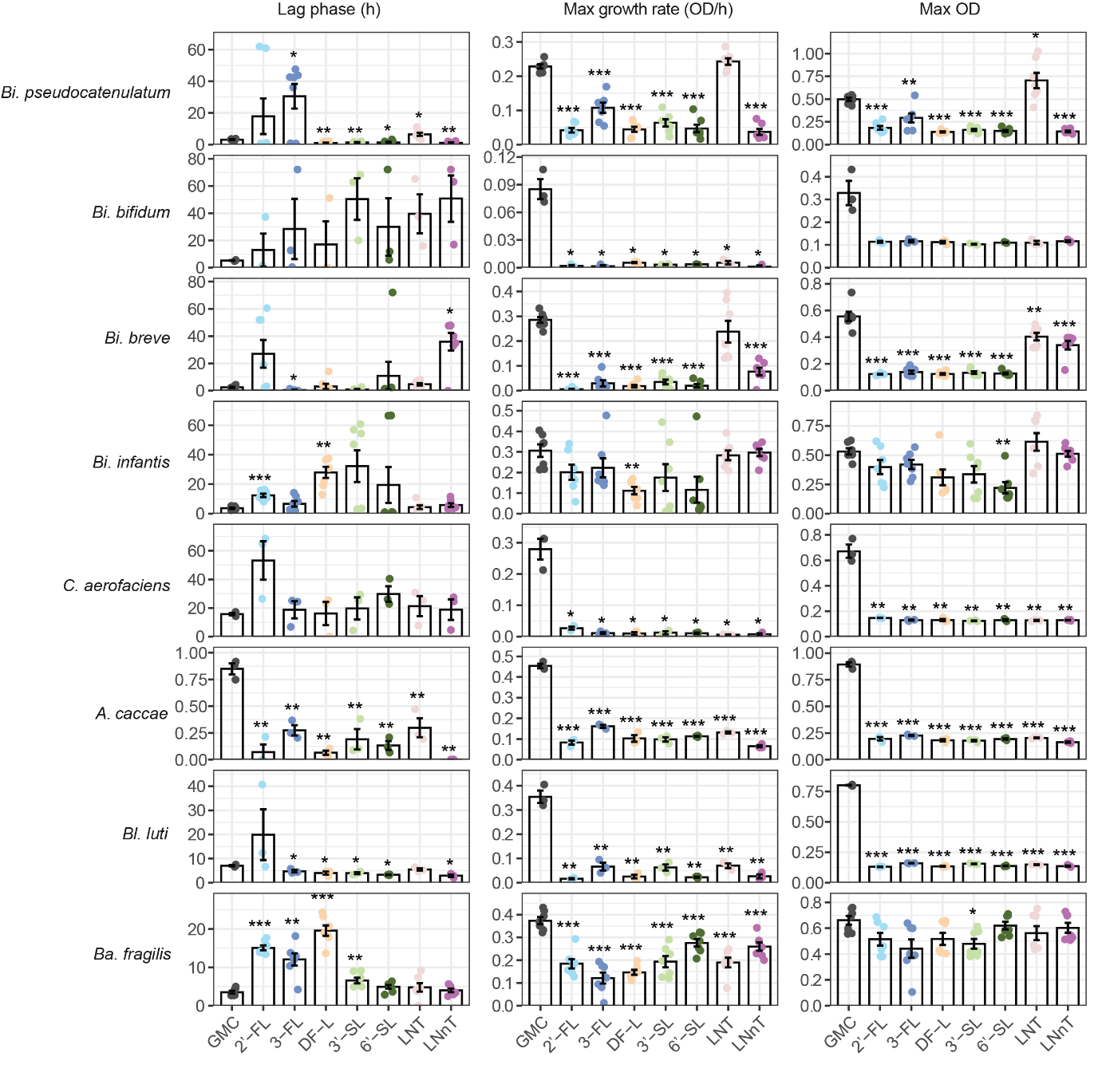
**Supplementary Figure 8.** **Growth parameters of individual iBaCo species in monoculture.** iBaCo species: *Bi. pseudocatenulatum*, *Bi. bifidum*, *Bi. breve*, *Bi. infantis*, *C. aerofaciens*, *A. caccae*, *Bl. luti*, *Ba. fragilis* were cultured in individually in YCFA media supplemented with single HMOs as the sole carbon source. Statistical significance was assessed by unpaired t-tests, comparing each HMO condition against the corresponding GMC control. *p < 0.05, *****p*** < 0.01, ******p*** < 0.001, *******p*** < 0.0001.

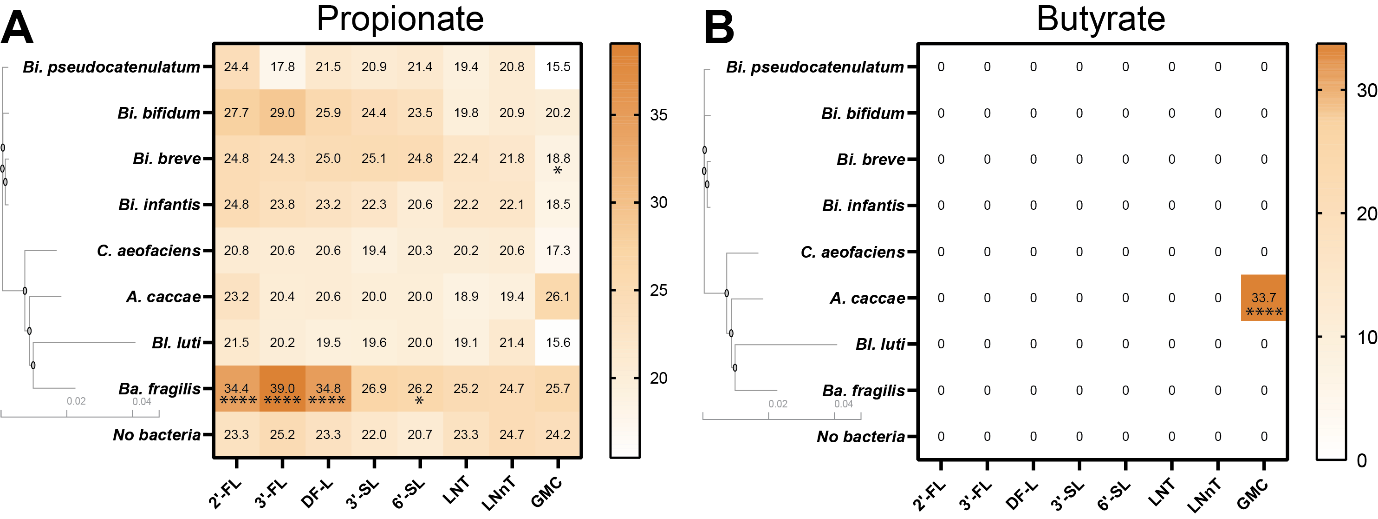

**Supplementary Figure 9.** **Production of propionate and butyrate by monocultures**. Concentrations of (A) propionate and (B) butyrate measured after 72 h. Data are mean of three independent experiments; points represent replicates, unit mM. Two-way ANOVA with Dunnett’s test, comparing each condition to “no bacteria” controls; *p < 0.05, *****p*** < 0.01, ******p*** < 0.001, *******p*** < 0.0001. Butyrate values below detection were set to 0.

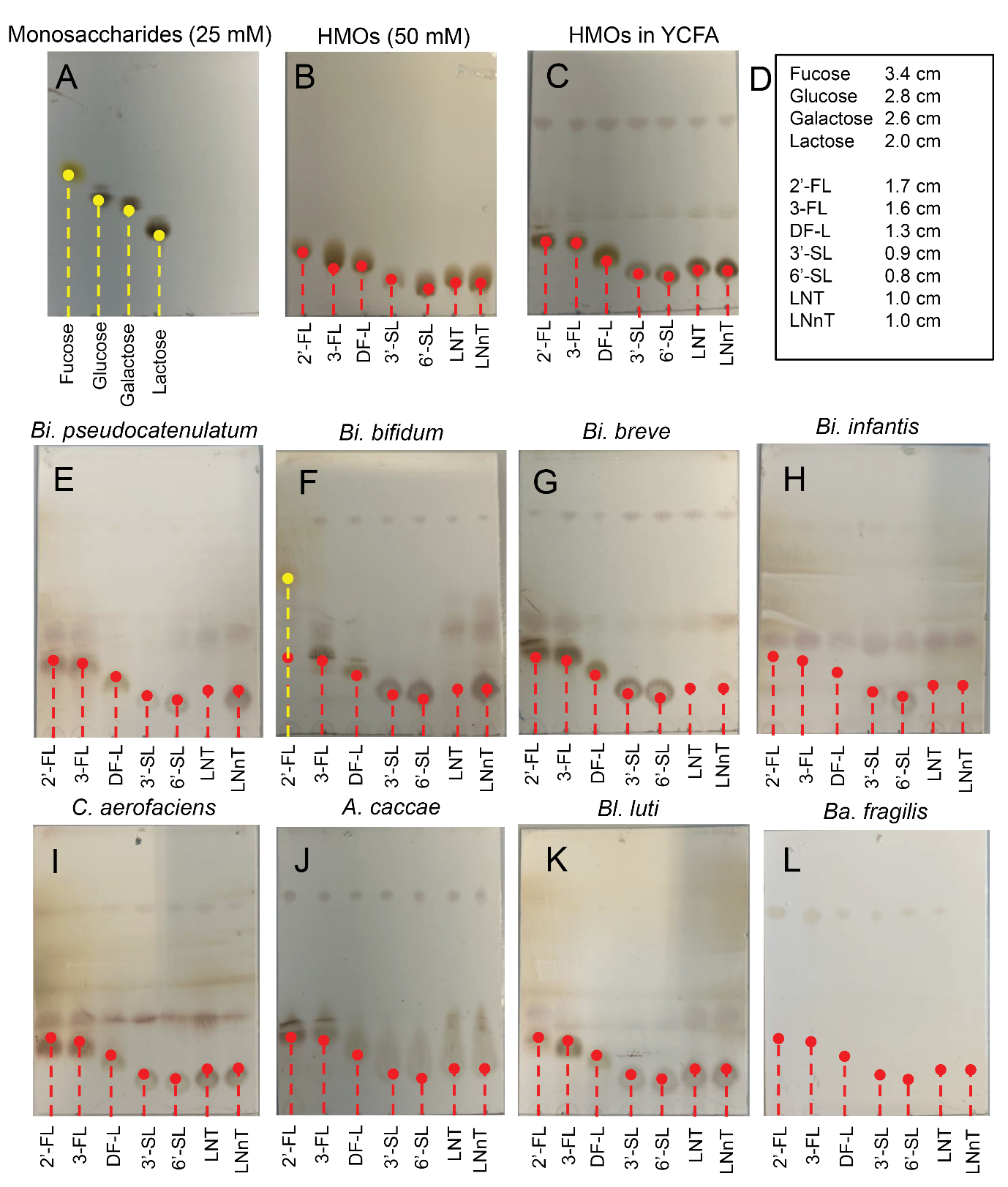

**Supplementary Figure 10.** **TLC analysis of monoculture fermentation**. (A) Monosaccharide standards, (B) HMO standards in water, (C) HMO standards in YCFA, with migration distances shown in (D). Spent media from monocultures of (E) B. pseudocatenulatum, (F) B. bifidum, (G) Bi. breve, (H) Bi. infantis, (I) C. aerofaciens, (J) A. caccae, (K) Bl. luti, and (L) Ba. fragilis.

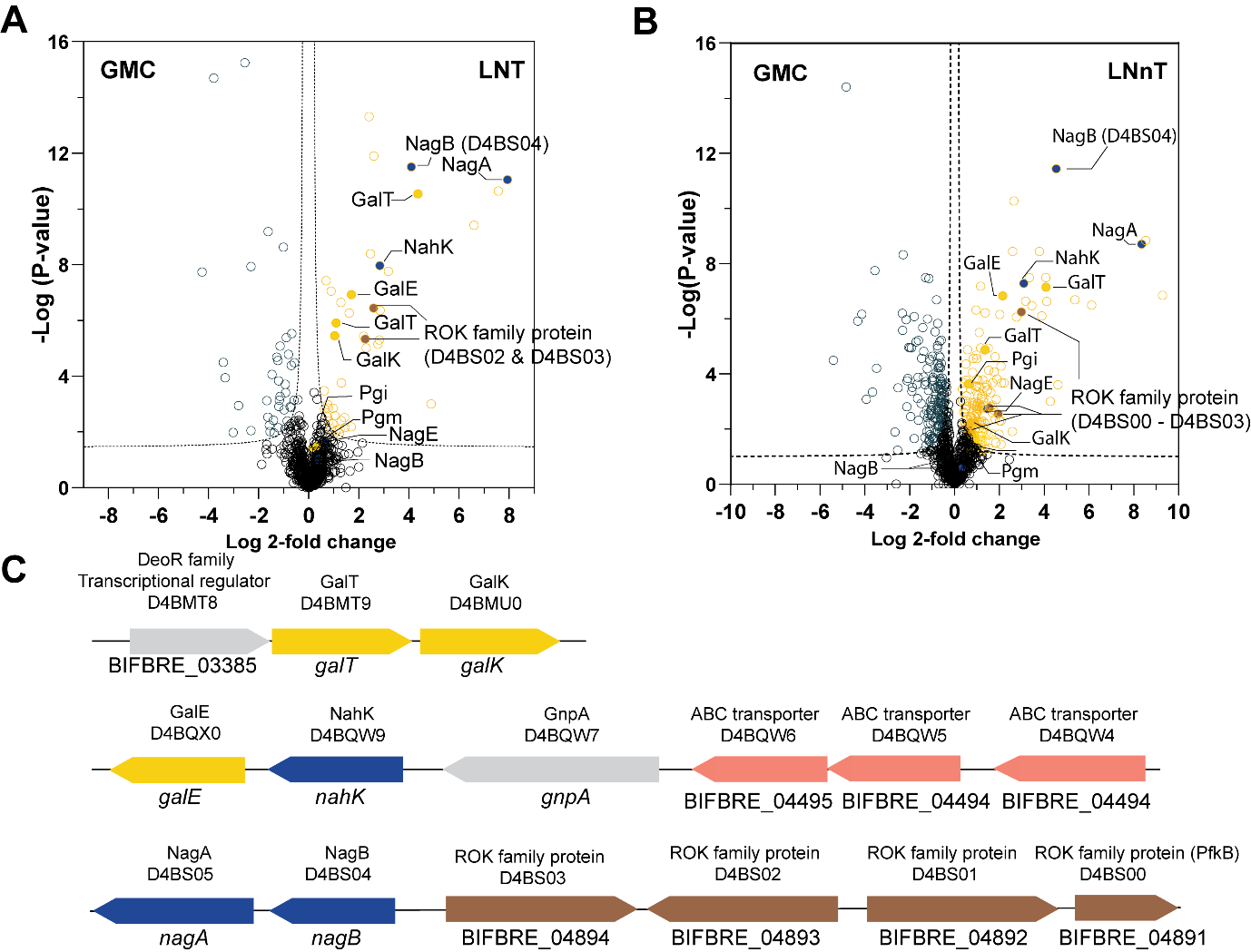
**Supplementary Figure 11.** **Proteins involved in LNT and LNnT metabolism in Bi. breve**. (A–B) Volcano plots of protein abundance in LNT (A, right) or LNnT (B, right) versus GMC (left). Each point represents one protein; data are from three independent experiments. Proteins enriched in LNT/LNnT are outlined in gold, those enriched in GMC in teal. Functional classes: Leloir pathway (yellow), GlcNAc metabolism (navy), ROK family (brown). (C) Genomic loci encoding upregulated proteins.

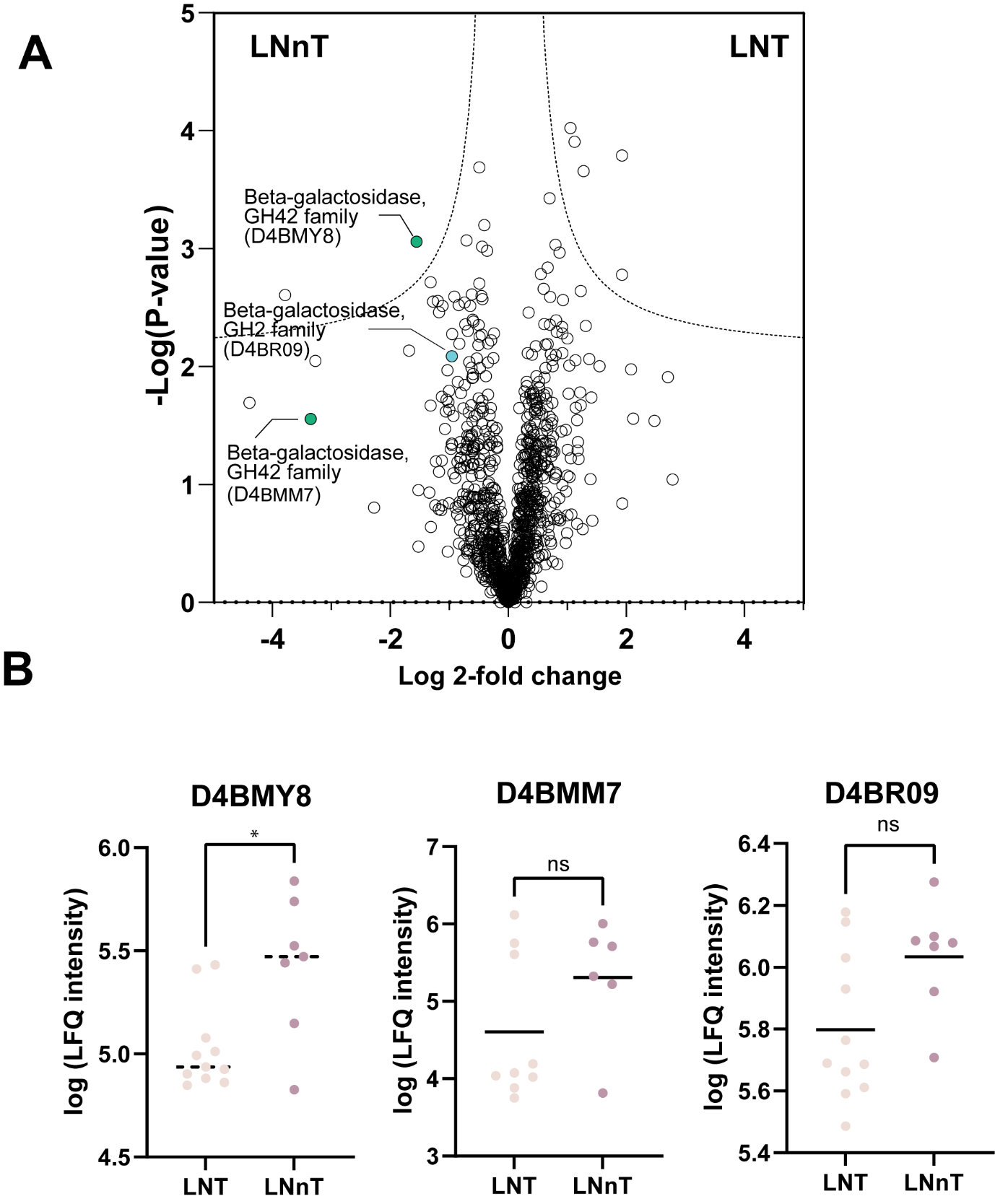
**Supplementary Figure 12.** **Differential protein expression of Bi. breve in LNT versus LNnT**. (A) Volcano plot comparing LNT and LNnT. (B) β-galactosidase LFQ intensities (log10-transformed) analyzed by Mann–Whitney test; *p < 0.05

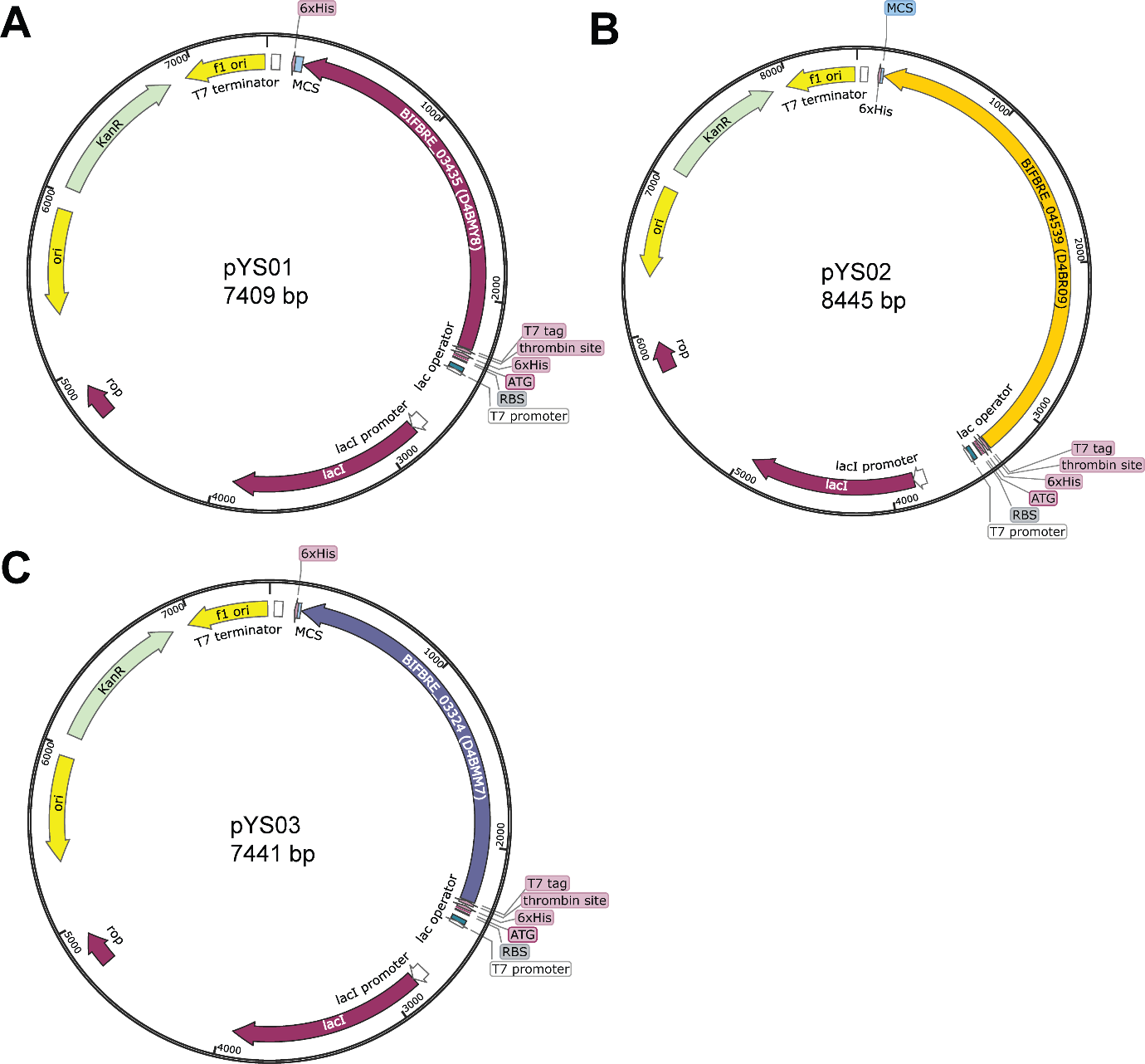
**Supplementary Figure 13. Plasmid maps for β-galactosidase expression constructs. (A) pYS01, (B) pYS02, (C) pYS03**

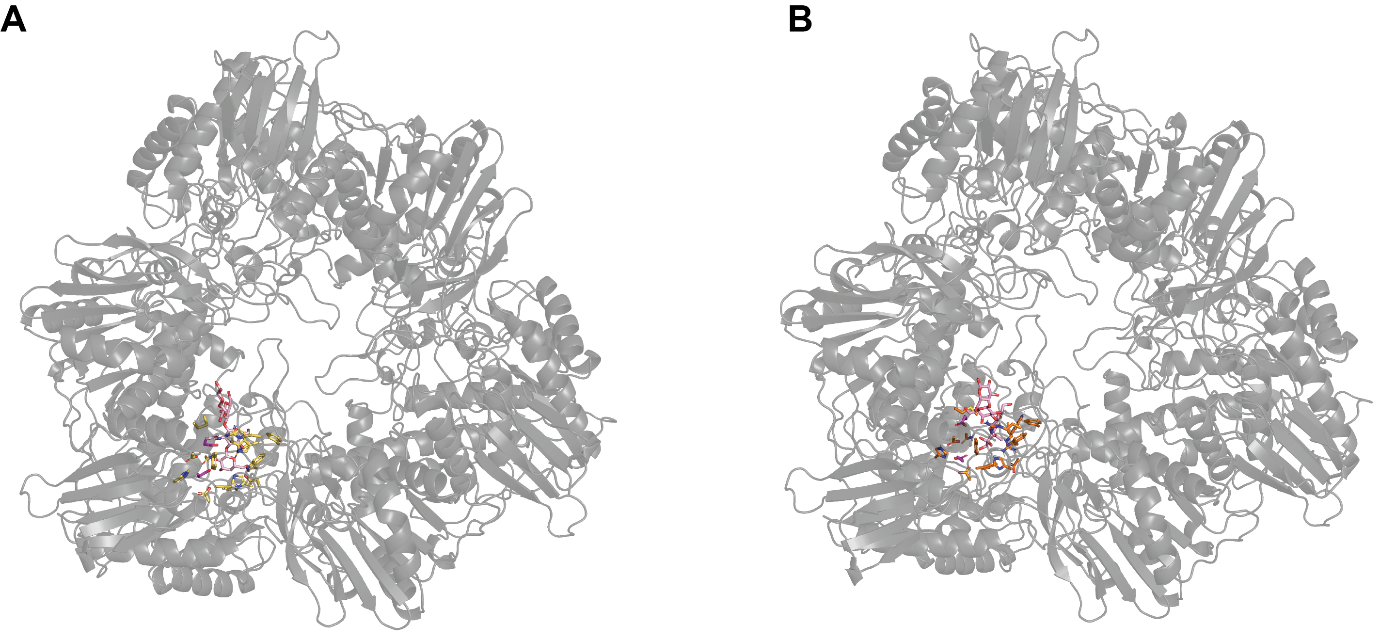

**Supplementary Figure 14.** **Trimeric structure of β-galactosidases and their catalytic pockets in complex with LNT/LNnT.** (A) structure of 8IBT, (B) structure of D4BMY8

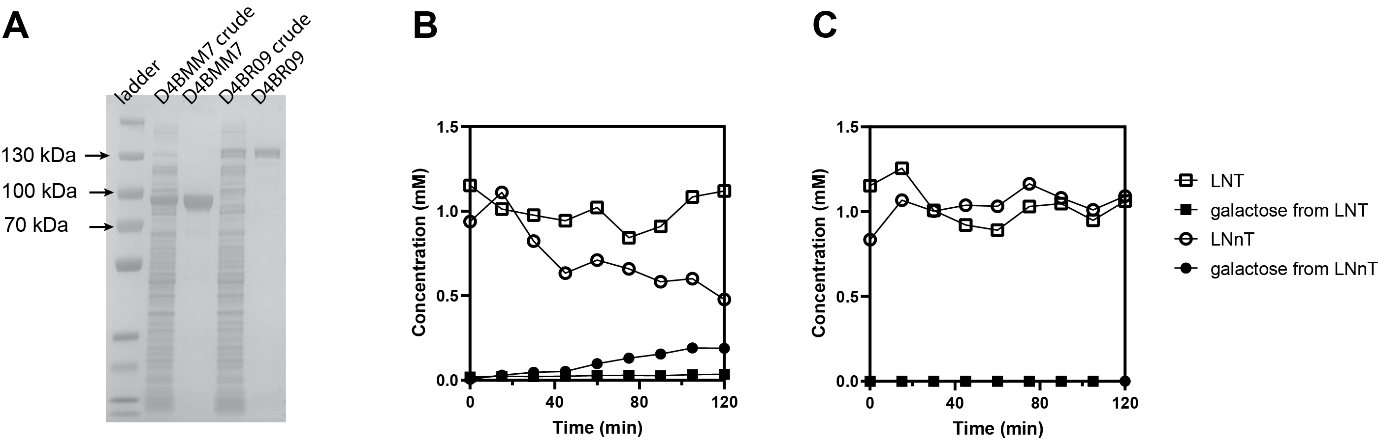

**Supplementary Figure 15.** **Profiling of recombinant D4BR09 and D4BMM7.** (A) SDS-PAGE of recombinant D4BR09 and D4BMM7. Hydrolysis of LNT and LNnT and galactose release by purified D4BR09 (B), and D4BMM7 (C).

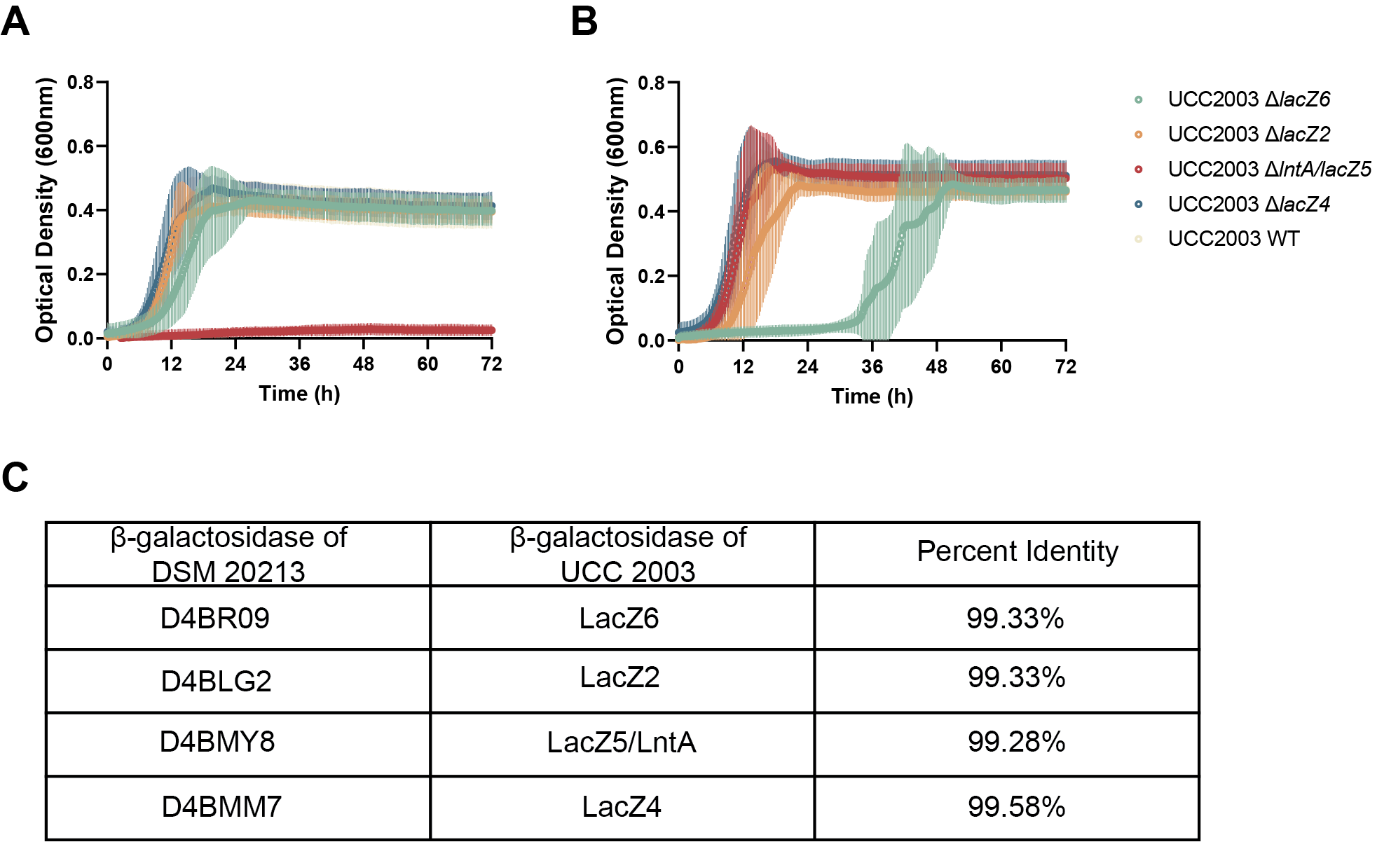
**Supplementary Figure 16. Growth of Bifidobacterium breve UCC 2003 wild-type and mutant strains and comparison of β-galactosidases.** (A,B) Growth curves of Bi. breve UCC2003 wild-type (WT) and knockout strains cultured in LNT (A) and LNnT (B). Data are presented as mean ± SD from three independent experiments. (C) Correspondence between β-galactosidases of Bi. breve DSM 20213 and their counterparts in UCC2003. Percent sequence identities were determined by amino acid alignment using BLASTP.
